# Plasma Membrane Phylloquinone Biosynthesis in Nonphotosynthetic Parasitic Plants

**DOI:** 10.1101/257519

**Authors:** Xi Gu, Ing-Gin Chen, Scott A. Harding, Batbayar Nyamdari, Maria A. Ortega, Kristen Clermont, James H. Westwood, Chung-Jui Tsai

## Abstract

Phylloquinone is a lipophilic naphthoquinone found predominantly in chloroplasts and best known for its function in photosystem I electron transport and disulfide bridge formation of photosystem II subunits. Phylloquinone has also been detected in plasma membrane preparations of heterotrophic tissues with potential transmembrane redox function, but the molecular basis for this noncanonical pathway is unknown. Here we provide evidence of plasma membrane phylloquinone biosynthesis in a nonphotosynthetic holoparasite *Phelipanche aegyptiaca*. A nonphotosynthetic and nonplastidial role for phylloquinone is supported by transcription of phylloquinone biosynthetic genes during seed germination and haustorium development, by plasma membrane-localization of alternative terminal enzymes, and by detection of phylloquinone in germinated seeds. Comparative gene network analysis with photosynthetically competent parasites revealed a bias of *Phelipanche* phylloquinone genes toward coexpression with oxidoreductases involved in plasma membrane electron transport. Genes encoding the plasma membrane phylloquinone pathway are also present in several photoautotrophic taxa of Asterids, suggesting an ancient origin of multifunctionality. Our findings suggest that nonphotosynthetic holoparasites exploit alternative targeting of phylloquinone for transmembrane redox signaling associated with parasitism.

## Introduction

Phylloquinone (vitamin K1) is a membrane-bound naphthoquinone derivative known to function as an essential electron acceptor in photosystem I (PSI) (Brettel et al., 1986). Phylloquinone also serves as an electron carrier for protein disulfide bond formation crucial for PSII assembly (Furt et al., 2010; Karamoko et al., 2011; Lu et al., 2013). Accordingly, phylloquinone is found predominantly in thylakoids, and most phylloquinone-deficient *Arabidopsis thaliana* mutants are seedling-lethal or growth-impaired (reviewed in Gilles et al., 2016). A sizable phylloquinone pool is stored in plastoglobules attached to thylakoid membranes (Lohmann et al., 2006; Eugeni Piller et al., 2012). A small portion of leaf phylloquinone is present as fully-reduced quinol, with potential redox function during senescence or dark growth (Oostende et al., 2008).

The eubacterial cognate menaquinone (vitamin K2) functions in respiratory electron transport across the cell membrane (Nowicka and Kruk, 2010). A similar role for phylloquinone in plant plasma membrane (PM) electron transport has also been proposed, and phylloquinone has been detected in PM preparations of maize roots (Lüthje et al., 1998; Lochner et al., 2003). UV-irradiation of cultured carrot cells destroyed phylloquinone, and blocked transmembrane electron transport until restoration by phylloquinone feeding (Barr et al., 1992). The PM redox activities can be inhibited by phylloquinone antagonists, dicumarol and warfarin, whereas applications of menadione (vitamin K3) or phylloquinone restored transmembrane electron flow (Döring et al., 1992a; Döring et al., 1992b; Lüthje et al., 1992). Despite these early reports, however, molecular evidence that directly supports PM-targeting of phylloquinone remains elusive.

The phylloquinone biosynthetic pathway has an endosymbiotic origin, comprising a series of “Men” proteins named after their eubacterial homologs (Figure 1). A notable exception is the penultimate enzyme (step 9, Figure 1A) of the flavin-containing NAD(P)H quinone oxidoreductase (FQR/NQR/QR) family recently shown to be necessary for phylloquinone biosynthesis in plants and cyanobacteria (Eugeni Piller et al., 2011; Fatihi et al., 2015). The corresponding enzyme in *A. thaliana*, type II NAD(P)H dehydrogenase C1 (NDC1), was first identified as a mitochondrial respiratory chain component (Michalecka et al., 2003), but later was also found to be targeted to the chloroplast (Carrie et al., 2008), with additional functions in plastoquinone reduction and redox cycling of α-tocopherols in plastoglobules (Eugeni Piller et al., 2011). Intracellular compartmentalization is a hallmark of phylloquinone biosynthesis, with the early (steps 1-4, Figure 1A) and late pathway steps (8-10) occurring in chloroplasts, and the intermediate steps (5-7) in peroxisomes (Babujee et al., 2010). Even within the chloroplast, the three terminal steps shuttle between envelope membranes (MenA, Schultz et al., 1981), plastoglobules (NDC1, Eugeni Piller et al., 2011), and thylakoid membranes (MenG, Kaiping et al., 1984), highlighting the complex trafficking involved in this pathway.

**Figure 1.**
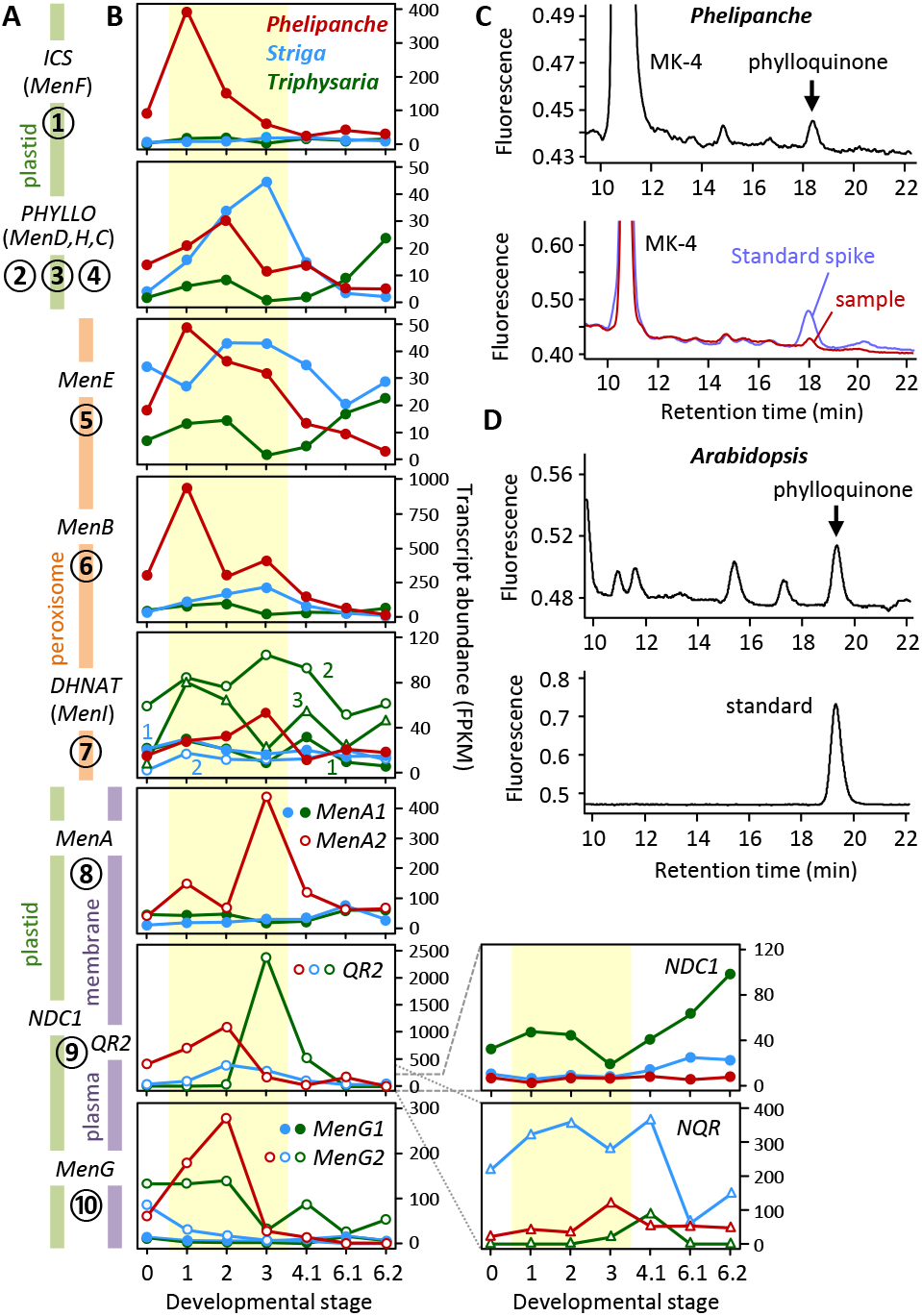
Phylloquinone biosynthesis in parasitic plants. (A) A simplified phylloquinone biosynthetic pathway from top to bottom as numbered and color-coded by their predicted subcellular localization. (B) Expression profiles of phylloquinone genes during parasitic plant development. Gene order is the same as in (A), and Y-axis is shown on the right. Developmental stages are as reported (Yang et al., 2015): 0, imbibed seeds; 1, germinated seeds/radicles; 2, HIF-treated seedlings; 3, haustoria, pre-vascular connection; 4.1, haustoria, post-vascular connection; 6.1, leaves/stems; 6.2, floral buds. Data from alternative gene candidates for the penultimate step are shown on the lower right panels, with grey dashed/dotted lines denoting the scale. (C) HPLC detection of phylloquinone in germinated *P. aegyptiaca* seeds (top panel), and with spiked authentic standard (lower panel). Menaquinone (MK-4) was included as a reference. (D) HPLC detection of phylloquinone in *A. thaliana* seed (top panel), with authentic standard (lower panel).

We have observed in multiple photosynthetic taxa that phylloquinone-specific genes such as *MenA* and *MenG* have measurable expression in heterotrophic tissues where photosynthetic genes are barely detected. To ascertain a noncanonical phylloquinone pathway, we exploited parasitic plants as a photosynthesis-free study system. Among angiosperm parasite families, only the Orobanchaceae contains species that span the full spectrum of photosynthetic capacities, and for which rich transcriptomic resources are available (Westwood et al., 2010; Yang et al., 2015). Of particular interest are obligate holoparasites, such as *Phelipanche aegyptiaca*, that are devoid of photosynthetic activity and obtain all of their carbon from their hosts. In contrast, obligate (*e.g.*, *Striga hermonthica*) and facultative (*e.g.*, *Triphysaria versicolor*) hemiparasites are partially or fully photosynthetic. Here, we report on the biosynthesis and PM targeting of phylloquinone in the nonphotosynthetic *Phelipanche aegyptiaca*. Gene network analysis revealed a strong link between phylloquinone and cellular oxidation-reduction processes implicated in parasitic invasion and haustorium development. We propose that parasitic plants exploit alternative phylloquinone targeting for PM redox regulation associated with parasitism.

## Results

### *P. aegyptiaca* contains the full complement of phylloquinone biosynthetic genes

Phylloquinone pathway protein sequences of *Mimulus guttatus*, a photoautotroph from Phrymaceae sister to Orobanchaceae, were searched against transcript assemblies available from the Parasitic Plant Genome Project (Yang et al., 2015). Full-length coding sequences were identified for *PaICS* and *PaMenE* genes in *P. aegyptiaca*, along with partial assemblies of other phylloquinone pathway genes. Fragmented transcripts may represent nonfunctional relics of genes undergoing degeneration or may reflect technical limitations of *de novo* assembly that prevented identification of the full complement of phylloquinone biosynthetic genes. The recovery of full-length *PaICS* and *PaMenE* transcripts strongly favored the latter scenario.

We addressed the fragmented assembly challenge by developing a pipeline called PLAS (Parallelized Local Assembly of Sequences) that combines reference-guided mapping (against the *Mimulus* proteome in this case) with iterative *de novo* assembly for transcriptome reconstruction. When applied to the RNA-seq datasets of parasitic plants, we successfully recovered full-length transcripts with intact ORFs for all known phylloquinone genes from the holoparasite (Supplemental Dataset 1). These transcripts were detected at moderate to high levels during *P. aegyptiaca* development, except *PaNDC1*—the latest addition to the pathway—which was poorly expressed (FPKM<10) throughout the holoparasite (Figure 1B). *PHYLLO*, *MenE* and *DHNAT* transcripts were detected at similar levels in the three parasites. In contrast, the expression patterns of *ICS*, *MenB*, *MenA* and *MenG* differed between *P. aegyptiaca* and its photosynthetic relatives *S. hermonthica* and *T. versicolor*, especially in response to germination stimulants and haustorium-inducing factors (HIFs) during early development (Stages 1-3, Figure 1B). However, the apparent lack of *PaNDC1* expression weakened support for a functioning phylloquinone pathway. Alternatively, retention and expression of the other seven phylloquinone pathway genes may point to a different pathway configuration at the penultimate step in the holoparasite.

We therefore sought to substantiate phylloquinone production in *P. aegyptiaca*, focusing on early development prior to host exposure and haustorium initiation to avoid potential contamination from the photosynthetic host. HPLC analysis confirmed the presence of phylloquinone in germinated *P. aegyptiaca* seeds (Figure 1C), with an estimated level of 0.05±0.02 pmol/mg dry weight (n=3). For reference, phylloquinone levels in *A. thaliana* seeds were 0.12±0.03 pmol/mg dry weight (n=3) (Figure 1D). The result lent support to phylloquinone biosynthesis in the holoparasite. We next examined other candidates that could participate in the reduction of demethylphylloquinone at the penultimate step (step 9). Besides the multifunctional NDC1 (Fatihi et al., 2015), menadione-reducing activities have been demonstrated for two other groups of evolutionarily conserved QR, one represented by *T. versicolor* TvQR2 (Wrobel et al., 2002) and *A. thaliana* AtFQR1 (Laskowski et al., 2002) of the type IV family (Patridge and Ferry, 2006), and the other by soybean (*Glycine max*) GmNQR of the DT-diaphorase type (Schopfer et al., 2008). *PaNQR* transcripts were detected at moderate levels throughout *P. aegyptiaca* development (Figure 1B), but they were not coexpressed with any phylloquinone pathway genes (see below). By contrast, *PaQR2* was well expressed during early stages of *P. aegyptiaca* development like other phylloquinone genes (Figure 1B). Type IV QR2 has in fact been implicated in two-electron reduction of menaquinone and phylloquinone to their respective quinols in the manually curated KEGG reference pathways (Kanehisa et al., 2018). The structural similarity between phylloquinone and its demethylated precursor, and the reported substrate promiscuity of the orthologous TvQR2 (Wrobel et al., 2002) lent further support to PaQR2 involvement in the penultimate step of *P. aegyptiaca* phylloquinone biosynthesis.

### Phylloquinone biosynthesis is redirected to the plasma membrane in the holoparasite

Identification of the phylloquinone pathway in *P. aegyptiaca* with vestigial plastids raised the possibility that the biosynthetic enzymes exhibit alternative targeting with nonplastidial function(s). Protein subcellular localization analyses predicted plastid- and peroxisome-targeting of early (PaICS and PaPHYLLO) and intermediate (PaMenE, PaMenB and PaDHNAT) pathway steps, respectively, similar to their orthologs in photoautotrophic taxa (Supplemental Table S1 and Supplemental Figures S1-S3). However, the predicted polypeptides of late pathway steps PaMenA and PaMenG are truncated at the N-terminus relative to their photoautotrophic orthologs (Figure 2) and scored poorly for plastid-targeting with multiple prediction programs (Figure 2A, Supplemental Figures S4 and S5). The N-truncation of PLAS-derived *PaMenA* and *PaMenG* transcripts was independently confirmed by RT-PCR cloning and sequencing (GenBank accession numbers MT506520 and MT506521), and by identification of N-truncated orthologs from other species (see below).

**Figure 2.**
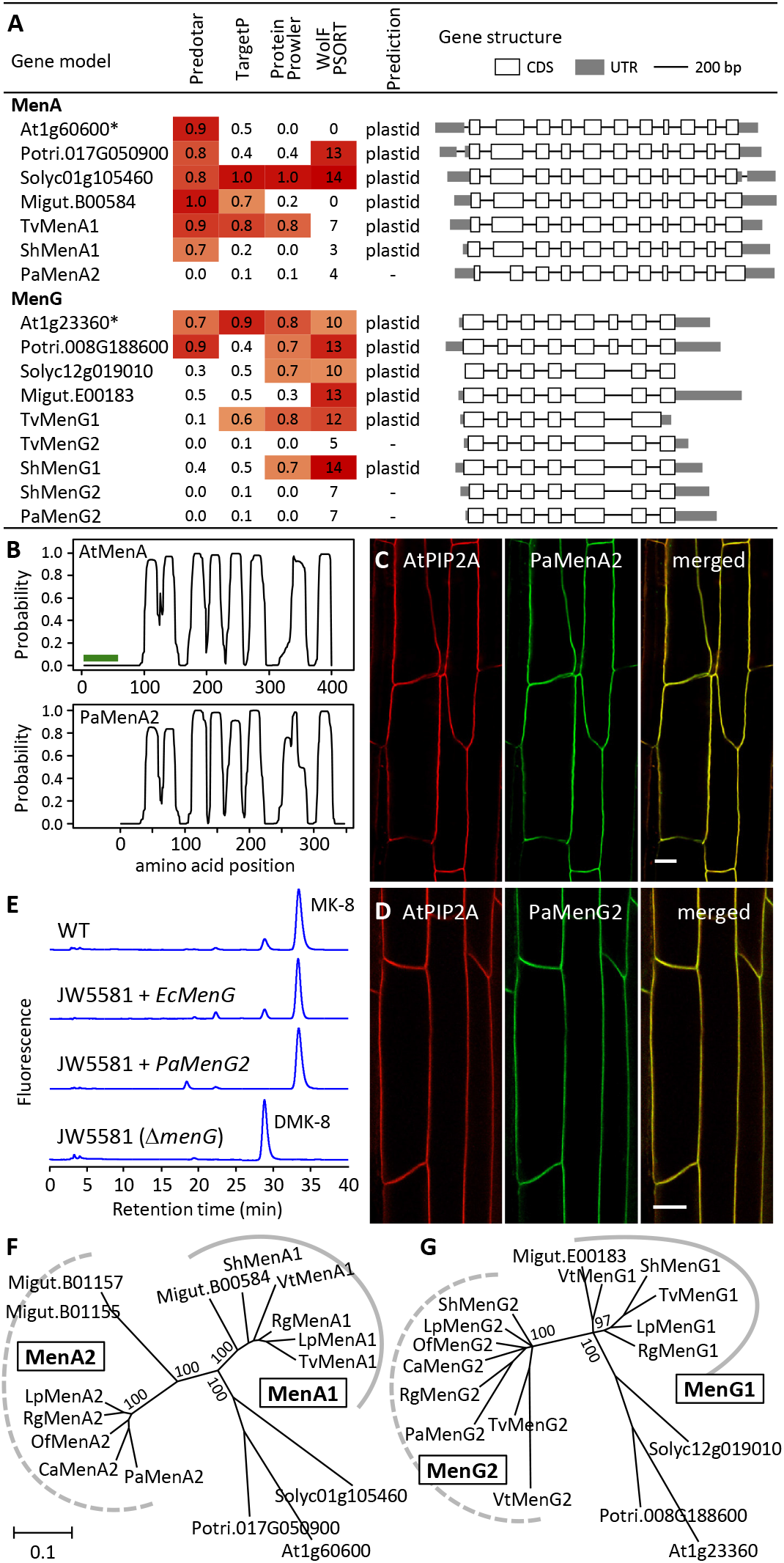
Characterization of MenA and MenG. (A) Plastid-targeting prediction of MenA and MenG polypeptides from various species. Heatmaps show prediction strengths above the 50^th^ percentile of each method. Those with a high prediction score from at least one program are deemed potentially plastidial, as not all methods accurately predict experimentally characterized plastidial proteins (asterisks). Introns are not drawn to scale. (B) Transmembrane domain prediction of AtMenA (green line denoting the transit peptide) and PaMenA2 (shifted x-axis for domain alignment). (C-D) Confocal images of PaMenA2-GFP (C) and PaMenG2-GFP (D) colocalization with a plasma membrane marker AtPIP2A-mCherry. Scale bars = 20 μm. (E) HPLC analysis of *E. coli* ΔmenG mutant strain JW5581 expressing *PaMenG2* or the *EcMenG* control. (D)MK-8, (demethyl)menaquinone-8. (F-G) Bayesian phylogeny of MenA and MenG from representative Asterids and two Rosids. At, *Arabidopsis thaliana*; Ca, *Conopholis americana*; Lp, *Lindenbergia philippensis*; Migut, *Mimulus guttatus*; Of, *Orobanche fasciculata*; Pa, *Phelipanche aegyptiaca*; Potri, *Populus trichocarpa*; Rg, *Rehmannia glutinosa*; Solyc, *Solanum lycopersicum*; Sh, *Striga hermonthica*; Tv, *Triphysaria versicolor*; Vt, *Verbascum thapsus*. Branch support is shown for major nodes.

PaMenA is predicted as an integral protein with eight transmembrane helices in a topology similar to that of AtMenA (Figure 2B). The absence of an N-terminal plastid-targeting peptide in PaMenA suggests its localization to other cellular membranes. The penultimate PaQR2, like its photoautotrophic orthologs, also lacks plastid-targeting sequence (Supplemental Figures S6), which contrasts with the alternative NDC1s that are predicted plastidial (Supplemental Table S1). Indeed, PM-association has been reported for *A. thaliana*, rice (*Oryza sativa*) and yeast (*Candida albicans*) QR2 orthologs (Marmagne et al., 2004; Natera et al., 2008; Li et al., 2015). These data suggest the post-peroxisomal steps are likely targeted to PM in *P. aegyptiaca*. We generated stably transformed *Nicotiana benthamiana* expressing *35S:PaMenA-GFP* or *35S:PaMenG-GFP* along with a PM marker *35S:AtPIP2A-mCherry* (Nelson et al., 2007). Confocal imaging showed co-localization of PaMenA and PaMenG with the PM marker in transgenic roots (Figure 2C-D), providing experimental support for alternative targeting of the N-truncated PaMenA and PaMenG in *P. aegyptiaca*.

### The plasma membrane pathway has canonical activity and an ancient origin

The absence of the N-terminal transit signal is not expected to impact (mature) enzyme catalysis. Using PaMenG as a test case, we performed complementation experiments with the *E. coli* Δ*menG* mutant strain JW5581 (Baba et al., 2006). The *E. coli* MenG, also called UbiE, is a dual C-methyltransferase involved in both menaquinone and ubiquinone biosynthesis, and its mutation leads to over-accumulation of demethylmenaquinone (Lee et al., 1997). Constitutive expression of *PaMenG* in the mutant strain restored menaquinone production similar to the *E. coli EcMenG*-complemented control (Figure 2E). The data provide biochemical evidence for canonical activity of the N-truncated PaMenG.

Interestingly, distinct *MenG1* and *MenG2* genes encoding long (plastidial) and short (PM) isoforms, respectively, are present in both *S. hermonthica* and *T. versicolor* (Figure 2A), suggesting the PM phylloquinone pathway may have evolved before the transition to parasitism. To strengthen this finding, we mined the One Thousand Plant Transcriptomes (1KP) database (Leebens-Mack et al., 2019) and identified several photoautotrophic species—all from Lamiales—that harbor both long and short isoforms of MenA and/or MenG, including *Lindenbergia philippensis*, *Rehmannia glutinosa* and *Verbascum thapsus*. In contrast, only N-truncated short isoforms were found in two other Orobanchaceae holoparasites, *Orobanche fasciculata* and *Conopholis americana* (Supplemental Figures S4-S5). Phylogenetic analysis clustered the plastidial and PM isoforms into distinct groups for both MenA and MenG (Figure 2F-G), suggesting their origin from lineage-specific duplication events (hereafter, the PM-localized short isoforms are referred to as MenA2 and MenG2). Of note, transcripts encoding PM-targeted MenG2 were detected at higher levels than those encoding plastidial MenG1 in both *S. hermonthica* and *T. versicolor* (Figure 1B), suggesting divergent regulation of the two MenG isoforms in photosynthetic hemiparasites.

### Photosynthetic and nonphotosynthetic parasites exhibit distinct phylloquinone gene coexpression patterns

We detected high levels of coexpression among phylloquinone genes in the holoparasites, reminiscent of the patterns observed in photosynthetic taxa, including *T. versicolor* (Figure 3) (except for *A. thaliana AtICSs* and *AtDHNATs* involved in salicylic acid biosynthesis (Garcion et al., 2008) and peroxisomal β-oxidation (Cassin-Ross and Hu, 2014), respectively). Interestingly, early- and late-pathway genes showed distinct coexpression patterns in the obligate hemiparasite *S. hermonthica*, perhaps indicative of dual involvement of those genes in plastidial and nonplastidial functions. To shed light on PM-phylloquinone functions, we extracted the top 500 most highly-correlated transcripts for each phylloquinone gene except the alternative *QR2* and *NDC1.* The union set contained 2447, 3677 and 3930 unique transcripts for *P. aegyptiaca*, *S. hermonthica* and *T. versicolor*, respectively. Interestingly, *QR2* but not *NDC1* was captured in all three networks. The smaller *P. aegyptiaca* gene set is consistent with stronger coexpression of phylloquinone genes when compared to *S. hermonthica* and *T. versicolor* (Figure 3A-C). Subsets of Gene Ontology (GO)-annotated (Biological Process) transcripts (645, 1199 and 1173 for *P. aegyptiaca*, *S. hermonthica* and *T. versicolor*, respectively) were then subjected to functional enrichment analysis. Transcripts associated with ‘photosynthesis’ comprised 3-4% of the GO-annotated transcripts in photosynthetic parasites but were negligible in *P. aegyptiaca* (Figure 3G). In contrast, transcripts associated with ‘oxidation-reduction process’, ‘protein phosphorylation’ and ‘defense response’ were more enriched in *P. aegyptiaca* relative to *S. hermonthica* and *T. versicolor* (Figure 3G).

**Figure 3.**
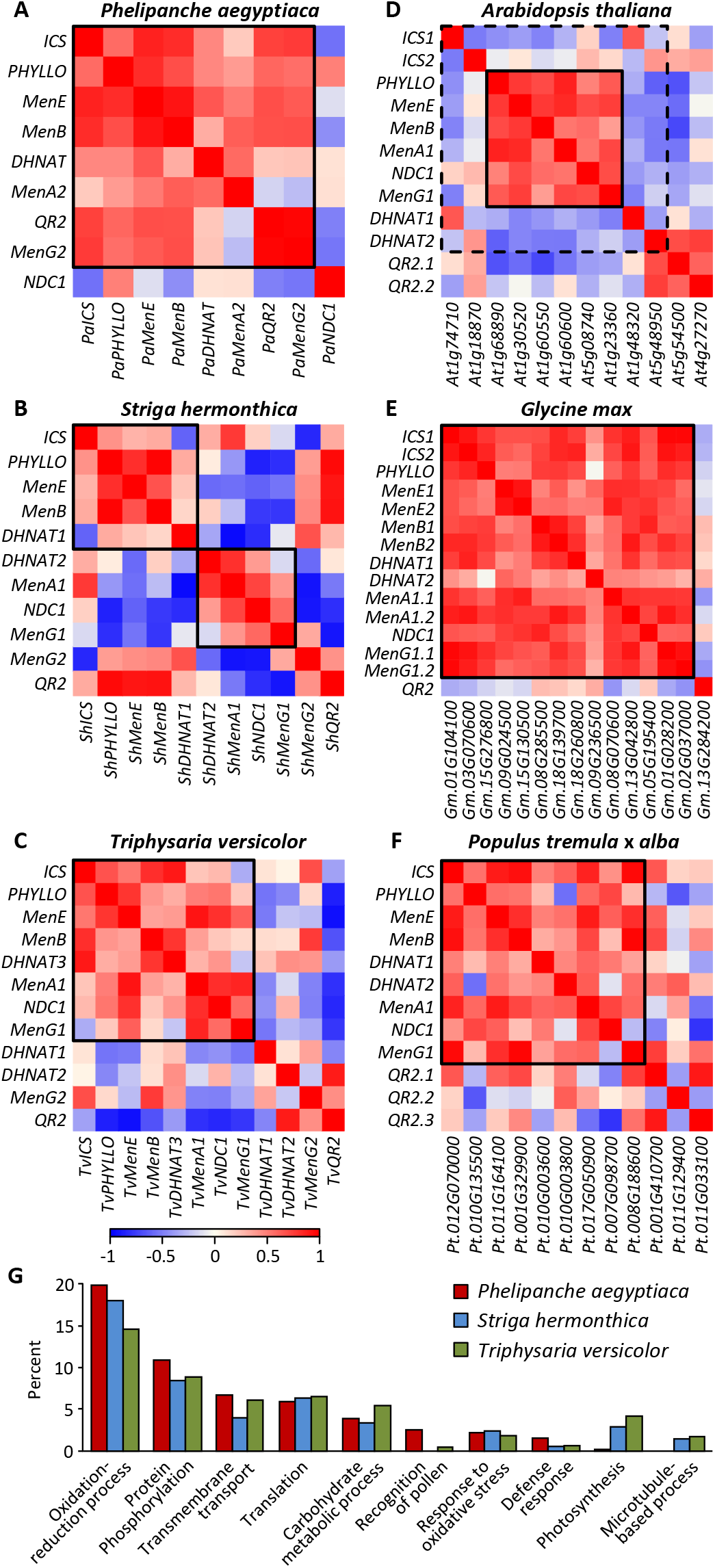
Coexpression of phylloquinone genes. (A-F) Coexpression patterns among phylloquinone biosynthetic genes, including multifunctional *NDC1* and *QR2*, based on Gini correlation coefficient. Relevant genes or gene members involved in phylloquinone biosynthesis are boxed. The exceptions are *A. thaliana AtICSs* involved in salicylic acid biosynthesis for defense and *AtDHNATs* in peroxisomal β-oxidation (dashed box), besides phylloquinone biosynthesis. The corresponding *Arabidopsis*, *Glycine* and *Populus* gene models are shown on the x-axis with shortened prefix for soybean (Gm = Glyma) and poplar (Pt = Potri). (G) GO enrichments of phylloquinone-coexpressed genes defined as the union set of the top 500 most highly-correlated transcripts for each phylloquinone gene. Only the top 10 categories are shown. Similar results were obtained using Gini correlation coefficient ≥0.8 to extract phylloquinone-coexpressed genes.

We focused on transcripts assigned to oxidation-reduction, defense and photosynthesis GO terms for coexpression network analysis. Inclusion of orthologs from all three parasites resulted in 359, 544 and 559 non-redundant transcripts from *P. aegyptiaca*, *S. hermonthica* and *T. versicolor*, respectively (Supplemental Dataset 2). Network visualization of coexpression patterns revealed striking differences (Figure 4). Two dense modules were detected for photosynthetic *S. hermonthica* and *T. versicolor*; one enriched with photosynthesis genes (Figure 4A, green nodes) and the other containing known parasitism genes (blue, magenta and cyan nodes, see below). However, only one dense module containing parasitism genes was detected for the holoparasite. The phylloquinone genes (orange nodes) were highly interconnected with parasitism genes in the *P. aegyptiaca* network but were scattered over the two modules in *S. hermonthica* and *T. versicolor* networks, presumably reflecting dual functionality in these taxa. We ranked genes by the number of edges they shared with phylloquinone genes (referred to as *k*_PhQ_) in each network and observed a strong enrichment of phylloquinone-interconnected genes in the smaller *P. aegyptiaca* network. More than 23% of *P. aegyptiaca* nodes had a *k*_PhQ_ =4-6 (*i.e.*, connected with a majority of the seven phylloquinone genes). However, less than 3% of the *S. hermonthica* and *T. versicolor* nodes met the same criterion (*k*_PhQ_ ≥5 of 9-10 phylloquinone genes), and only 10 and 15% of their respective nodes had a *k*_PhQ_ ≥4 (Figure 4A, *k*_PhQ_ bars).

**Figure 4.**
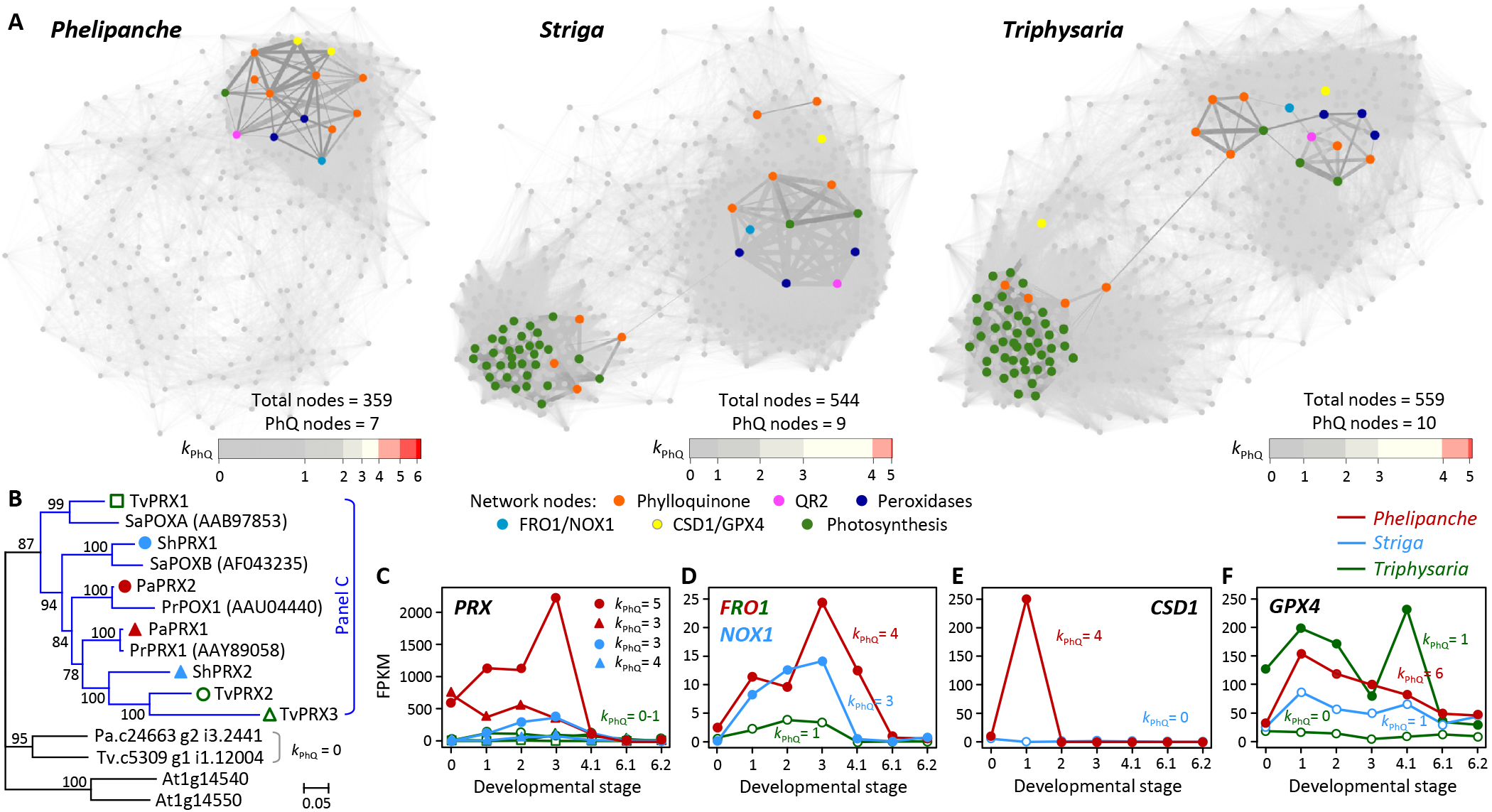
Phylloquinone gene coexpression networks. (A) Network visualization of the three parasitic plants. Edge thickness reflects the coexpression strength. Key gene nodes are color-coded by pathway or family, and *QR2* is colored differently from the other phylloquinone genes (*NDC1* was not captured in any of the networks). Horizontal bars depict the distribution of nodes according to their connectivity with phylloquinone genes (*k*_PhQ_). (B) Bayesian phylogeny of peroxidases (PRX). Orthologs of experimentally characterized PRXs are color-coded by species in the blue clade. (C-F) Expression profiles of *PRX* (C), *FRO1/NOX1* (D), *CSD1* (E), and *GPX4* (F) orthologs. Solid symbols denote phylloquinone-coexpressed genes (*k*_PhQ_ ≥3), and others are shown in open symbols. Developmental stages are the same as in Figure 1.

### *P. aegyptiaca* phylloquinone gene networks associate parasitism with plasma membrane redox signaling

Several oxidoreductases known to be involved in parasitism were captured in the phylloquinone gene network of *P. aegyptiaca*. Root-specific apoplastic/secretory peroxidases (PRXs or POXs), including *S. asiatica* SaPOXA and SaPOXB and *P. ramosa* PrPOX1 and PrPRX1, have established roles in haustorium induction via generation of reactive oxygen species (ROS) and oxidation of host cell wall-derived phenolics as HIFs (Kim et al., 1998; González-Verdejo et al., 2006; Veronesi et al., 2007). The *PRX* orthologs (Figure 4B, blue clade) were abundantly expressed in *P. aegyptiaca* throughout seed germination and haustorium development (Figure 4C). The phylloquinone-coexpression strength (*k*_PhQ_) was highest in *P. aegyptiaca*, followed by *S. hermonthica*, but was not observed for *T. versicolor PRXs*, or for homologs in the neighboring clade of the phylogenetic tree (Figure 4B-C). Thus, the phylloquinone-coexpression strength of the secretory PRXs appears to be positively correlated with parasitism.

Also implicated in parasitism are QR1 and QR2, depending on the species. QR1 is a chloroplast envelope protein of the ζ-crystalline type NAD(P)H oxidoreductase family involved in haustorium development of *T. versicolor* (Bandaranayake et al., 2010). However, haustorium formation in *Phtheirospermum japonicum* involves QR2 (Ishida et al., 2017). Accordingly, *QR2* but not *QR1* is responsive to HIFs in *S. asiatica* and *Phtheirospermum japonicum* (Ishida et al., 2016; Liang et al., 2016), similar to the patterns observed for *P. aegyptiaca* and *S. hermonthica* (Figure 1B). *QR1* was not coexpressed with phylloquinone genes, whereas the phylloquinone-coexpression strength of *QR2* is positively correlated with parasitism like the secretory PRXs. These results support a link between phylloquinone biosynthesis and parasitism.

We next searched the *P. aegyptiaca* phylloquinone subnetwork for redox proteins known to participate in transmembrane electron transport (Bérczi and Møller, 2000; Keyes et al., 2000). Several flavocytochrome *b* superfamily proteins are known to catalyze PM electron transport from cytosolic NAD(P)H to apoplastic acceptors, including NAD(P)H oxidases (NOX, also known as respiratory burst oxidase homologs) and ferric reductase oxidases (FRO) (Bérczi and Møller, 2000; Jain et al., 2014). The *P. aegyptiaca* transcriptome lacked *NOX*, but contained an *FRO* (*PaFRO1*) orthologous to *AtFRO4/AtFRO5* tandem duplicates that function as Cu-specific reductases at the root surface (Bernal et al., 2012). *PaFRO1* transcript levels were highest in prehaustorial and haustorial tissues (Figure 4D), consistent with potential involvement in the PM redox system. Interestingly, *S. hermonthica* harbors a *NOX*, but not *FRO*, perhaps indicative of lineage-specific transcriptome rewiring. Its counterpart in *S. asiatica* (*SaNOX1*) is indeed root-specific and HIF-responsive (Liang et al., 2016).

Electron transport chains are major sources of ROS which are tightly controlled by antioxidant systems, such as Cu/Zn superoxide dismutases (Cu/Zn-SOD or CDS) and glutathione peroxidases (GPX) (Mittler et al., 2004; Margis et al., 2008). *P. aegyptiaca PaCSD1* (*k*_PhQ_ =4) and *PaGPX4* (*k*_PhQ_ =6) were highly coexpressed with phylloquinone genes (Figure 4E-F). CSD1 is a known leaderless secretory protein frequently detected in apoplastic/secreted proteomes (Cheng et al., 2009; Pechanova et al., 2010), while GPX4 orthologs in *A. thaliana* (*AtGPX4/5* genome duplicates) were recently shown to be PM-anchored based on redox-sensitive GFP fusions (Attacha et al., 2017). Thus, besides the secretory PaPRXs, PaCSD1 and PaGPX4 may also participate in apoplastic redox modulation in the holoparasite. By contrast, the phylloquinone genes of photosynthetic parasites were coexpressed with gene family members specifically targeted to plastids (orthologs of AtCSD2 and/or Fe-SOD AtFSD2/3), mitochondria (AtGPX6) or cytosol (AtGPX8; Supplemental Dataset 2). The differential coordination between *P. aegyptiaca* and its photosynthetic relatives with distinct subcellular antioxidant systems is consistent with their contrasting photosynthetic capabilities, and with a prominent role of phylloquinone in PM redox homeostasis of the nonphotosynthetic holoparasite.

## Discussion

### An ancient origin of flexible phylloquinone biosynthesis and targeting

We present multiple lines of transcriptional, protein subcellular localization, enzymatic, and metabolic evidence to support extraplastidial phylloquinone biosynthesis in the nonphotosynthetic *P. aegyptiaca*. Our data further revealed the post-peroxisomal steps as key to flexible phylloquinone biosynthesis (Figure 5). Alternative targeting to the PM is facilitated by N-truncated MenA2 and MenG2 in conjunction with QR2. QR2 has previously been shown to function in mitigating oxidative stress in bacteria, yeast and plants (Laskowski et al., 2002; Wrobel et al., 2002; Patridge and Ferry, 2006; Li et al., 2015). In plants, *QR2s* exhibit root-biased expression and are highly responsive to auxins and quinones, including HIFs in the case of parasitic plants (Matvienko et al., 2001; Laskowski et al., 2002; Ishida et al., 2017) (Figure 1B). The proposed QR2 involvement in PM phylloquinone biosynthesis thus represents another example of a multi-functional NAD(P)H oxidoreductase for the penultimate step, analogous to NDC1 in plastidial phylloquinone biosynthesis (Fatihi et al., 2015).

**Figure 5.**
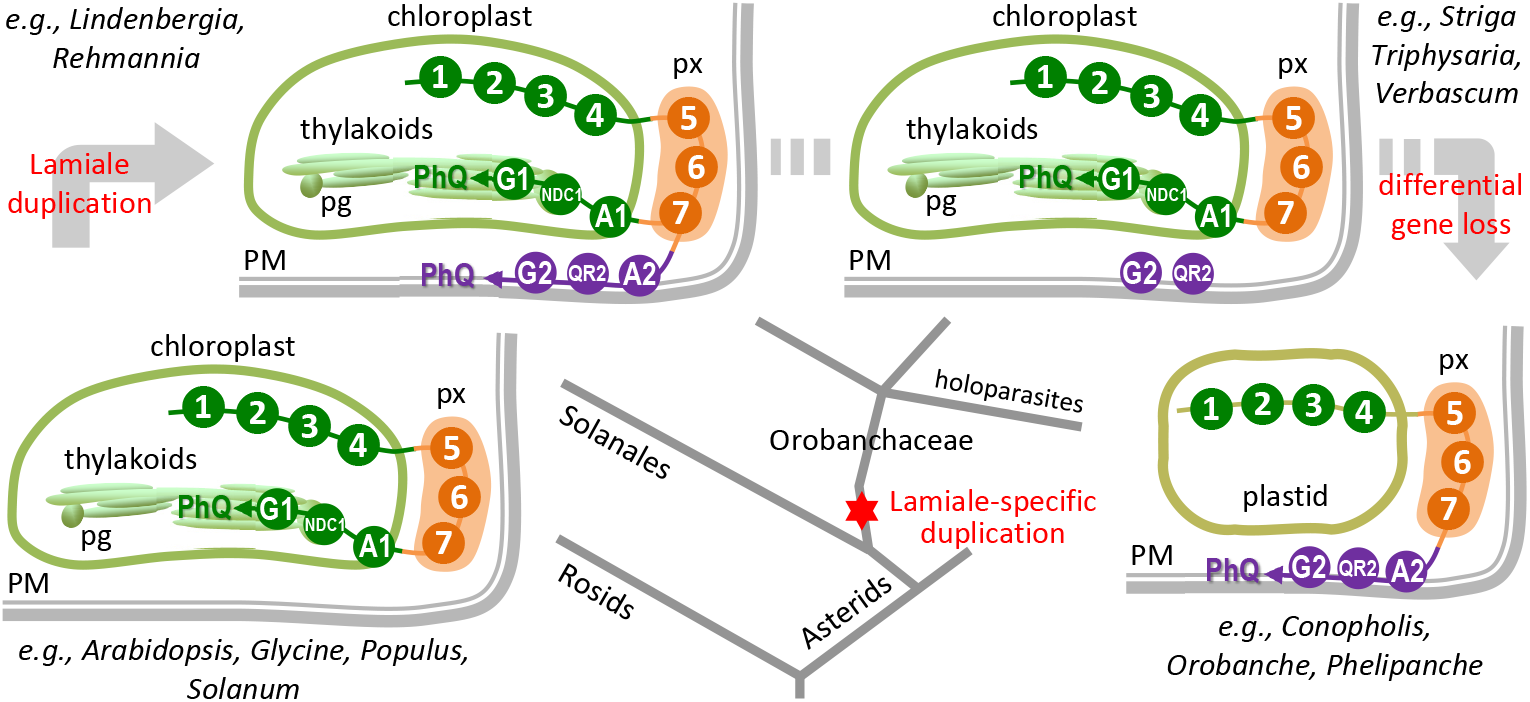
Evolutionary changes in subcellular localization of phylloquinone biosynthesis. Conserved early-and mid-pathway steps are shown as circled numbers (1-7, see Figure 1) and late pathway steps MenA and MenG are abbreviated as A1/A2 and G1/G2, respectively. Circle colors denote different subcellular compartments: green, plastid; orange, peroxisome (px); purple, plasma membrane (PM). Branches of the simplified eudicot phylogeny in the middle point to corresponding illustrations of changing pathway organization with representative species indicated. Clockwise from lower left, exclusively plastidial late steps in rosids and some asterids such as solanales; top left, dual plastidial and plasma membrane targeting in certain photosynthetic Orobanchaceae, attributable to lamiale-specific duplication of MenA and MenG. Differential losses of *MenA* and *MenG* genes in other photosynthetic, hemiparasitic (top right) and holoparasitic (lower right) lineages, with the latter exclusively PM targeting. pg, plastoglobules.

PM-localized MenA2 and MenG2 likely arose from their plastidial counterparts via gene duplication in the common ancestor of Lamiales (Figure 5). Both *MenA* and *MenG* duplicates are present in *Lindenbergia* and *Rehmannia*, two nonparasitic genera sister to the parasitic Orobanchaceae (McNeal Joel et al., 2013). However, the evolutionary fate of the duplicates varied among parasitic lineages with different photosynthetic capabilities (Figure 5). The holoparasitic *P. aegyptiaca*, *Orobanche fasciculata* and *Conopholis americana* have dispensed with *MenA1* and *MenG1* along with the photosynthetic machinery, retaining only *MenA2* and *MenG2*. On the other hand, the photosynthetic *T. versicolor* and *S. hermonthica* have lost *MenA2* but retain the *MenG1/2* duplicate, as confirmed in the recently released *S. asiatica* genome (Yoshida et al., 2019). In both *T. versicolor* and *S. hermonthica*, the PM-targeted MenG2 became the dominantly expressed isoform over the plastidial MenG1 (Figure 1B), which was subsequently lost in multiple holoparasites. The data suggest expression divergence of the two phylloquinone pathways in the hemiparasites that predated emergence of holoparasites.

An outstanding question in PM phylloquinone biosynthesis regards the co-substrate for MenA-mediated prenylation. In photoautotrophs, the prenyl precursor of plastidial prenylquinones is phytyl-diphosphate derived from geranylgeranyl-diphosphate via *de novo* synthesis or from phytol released upon chlorophyll degradation (Keller et al., 1998; Ischebeck et al., 2006). In *A. thaliana*, the chlorophyll salvage pathway is indispensable for leaf tocopherol and phylloquinone synthesis, but seeds depend on a distinct phytol pool of as-yet-undefined origin (Valentin et al., 2006; Zhang et al., 2014; Wang et al., 2017). *Phelipanche* and related *Orobanche* seeds are filled with protein and oil bodies (Joel et al., 2012), and lipids are the main storage reserve comprising up to 30% of dry seed mass (Bar-Nun and Mayer, 2002). Whether the prenyl moiety of PM phylloquinone is supplied by the cytosolic isoprenoid pathway or via other mechanisms requires further investigation.

### Phylloquinone involvement in PM redox modulation

Involvement of phylloquinone in PM electron transport has been proposed by multiple research groups, based on feeding experiments (Barr et al., 1992; Döring et al., 1992a) and on detection of phylloquinone and QR activities in PM preparations of maize roots and soybean hypocotyls (Lüthje et al., 1998; Bridge et al., 2000; Schopfer et al., 2008). This noncanonical function, however, has not gained widespread acceptance owing to concern over potential organellar contamination in the PM preparations from heterotrophic tissues (Oostende et al., 2008), and to the unexplained biosynthetic origin of PM phylloquinone, until now.

The *P. aegyptiaca* phylloquinone network comprised redox proteins typical of a transmembrane electron transport chain, including PaQR2, PaFRO1, PaCSD1, PaGPX4 and PaPRXs. PaQR2 can function both as a phylloquinone biosynthetic enzyme and an electron transport chain component with demethylphylloquinone or phylloquinone as acceptors. PaFRO1 orthologs have been genetically characterized in *A. thaliana*, and catalyze transmembrane electron transfer associated with Cu(II) reduction in root tips (Bernal et al., 2012). It is worth noting that redox-active Cu is an essential cofactor of several metalloproteins involved in cellular electron transport chains, and Cu proteins play important roles in ROS homeostasis during growth, development and defense (Bérczi and Møller, 2000; Palmer and Guerinot, 2009). This suggests that the transmembrane redox system for Cu homeostasis has been co-opted, along with PM phylloquinone, to support parasitism in *P. aegyptiaca*. PaCSD1, PaGPX4 and PaPRXs likely function synergistically in apoplastic ROS scavenging and signaling associated with PM electron transport. Our data thus provide molecular evidence from parasitic plants to corroborate the long-hypothesized involvement of phylloquinone in PM redox processes in photoautotrophic species (Lüthje et al., 1998; Bérczi and Møller, 2000; Lochner et al., 2003).

### A role for PM phylloquinone in redox signaling associated with parasitism

The early stages of the parasitic lifecycle can be characterized as a continuum of oxidative events, from activation and perception of host HIFs, to induction of haustoria for host penetration and vascular connection (Keyes et al., 2000; Fernández-Aparicio et al., 2016). Redox modulation is also integral to normal growth and development, as well as defense and counter-defense in the case of host-parasite interactions (Kim et al., 1998; Mittler et al., 2004). Redox-active phylloquinone in the PM may function in all of these processes, but most likely in ways specifically associated with haustorium initiation based on network inference and coexpression with known parasitism genes. In the xenognosis model depicted by Keyes et al. (2000), host-derived signals are perceived by a PM-bound electron transport chain for haustorium initiation. Because haustorium signaling shares similar quinone cofactor and redox range requirements with organellar redox circuits (Smith et al., 1996), Keyes et al. (2000) have speculated on the involvement of organelles in the redox relay of haustorium signaling. Our finding that *P. aegyptiaca* exploited phylloquinone from a defunct photosynthetic electron transport chain for PM redox systems is consistent with the model.

Comparative analysis of phylloquinone gene coexpression networks recapitulated the evolutionary shifts in photosynthetic competence, decreasing from facultative (*T. versicolor*) and obligate (*S. hermonthica*) hemiparasites to the obligate holoparasite *P. aegyptiaca*. Conversely, the increasing representation of genes involved in redox processes is positively correlated with the degree of parasitism. This is consistent with the view that some redox-regulated processes in host recognition and haustorium development diversified during the evolution of parasitism (Westwood et al., 2010; Yoshida et al., 2016). *Phelipanche* and related *Orobanche* spp. differ from hemiparasites in many aspects of development and host interactions (Yoshida et al., 2016). For instance, *Phelipanche* and *Orobanche* spp. do not develop the same swollen structure with modified root hairs typical of *S. hermonthica* and *T. versicolor* haustoria, and the haustorium induction chemistry in these holoparasites remains poorly understood (Fernández-Aparicio et al., 2016). Perhaps not surprisingly, taxon-dependent coordination between phylloquinone genes and redox proteins was evident. Phylloquinone-coexpression with chloroplastic ROS scavengers in the hemiparasites is consistent with retention of the plastidial pathway for photosynthetic electron transport. The coordination between components of PM phylloquinone electron transport chain and apoplastic ROS metabolism in *P. aegyptiaca* again attests to the increased specialization of signaling mechanisms associated with holoparasitism.

In closing, the previously proposed involvement of phylloquinone in PM electron transport gains molecular support in this study. Our work highlights a novel link between PM phylloquinone and parasitism that warrants future experimental verification. Repurposing phylloquinone for PM redox signaling associated with haustorium development thus adds to documented evolutionary innovation of parasitic plants. The existence of the alternative pathway in nonparasitic lamiales should also motivate research into the functions of PM phylloquinone in photoautotrophic species.

## Methods

### Transcriptome assembly of parasitic plants

RNA-Seq data of *P. aegyptiaca*, *S. hermonthica* and *T. versicolor* were downloaded from the Parasitic Plant Genome Project database (http://ppgp.huck.psu.edu/). After quality control filtering, cleaned reads were assembled using the custom PLAS pipeline. Briefly, PLAS performs reference-guided read mapping using Bowtie 2 v2.2.3 (Langmead and Salzberg, 2012) against the closely related *Mimulus guttatus* proteome binned by homology to facilitate parallel computing. Mapped reads were used for *de novo* assembly by Trinity (Haas et al., 2013), and the assembled contigs were quality-checked against the reference in each bin by Blast before being used as baits in the next round of *de novo* assembly. This process was repeated for up to 10 iterations until the output was stable. Assembled sequences were filtered to remove potential contaminations (*e.g.*, host plants, protozoa, invertebrates, bacteria, fungi and human sequences) and redundant contigs sharing ≥95% sequence identity. The transcriptome was annotated against *A. thaliana* and *M. guttatus* proteomes and UniProt database. Transcript abundance was estimated using eXpress 1.5.1 (Roberts and Pachter, 2013). Additional *MenA* and *MenG* sequences were obtained from the 1KP database (https://db.cngb.org/onekp) by BlastN against the Core Eudicots/Asterids clade using *P. aegyptiaca* sequences as query. The assembled transcriptome sequences of parasitic plants, the PLAS pipeline and other codes are available at https://github.com/TsailabBioinformatics/PM-PhQ. All phylloquinone and relevant transcripts described in this work were manually curated (Supplemental Dataset S1). *PaMenA2* and *PaMenG2* were further confirmed by RT-PCR. Briefly, cRNA derived from RNA isolated from stages 2 and 3 as described in Yang et al. (2015) was reverse transcribed using SuperScript™ IV reverse transcriptase (Invitrogen) with mixed random hexamers and gene-specific reverse primers (Supplemental Table S2). PCR was performed using Q5 High-Fidelity DNA Polymerase (NEB) with gene-specific primers (Supplemental Table 2), column purified for Gibson assembly into pUC19 using NEBBuilder® HiFi DNA Assembly Master Mix, and confirmed by Sanger sequencing (Eurofins Genomics).

### Subcellular and transmembrane domain prediction and gene structure

Phylloquinone gene sequences of photosynthetic species were downloaded from Phytozome v11. Subcellular localization was predicted by Predotar 1.04 (Small et al., 2004), TargetP 1.1 (Emanuelsson et al., 2007), Protein Prowler 1.2 (Bodén and Hawkins, 2005), and WolF PSORT (Horton et al., 2007). Transmembrane domain was predicted by TMHMM Server v.2.0 (Krogh et al., 2001), and data plotted in *R*. Gene structures were drawn by Gene Structure Display Server 2.0 (Hu et al., 2015). The exon-intron junctions of parasitic *MenA* and *MenG* were inferred by Blast alignment of transcripts against genomic short read data of *P. aegyptiaca* (NCBI Sequence Read Archive accession SRX2067908), *S. hermonthica* (SRX2067907) and *T. versicolor*. Sequence alignment was performed using MUSCLE 3.8.31 (Edgar, 2004) and visualized with Color Align Conservation (www.bioinformatics.org/sms2/color_align_cons.html).

### Coexpression analysis

Transcripts with an FPKM ≥2 in at least two samples were subject to pair-wise Gini Correlation Coefficient (GCC) computation using a python script. Phylloquinone-coexpressed transcripts were defined as the 500 most highly correlated transcripts or those with a GCC ≥0.8 for each phylloquinone gene, excluding the alternative *NDC1* and *QR2*. The union sets were subjected to Gene Ontology enrichment analysis using topGO 2.26.0 (Alexa et al., 2006). To facilitate comparative analysis between species, ortholog groups were constructed by OrthoFinder 1.0.8 (Emms and Kelly, 2015). Network visualization was performed in Cytoscape 3.4.0 (Shannon et al., 2003) using prefuse force-directed layout, with a GCC cutoff of 0.6.

### RNA-seq data processing of photoautotrophic species

RNA-seq data of *A. thaliana*, *G. max* and *Populus tremula* × *alba* were downloaded from the NCBI SRA and quality control processed by Cutadapt 1.9.dev1 (Martin, 2011) and Trimmomatic 0.32 (Bolger et al., 2014). Reads were mapped by Tophat 2.0.13 (Kim et al., 2013), alignment sorted by Samtools 1.2 (Li et al., 2009), and read count and expression estimation obtained by HTseq 0.6.1p1 (Anders et al., 2015) and DESeq2 (Love et al., 2014). *A. thaliana* data used for GCC computation (excluding stressed samples) were SRA236885, SRA091517, SRA269936, SRA219425, SRA308579, SRA050132, SRA067724, SRA291734, SRA269101, SRA098075, SRA100242, SRA122395, SRA163488, SRA064368, SRA246225, SRA248861, SRA202878, SRA201550, SRA303151, SRA221137, SRA272654, and SRA221060. *G. max* data included SRA187830, SRA047293, SRA036577, SRA116533, SRA091756, SRA187830, SRA036538, SRA036577, and SRA129337. *P. tremula* × *alba* datasets were SRA274261 and SRA097208.

### Phylogenetic tree construction

The protein sequences of *P. ramosa* PrPRX1 (AAY89058) and PrPOX1 (AAU04440), and *S. asiatica* SaPOXA (AAB97853) and SaPOXB (AF043235) were searched against the PLAS assemblies of *P. aegyptiaca*, *S. hermonthica* and *T. versicolor* to identify orthologs. Their protein sequences were aligned by MUSCLE 3.8.31 (Edgar, 2004) and the alignment trimmed by Gblocks (Castresana, 2000). Bayesian phylogenetic tree was constructed using MrBayes 3.2.5 (Ronquist et al., 2012). Phylogenetic trees for MenA and MenG were similarly constructed using protein sequences obtained from Phytozome or 1KP followed by manual curation (Supplemental Dataset S1), and visualized by MEGA X (Kumar et al., 2018).

### Phylloquinone analysis

*P. aegyptiaca* seeds were surface-sterilized, pre-conditioned on moist filter paper for 7-10 days, and treated with GR-24 for 4-6 days before collection of stage 1 germinated seeds as described (Westwood et al., 2012). Non-imbibed *A. thaliana* (Col-0) seeds were used without further treatment. Tissues were snap-frozen in liquid nitrogen, freeze-dried and milled to a fine powder for phylloquinone analysis as described (Booth et al., 1994). Tissues were partitioned twice in isopropanol:hexane (3:2 v/v), with menaquinone MK-4 (Sigma V9378) as a reference for *P. aegyptiaca* analysis. The hexane phase was dried under nitrogen, and resuspended in methylene chloride:methanol (1:4, v/v), 10 mM ZnCl2, 5 mM Na-acetate, and 5 mM acetic acid. Isocratic reverse-phase HPLC (Agilent Eclipse Plus C18 column, 5 μm, 4.6×250 mm) was carried out using methanol:methylene chloride (9:1, v/v), with post-column Zn reduction and fluorescence detection (excitation 244 nm, emission 418 nm). Phylloquinone peaks were verified by comparison with the authentic standard (Sigma 47773) and concentrations were estimated using calibration curves of the standard.

### *Nicotiana benthamiana* transformation and confocal microscopy

*PaMenA* and *PaMenG* coding sequences were gene-synthesize (General Biosystems) and subcloned into *Xba*I-digested pCX-AtMenA.1-GFP vector to generate pCX-PaMenA-GFP and pCX-PaMenG-GFP, respectively. The pCX-AtMenA.1-GFP vector was previously constructed by Gibson assembly of PCR-amplified AtMenA.1 (obtained from Arabidopsis Biological Resource Center) and GFP (from pCXDG) into *Bam*HI-digested pCXSN vector (Chen et al., 2009) (see Supplemental Table S2 for primers). Due to concern over 35S promoter silencing in co-transformation experiments, another PaMenA-GFP construct was prepared by PCR using pCX-PaManA-GFP as template and primers containing a viral 2A peptide bridge sequence (Luke et al., 2015) to link the HPT (hygromycin phosphotransferase) coding sequence to PaMenA-GFP behind double 35S promoter as one transcriptional unit (*35S:PaMenA-GFP-2A-HPT*) (Supplemental Table S2). All constructs were sequence verified. *Agrobacterium*-mediated transformation of *Nicotiana benthamiana* was performed as described and regenerated on selection media (Pathi et al., 2013) using individual constructs (pCX-PaMenA-GFP, pCX-PaMenG-GFP and pCX-AtMenA.1-GFP) or in conjunction with a PM (PIP2A-mCherry) or tonoplast intrinsic protein (γ-TIP-mCherry) marker (Nelson et al., 2007) (pCX-PaMenA-GFP-2A-HPT + AtPIP2A-mCherry, pCX-PaMenA-GFP-2A-HPT + At γ-TIP-mCherry, pCX-PaMenG-GFP + AtPIP2A-mCherry and pCX-PaMenG-GFP + At γ-TIP-mCherry). Root samples from independent transformants were screened under a fluorescence microscope and at least three positive lines were further examined using a Zeiss LSM 880 upright confocal microscope in the Biomedical Microscopy Core at the University of Georgia. GFP signal was detected using an Argon excitation laser (488 nm) and an emission filter at 490-540 nm, and mCherry signal was obtained using a HeNe excitation laser (543 nm) and an emission filter at 547-697 nm.

### *E. coli* complementation

The *PaMenG2* and *EcMenG* (positive control) coding sequences were PCR-amplified (see Supplemental Table S2 for primers) for Gibson assembly into a constitutive expression vector pUCM (Kim et al., 2010), and transformed into *E. coli* DH5α. Sequence-verified plasmids were transformed into *E. coli* strain JW5581 carrying the mutated *menG* (*ubiE*) gene (obtained from the *E. coli* Genetic Stock Center). At least two PCR-confirmed colonies per construct were used for complementation experiments. Approximately 30 mg of freeze-dried bacterial cells were extracted as described (Booth et al., 1994) using a Misonix sonicator. Reverse-phase HPLC conditions were the same as above except with a run time of 40 min.

## Supplemental Materials

Supplemental Table S1. Plastid-targeting prediction for ICS, PHYLLO and NDC1. Supplemental Table S2. List of primers

Supplemental Figure S1. MenE sequence alignment with C-terminal peroxisome targeting signal PTS1.

Supplemental Figure S2. MenB sequence alignment with N-terminal peroxisome targeting signal PTS2.

Supplemental Figure S3. DHNAT sequence alignment with C-terminal peroxisome targeting signal PTS1.

Supplemental Figure S4. MenA sequence alignment. Supplemental Figure S5. MenG sequence alignment. Supplemental Figure S6. QR2 sequence alignment.

Supplemental Dataset S1. PLAS-assembled or 1KP-derived and manually curated transcript sequences of phylloquinone biosynthetic and coexpressed genes described in the manuscript. Supplemental Dataset S2. Lists of transcripts used in coexpression network analysis.

## Acknowledgments

We thank L.-J. Xue for the GCC Python script, M. Curley, M. Tsai, N. Rodman, and K.B. Aulakh for laboratory assistance, M.K. Kandasamy for confocal microscopy assistance, and D. Lynn (Emory University) for insightful discussion. This research was supported by Georgia Research Alliance-Hank Haynes Forest Biotechnology Endowment (C.-J.T.), National Science Foundation grants IOS-1444567 (C.-J.T.) and IOS-1238057 (J.H.W.), National Institute of Food and Agriculture grants 2015-67013-22812 (C.-J.T.) and VA-135997 (J.H.W.), and University of Georgia Graduate School’s Innovative and Interdisciplinary Research Grant (X.G.).

## Author contributions

C.-J.T and X.G. conceived the study; X.G. performed all bioinformatic analyses with contributions from C.-J.T.; I.-G.C. performed bacterial complementation, tobacco transformation and microscopic analyses with help from M.A.O.; S.A.H, B.N., and I.-G.C. performed metabolic analyses; K.C. and J.H.W. contributed materials; X.G. and C.-J.T. wrote the manuscript with input from S.A.H and J.H.W.

