## Supplemental Material for "Plasma Membrane Phylloquinone Biosynthesis in Nonphotosynthetic Parasitic Plants"

**Supplemental Table 1.** Plastid-targeting prediction for ICS, PHYLLO and NDC1

| | Predotar | TargetP | Protein Prowler | WoLF PSORT | Prediction |
|---|---|---|---|---|---|
| ***ICS*** | | | | | |
| AT1G18870* | 0.79 | 0.95 | 0.97 | 13 | plastid |
| AT1G74710 | 0.82 | 0.77 | 0.99 | 11 | plastid |
| Potri.012G070000 | 0.21 | 0.27 | 0.07 | 9.5 | plastid |
| Solyc06g071030 | 0.11 | 0.52 | 0.08 | 5 | plastid |
| Migut.I00130 | 0.70 | 0.16 | 0.08 | 10 | plastid |
| Migut.I00129 | 0 | 0.03 | 0 | 13 | plastid |
| TvICS | 0 | 0.18 | 0 | 4 | - |
| ShICS | 0.01 | 0.30 | 0.07 | 6 | - |
| PaICS | 0.18 | 0.22 | 0.03 | 8 | plastid |
| ***PHYLLO*** | | | | | |
| AT1G68890* | 0.37 | 0.94 | 0.89 | 12 | plastid |
| Potri.010G135500[1] | 0.07 | 0.58 | 0.92 | 14 | plastid |
| Solyc04g005190[1] | 0.24 | 0.46 | 0.31 | 12 | plastid |
| Migut.L01140 | 0.20 | 0.64 | 0.31 | 13.5 | plastid |
| TvPHYLLO | 0.37 | 0.45 | 0.89 | 11 | plastid |
| ShPHYLLO | 0.57 | 0.74 | 0.98 | 12 | plastid |
| PaPHYLLO | 0.63 | 0.40 | 0.28 | 13 | plastid |
| ***NDC1*** | | | | | |
| At5g08740* | 0.4 | 0.9 | 1.0 | 11.5 | plastid |
| Potri.007G098700 | 0.6 | 0.8 | 1.0 | 11.5 | plastid |
| Solyc03g043750 | 0.9 | 0.9 | 0.9 | 13.5 | plastid |
| Migut.L00271 | 1.0 | 0.9 | 1.0 | 12 | plastid |
| Migut.L00582 | 1.0 | 0.9 | 1.0 | 12 | plastid |
| TvNDC1 | 1.0 | 0.8 | 0.8 | 13 | plastid |
| ShNDC1 | 1.0 | 0.7 | 1.0 | 14 | plastid |
| PaNDC1 | 0.9 | 0.9 | 1.0 | 12 | plastid |

Heatmaps show prediction strengths for scores above the 50$^{th}$ percentile of each method
*Experimentally verified for plastid-targeting.
[1] GenBank accession numbers XP_024466568 (poplar) and XP_004237229 (tomato) were used due to erroneous gene model annotation in the reference genomes.

### Supplemental Table S2. List of primers

Primer tails (overlapping sequences) for Gibson assembly are underlined.

| Gene/vector | Primer sequence (5'-3') | Purpose |
|---|---|---|
| PaMenA2 | GGACCAGATCCCAAGCAAATTA | RT-PCR cloning |
| PaMenA2 | GTTGTAAAACGACGGCCAGTGAATTCGTAGAGTGTGCAAAGACTCRGA | RT-PCR cloning, pUC19 tail |
| PaMenA2 | CTATGACCATGATTACGCCAAGCTTGGACCAGATCCCAAGCAAATTA | RT-PCR cloning, pUC19 tail |
| PaMenG2 | TTAACAARATCCACTGGATCG | RT-PCR cloning |
| PaMenG2 | CTATGACCATGATTACGCCAAGCTTGAGACCGAAAAKCGAAAATGGC | RT-PCR cloning, pUC19 tail |
| PaMenG2 | GTTGTAAAACGACGGCCAGTGAATTCCAAACACGAACRCAAAGCA | RT-PCR cloning, pUC19 tail |
| pUC19 | AAGCTTGGCGTAATCATGGTCATAG | pUC19 vector |
| pUC19 | GAATTCACTGGCCGTCGTTTTACAAC | pUC19 vector |
| PaMenG2 | TCTAGAAGGAGGATTACAAAATGGCTACAGCTACACTTAG | pUCM cloning, vector tail |
| PaMenG2 | GTTGTAAAACGACGGCCAGTGAATTCCTAGCGCGTGGCGACCAAATTTC | pUCM cloning, vector tail |
| EcMenG | TCTAGAAGGAGGATTACAAAATGGTGGATAAGTCACAAGAAAC | pUCM cloning, vector tail |
| EcMenG | GTTGTAAAACGACGGCCAGTGAATTCAGAACTTATAACCACGATGC | pUCM cloning, vector tail |
| pUC19 | CATTTTGTAATCCTCCTTCTAGACGCTCACAATTCCACACAAC | pUCM vector |
| pUC19 | GAATTCACTGGCCGTCGTTTTACAAC | pUCM vector |
| GFP | AGAAGACTGAAGTTAGTAGCTCGAGATCTGAGTCCGGACT | PaMenA2-GFP-2A-HPT cloning, 2A tail |
| HPT | GCTACTAACTTCAGTCTTCTGAAGCAGGCTGGAGATGTGGAGGAGAACCC TGGACCAATGAAAAAGCCTGAACTCACC | PaMenA2-GFP-2A-HPT cloning, 2A tail |
| AtMenA.1 | CAATCAAGCATTCTACTTCTAGATGTACTCTGTCGCTTTAGTTC | pCX-AtMenA.1-GFP cloning, pCXSN (35SP) tail |
| AtMenA.1 | AAAGTTCTTCTCCTTTACCCATTCTAGAGCGCGCGATAACTAGCCCCAA | pCX-AtMenA.1-GFP cloning, GFP tail |
| GFP | ATGGGTAAAGGAGAAGAACTTT | pCX-AtMenA.1-GFP cloning |
| GFP | GAACGATCGGGGAAATTCGCTATCTGGCTTTTAGTAAGCCCC | pCX-AtMenA.1-GFP cloning, pCXSN (NosT) tail |

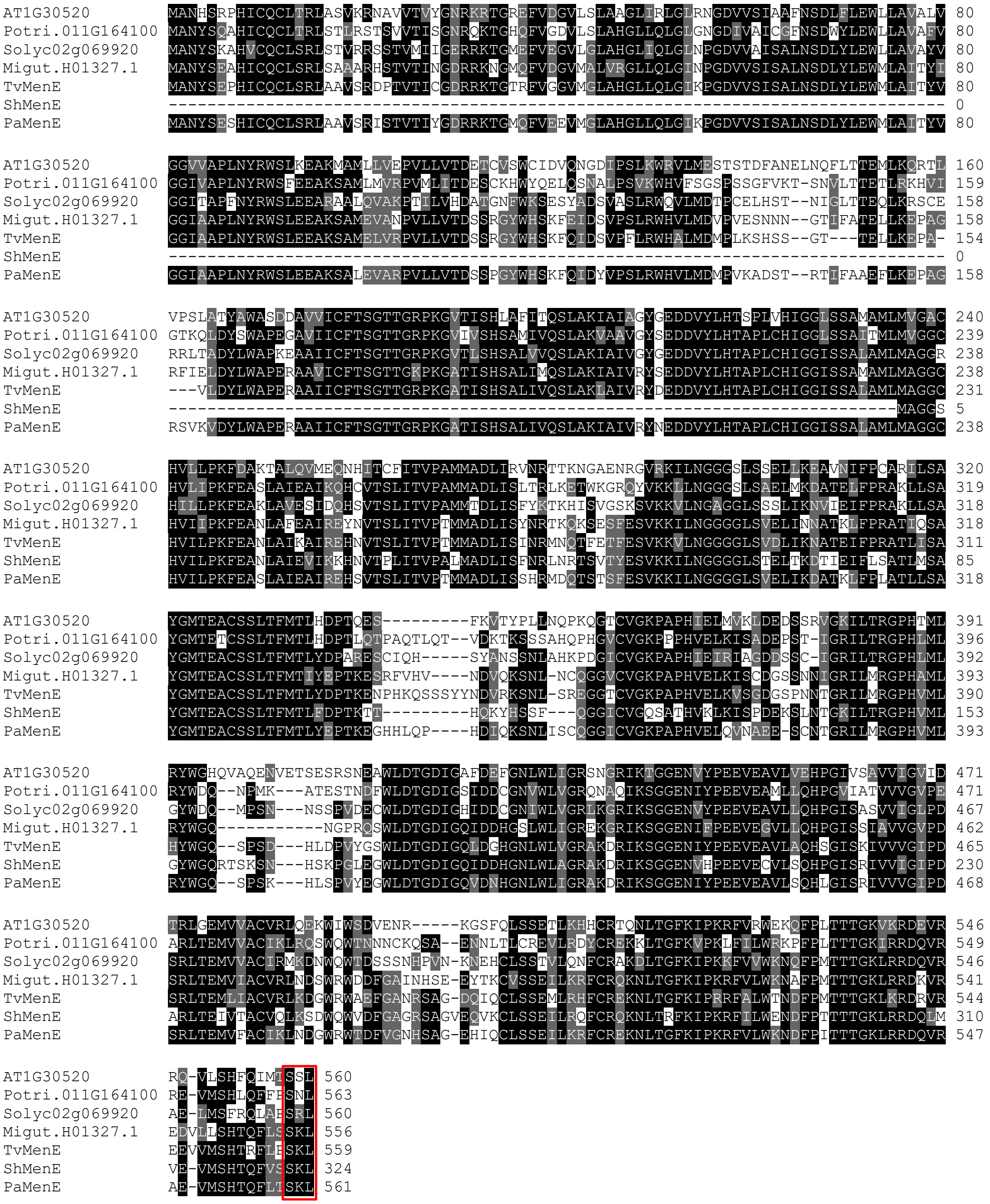


**Supplemental Figure 1. MenE sequence alignment with C-terminal peroxisome targeting signal PTS1**

MenE sequences from the three parasitic species and representative non-parasitic plants from Phytozome are shown. The C-terminal PTS1 is boxed in red. The assembled *ShMenE* transcript sequence contains stop codons in the 5'-end, leading to truncated polypeptide sequence.

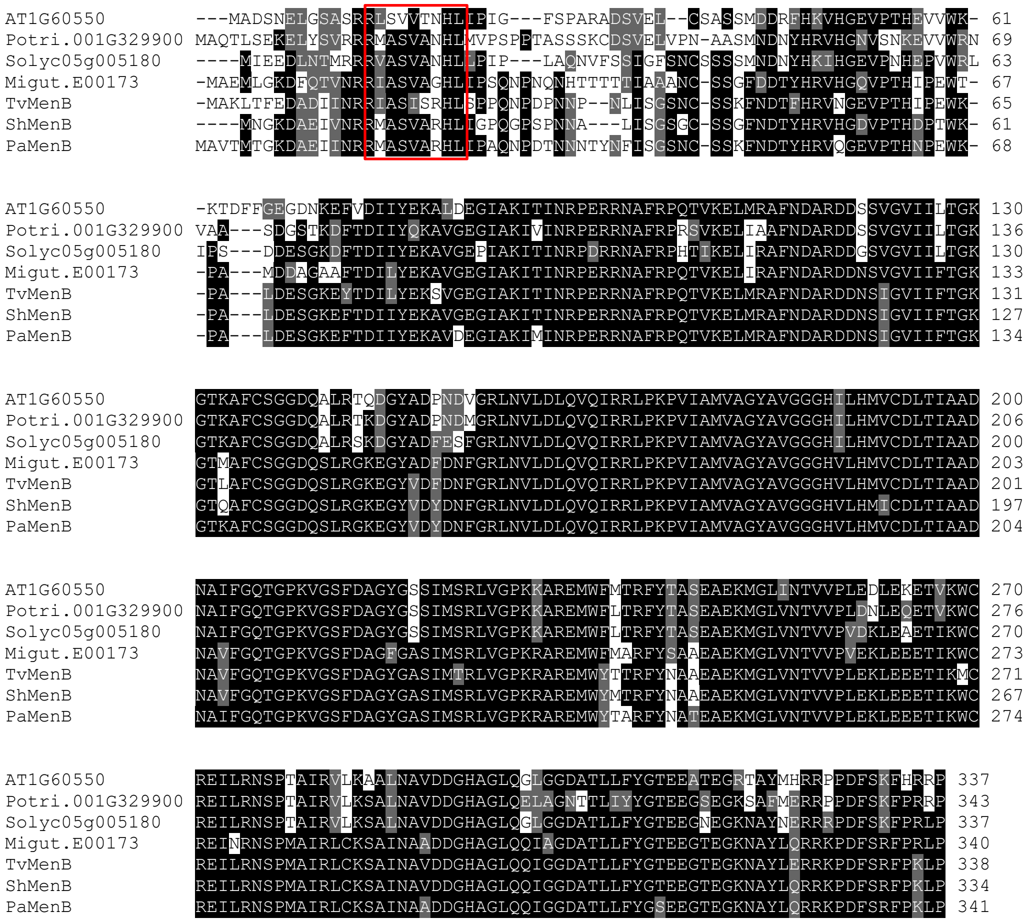


**Supplemental Figure 2. MenB sequence alignment with N-terminal peroxisome targeting signal PTS2**

MenB sequences from the three parasitic species and representative non-parasitic plants from Phytozome are shown. The N-terminal PTS2 is boxed in red.

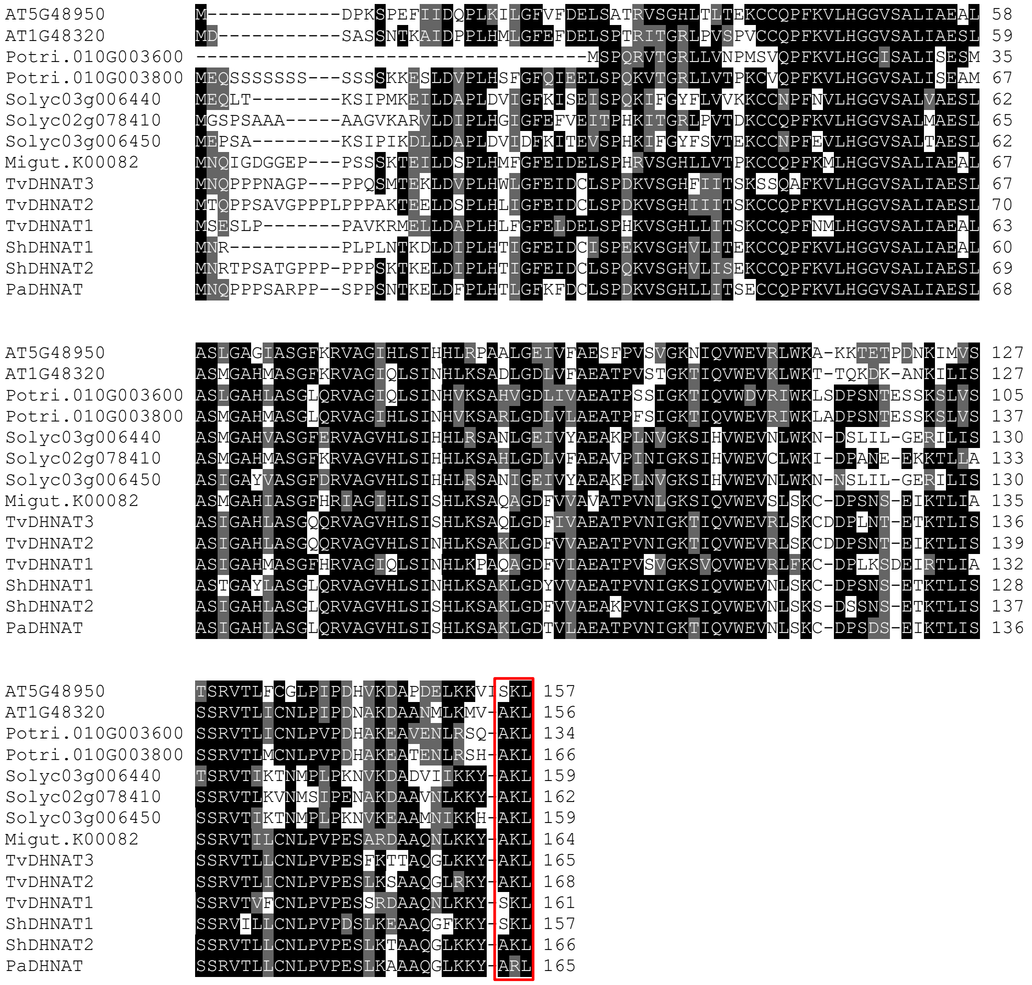


**Supplemental Figure 3. DHNAT sequence alignment with C-terminal peroxisome targeting signal PTS1**

DHNAT sequences from the three parasitic species and representative non-parasitic plants from Phytozome are shown. The C-terminal PTS1 is boxed in red.

```
At1g60600          ----MVNFVSLC--DIKYGFVPKNSTDLFVKRKI-----HKLPSRGDVITRLPVFGSNARENLNAKPRRNLRVRPIFCKSY--------- 70
Potri.017G050900   ------MAAAFC--SLHQGSSLKKLQYYHVRQSYATRSYPGALTSNASKKSSLCCCTAQNFVCGSRISRLEFKKGIQCWRAFSGSIANYA 82
Solyc01g105460     ----MASSATFNVPILVSAFNKTSHAPISSRNFILRTC----SLPSHVRQRSICCFQCTGRSGSVIDLGWKEVKQWHSSLRGFFMCKVQQ 82
Migut.B00584       MAMAAAAAATFC--SISHGYGVTKLNSF---RICIYSRYQSRAITHENVFRL---DARK----------LHSIQRRYIIPLKPRASSN-- 70
VtMenA1            -----MAAANFC--SMSHGYGVKNLNQFLPRRCNNYRAYQELSPQDAVRMPL---CSKTYFDKTIIRR-LHSKKPHHRILFKCAAELI-- 77
LpMenA1            -----MAAATFC--SISHGYGVSRVNDYLLNRHGINRIDQVLPLPDASRRRK---LVKKDFNKTVIRQ-LHSIRRHYTMSLKFRAENN-- 77
RgMenA1            -----MAAATFC--SISHGYGVNKLNQHLLNRHNFNRIYQVIPLPDASRRQL---RMEMCFDKTIIRQ-LHCIRRHYGILLKCRSEHN-- 77
TvMenA1            -----MAAATFCSISISHGYAVQRLN-----RHKINRTYQVLPLVCGSRTTK------VHLNKTIIRQYLHSIQRRYKISPKFRSENNAD 74
ShMenA1            MAPLAVAAAVYC--STSHGYGVKKLDDYLTRKLSISRIHQVLLLPDACQRSL---CTKFNFKKASMRQ-LYSLRGHYINSFKQRAEHCG- 83
Migut.B01155       ----MLS-------SIQENYAVTG------------------NLHADDVTPE-------------------------------------- 23
Migut.B01157       ----MLS-------SIQENYAVTG------------------NLHADDVTPE-------------------------------------- 23
RgMenA2            -----MAGTSF---SSSDN-----------------------PSKEKNGKRS---KKE-------------------------------- 24
LpMenA2            -----MAGTSL---NLVEN-----------------------ASKEKNIKRS---KNGS------------------------------- 25
PaMenA2            ----MAEAANH-------------------------------PNQANIVKRL-------------------------------------- 17
CaMenA2            ----MAGGANFI--SLADN-----------------------PSLEKNVKRS-------------------------------------- 23
OfMenA2            ----MAGGSSI---SLADN-----------------------PTQEKVVRRS-------------------------------------- 22

At1g60600          GDAAKVYQE-EE----IPRAKLIWRAIKLPMYSVALVPLTVGASAAYLETGLFLARRYVTLLLSSILIITWLNLSNDVYDFDTGADKNKM 155
Potri.017G050900   DKSHEGETG-ED----VPKATLIWRAVKLPIYSVALVPLTVGGAAAYLQTGMFSARRYFSLLVSSILIITWLNLSNDVYDFDTGADKNKK 167
Solyc01g105460     HDATSEENE-EN----ISQATLIWRAIKLPIYSVALIPLTVGSAAAYLQTGLFSARRYFMLLASSVFIITWLNLSNDVYDFDTGADKDKK 167
Migut.B00584       EDHIQHEKEVNN----ISKATLIWRAVKLPMYTVAFIPISVGSAAAYLQTGKYFAKRYFVLLVSSVLVIAWLNLSNDVYDFDTGADKNKK 156
VtMenA1            DTHGEDEKE-EH----ISKATLIWRAIKLPIYTVALIPITVGNAAAYLQTGQYYGKRYVMLLVSSFLIIAWLNLSNDVYDFDTGADKNKK 162
LpMenA1            GDRIEEEKE-GN----VSKATLIWRAIKLPIYTVALIPITVGSAAAYLQTGQYFGKRYVMLLISSVFVIAWLNLSNDVYDSETGADKNKK 162
RgMenA1            DSHVVEEKE-ES----ISKATLIWRAIKLPMYTVALIPITVGSAAAYLQTGQYFGKRYLMLLISSVLIIAWLNLSNDVYDFETGADKNKK 162
TvMenA1            NTHVEEEEEDEDEQESVSKATLIWRAIKLPMYTVALIPITVGSAAAYLQTGQYFGKRYIMLLVSSVLIIAWLNLSNDVYDFDAGADKNKK 164
ShMenA1            SSNIEENKE-ES----ISRVALMWRAIKLPIYSVALIPITVGSAAAYLQTGQFFGKRYLKLLLSSVLIITWLNLSNDVYDFETGADKNKK 168
Migut.B01155       --DETKTRD-DD----VSRATLIWRAAKLPMYTVALIPVTVGTAAAYWQTGLYSSARYFKILASYVLVIVWLNLSNDVYDFDTGVDVNKK 106
Migut.B01157       --DETKTRD-DD----VSRATLIWRAAKLPMYTVALIPVTVGTAAAYWQTGLYSSARYFKILASYVLVIVWLNLSNDVYDFDTGVDVNKK 106
RgMenA2            TTSHDEKKE-ED----IGRATLIWRAAKLPMYTVALIPLTVGTAAAYWESGYYSLERYFILLASFVLVNLWVNLSNDVYDFDTGADINKK 109
LpMenA2            TSPNEEKNE-EE----ISRATLIWRAAKLPMYTVALIPLTVGTSAAYWESGYYSLERYFILLASFVLVNVWVNLSNDVYDFDTGADKNKK 110
PaMenA2            --HKKEKKE-GD----ISRATLIWRAAKLPMYTVALTPLTVGTAAAYWESGFYSLERYFTLLASFVLVNLWINLSNDVYDFDTGADKNKK 100
CaMenA2            KKKGEKKEE-ED----ISRATLIWRAAKLPMYTVALIPLTVGTATAYWESGYYSLERYFILLASFVLVNVWINLSNDVYDFDTGADKNKR 108
OfMenA2            ----KSKKE-ED----ISRAILIWRAAKLPMYTVALIPLTVGTAAAYLESGFYSLERYFILLASFVLVNVWVNLSNDVYDFDTGADKNKK 103

At1g60600          ESVVNLVGSRTGTLAAAITSLALGVSGLVWTSLNASNIRAILLLASAILCGYVYQCPPFRLSYQGLGEPLCFAAFGPFATTAFYLLLGSS 245
Potri.017G050900   ESVVNLVGSRSVAFIAAYSSLLLGFAGLAWTSMGEGNIHAILFLACAILCGYVYQCPPFRLSYQGLGEPLCFAAFGPFATSAFYLLLGST 257
Solyc01g105460     ESVVNIVGSRTGTFVAACLLLALGFLGLTYTSVEAANLRAILLLASAITCGYIYQCPPFRLSYQGLGEPLCFAAFGPFATTAFYLLQSNT 257
Migut.B00584       ESVVNLLGSRTGAHVIAWLLLGIGFTGLTLVSIEAKSIRSILLLACAVVCGYIYQCPPFRLSYMGLGEPLCFAAFGPLATTAFYLLQSGR 246
VtMenA1            ESVVNLIGSRKGTRAFAWSLLALGFMGLTRVSMDAGSFYPLFLLACAIICGYIYQCPPFRLSYMGLGEPLCFAAFGPFATTAFYFLQTGT 252
LpMenA1            ESVVNLFGSRTGTHVFAWLLLALGFTGLTLVSVEARSRHSIFLLACSVFCGYVYQCPPFRLSYLGLGEPLCFAAFGPLATIAFYLLQGGT 252
RgMenA1            ESVVNLFGSRTGTHVFAWLLLALGFTGLTWVSVEAGSLRSIFLLGCAVFCGYIYQCPPFRLSYLGLGEPLCFAAFGPFATTAFYLLQSGT 252
TvMenA1            ESVVNLFGSRTGTHVFAWLLLALGFAGLARVSVEAGSLRSIFLLGCAVFCGYIYQCPPFRLSYMGLGEPLCFVAFGPFATTAFYLLQSGT 254
ShMenA1            ESVVNLIGSQTGTHILAWVLLELGFGGLTWVSIEAGSIRSIILLACAIFCGYIYQCPPFRLSYLGLGEPLCFAAFGPFATTAFYLLQSGT 258
Migut.B01155       ESVINLFGSRTGINIIAWLILVLGFLGLAWVAVEARNPRSMILLASAVFCFFLYQCPPFRLSYYGVGEPLLFVAFGPLSTIAFYLLHSIK 196
Migut.B01157       ESVINLFGSRTGINIIAWLILVLGFLGLAWVAVEARNPRSMILLASAVFCSFLYQCPPFRLSYYGVGEPLLFVAFGPLSTIAFYLLHSIK 196
RgMenA2            ESVVNIVGSRNAIHILSWLLLAIGFAGLSWVGVEAKNPRAILLLAAAVFCGYIYQCPPFRLSYHGLGEPLCFASFRPFSTVAFYLLQSRS 199
LpMenA2            ESVVNIVGSRTAIHILSWLILALGFAGLTWVGVEAKNPRAILLLASAVFCGYIYQCPPFRLSYHGLGDPLCFAAFGPFSTVAFYLLQSSS 200
PaMenA2            ESVVNIFGSRTAIHILSLLVLAIGFAGLCWVALEAKNPRAILLLASTVFCVYAYQFPPYRLSYHGLGEPLCFAAYGPFATVAFYLLQSGS 190
CaMenA2            ESVVNMVGSRTAIYILSWLLLALGFAGLSWVGIEAKNPRAILLLASSIFCVYIYQCPPYRLSYHGLGEPICFAAYGPFSTIAFYLLQSTS 198
OfMenA2            ESVVNIVGSRTAIHILSWLLLALGFAGLCWVGLEANNPRAILLLASAIFCGYIYQCPPFRLSYHGLGEPLCFASFGPFSTIAFYLLQSSS 193

At1g60600          S-EMRHLPLSGRVLSSSVLVGFTTSLILFCSHFHQVDGDLAVGKYSPLVRLGTEKGAFVVRWTIRLLYSMLLVLGLTRILPLPCTLMCFL 334
Potri.017G050900   S-EMSILPLTGTILSASLLVGFTTTLILFCSHFHQVEEDKAVGKFSPLVRLGTERGSGVVKVAVAMLYSLLFASGLSRTLPLACILLCSL 346
Solyc01g105460     S-E---LPITYTTLSASALVGFTTSLILFCSHFHQVEGDKAVRKFSPLVRLGTEAGSRVVKVGVATLYSGMLLLGFTQTLPLTCIILCAL 343
Migut.B00584       S-E---LPVSGTIVSSSILVGLTTSLILLCSHFHQIEDDKAVGKISPLVRLGTERGAKVVNVAVGLLYSLVFLLGLNQTLPFFSLVLCAL 332
VtMenA1            R-D---LSISATVISSSILVGITTSLILFCSHFHQIDDDKTVGKLSPLVRLGTETGAKVVKIAVITLYSLLFVLGLSQTLPFSSIVLCAL 338
LpMenA1            R-E--QLSISCAVVASSILVGFTTSLILFCSHFHQIEDDKAVGKFSPLVKLGTEGGAKVVKVAIGTIYSLLFILGLSQTLPFSSIVLCAL 339
RgMenA1            R-E---LSISVTIVSSSILVGLTTSLILFCSHFHQIDDDKAVGKFSPLVRLGSDGGAKVVKVAVGTLYSLLFILGLSQTLPFSSIVLCAL 338
TvMenA1            R-E---LSISGIVIVSSVLVGITTSLILFCSHFHQIEDDKAVGKYSPLVRLGSEGGANVVKMVVRTIYSLLFILGLVQTLPFASIVLCAL 340
ShMenA1            S-E---LSISATVISSSILVGFTTSLILFCSHFHQIEDDKAVGKFSPVVRLGTEGGANVVKVAVRILYSLLFVLGLAQILPFPSIVLCAL 344
Migut.B01155       S-E---LPITSTIVWSSILVGLTTSLILFCSHFHQIKDDKAVGKMSPLVRLGTETGSRVVKFAVVGLYSIILVLGFTQNLPLASVILCAM 282
Migut.B01157       S-E---LPITSTIVWSSILVGLTTSLILFCSHFHQIKDDKAVGKMSPLVRLGTETGSRVVKFAVVGLYSIILVLGFTQNLPLASVILCAM 282
RgMenA2            S-E---LPISSTIISAAILVGFTSALILFCSHFHQIEDDKAVGKISPLVRLGTETGSKVVKMAVLGLYWLVFGLGLAQTLPYPCIMLCIM 285
LpMenA2            S-E---LPISSTIVSSAILVGFTSALILFCSHFHQIEDDKAVGKISPLVRLGTEKGSKVVKMAILGLYWLVLGLGLAQTLPYACIVLCAM 286
PaMenA2            SSE---LPISSTVVFASIFVGFTSALILFCSHFHQIEDDKAVGKISPLVRLGTEKASKVVKMSVLGFYWLVFGLGLSQTLPYACVVLCTM 277
CaMenA2            KFE---LPISSTIVSAAILVGFTTALILFCSHFHQIEDDKAVGKISPLVRLGTEKASKVVKMAVLGLYWLVFGFGLTQTLPYACLVLCTM 285
OfMenA2            S-E---LPLSSTIVSAAILVGSTSALILFCSHFHQIEDDKAVGKISPLVRLGTEKASEIVKMAVLGLYWLVFGLGLSQTLPYACIVLCTM 279
```

```
At1g60600          TLPVGNLVSSYVEKHHK--DNGKIFMAKYYCVRLHA---LLGAALSLGLVIAR------------------------ 382
Potri.017G050900   TLPMGKLVVGFVEENYK--DKGKIFMAKYFCVRLHA---LFGAALASGLVAAR--VFQG-YFP--------------- 401
Solyc01g105460     TIPVGKLVVSYVEKNHKVRQSKDLYGKVLLCETSCSIWSGSGCWLGSC----------------------------- 391
Migut.B00584       TLPVGRLVSSFVEKNHK--DETKIFMAKYYCVRLHT---VFGGALAAGMVAAR--MFARRQLPHTVLL--------- 393
VtMenA1            TLPIGNKIVRFVEENHK--DKTKIFMAKYYCVRLHT---VFGAALAAGLVGAK--MFARKKIPHAMIIS-------- 400
LpMenA1            TLPMGKFVVSFIEENHK--DKTKIFMAKYYCVRLHT---VFGAALAAGLVAAR--MLARKPLPYAVIL--------- 400
RgMenA1            TLPVGKLVVSFVEKNHK--DKTKIFMAKYYCVRLHT---VFGAALAAGMVAAR--MFARKQLPHAIIL--------- 399
TvMenA1            TLPVGNLVVSFVEKNHK--DKTKIFMAKYYCVRLHT---VFGAALATGLVAAR--VLARKPIPNAIIL--------- 401
ShMenA1            TLPVGNLVVSFVGTNHK--DKMKIFMAKYYCVRLHT---IFGAALAAGMVAAR--MFARKQLPHAIIL--------- 405
Migut.B01155       TLPMGNLVVSFVEKNHK--DKTKIFMGKYLCVRLHT---VFGAALAAGLVVAR--KFAVQQLPHAIVP--------- 343
Migut.B01157       TLPMGNLVVSFVEKNHM----VNIHGEVFVCEVAHCFWGCFGCWACSG----------------------------- 326
RgMenA2            TLPMGNLVVSFVQENHK--DKSKIFMAKYYCVRLHT---VFGAALAAGLVASRL-IQSGEQLQSLSTYFDRAKFAYF 356
LpMenA2            TLPMGNLVVSFVQENHK--DKSKIFMAKYYCVRLHT---VFGAALAVGLAASR--INGGDYLQSPSTYFDRAKFAYF 356
PaMenA2            TLPMGNLVVSFVQENHK--DKSKIFMAKYYCVRLHT---VFGAALAAGLVASRLIMHSGEQLQQSI--FDYAKFLYL 347
CaMenA2            TLPIGNLIVSFVQENHK--DKSKIFMAKYYCVRLHT---VFGSALAAGLVASR--MHSGELTSEFINVLRATKFAYF 355
OfMenA2            TIPMGNLVVSFVQKNHK--DKSKIFMAKYYCVRLHT---VFGAALAVGLVASRL-MHSGEQFQSLLTYFDRAEIA-- 348
```

**Supplemental Figure 4. MenA sequence alignment.**
MenA sequences from the three parasitic plants were aligned with representative MenAs of non-parasitic plants from Phytozome and additional sequences identified from 1KP (Supplemental Dataset S1). Of, *Orobanche fasciculata*; Ca, *Conopholis americana*; Rg, *Rehmannia glutinosa*; Lp, *Lindenbergia philippensis*; and Vt, *Verbascum thapsus*.

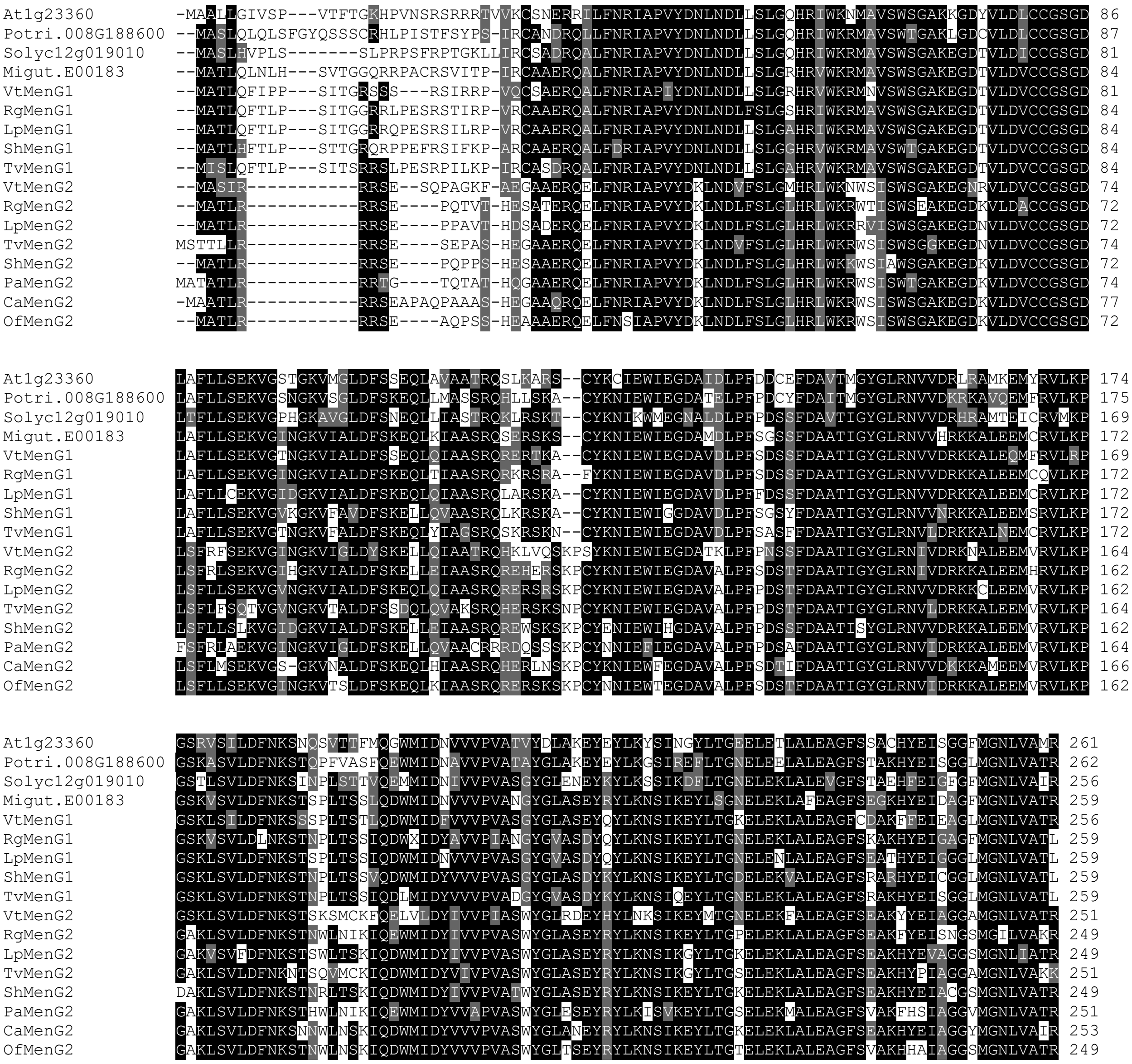


**Supplemental Figure 5. MenG sequence alignment.**

MenG sequences from the three parasitic plants, representative non-parasitic plants from Phytozome and additional species from 1KP (Supplemental Dataset S1) are shown. Of, *Orobanche fasciculata*; Ca, *Conopholis americana*; Rg, *Rehmannia glutinosa*; Lp, *Lindenbergia philippensis*; and Vt, *Verbascum thapsus*.

```
Potri.011G033100 MAVKIYIIYYSMYGHVARLAEEIKKGADTVEGVEIKLWQVPETLPEEVLGKMGAPPK-SDVPIIKPNDLTEADGVLFGFP 79
Migut.M01839     MATKVYIVYYSMYGHVEKLAEEIKKGAASVEGVEAKLWQVPETLQEDVLTKMSAPPK-SEVPIITPSELAEGDGFIFGFP 79
AT5G54500        MATKVYIVYYSMYGHVEKLAEEIRKGAASVEGVEAKLWQVPETLHEEALSKMSAPPK-SESPIITPNELAEADGFVFGFP 79
AT4G27270        MATKVYIVYYSMYGHVEKLAQEIRKGAASVDGVEAILWQVPETLQEDVLSKMSAPPK-SDAPIITPNELAEADGFIFGFP 79
Potri.001G410700 MATKVYIVYYSMYGHVEKLAEEIKKGASSVEGVEAQLWQVPETLPEEVLGKMSAPPK-SDVPIITPGELAEADGFVFGFP 79
Potri.011G129400 MATKVYIVYYSMYGHVEKLAEEIRKGASSVEGVEAKLWQVPEILPEEVLGMMSAPPK-SDVPIITPSELAEADGFVFGFP 79
TvQR2            MATKVYIVYYSTYGHVERLAQEIKKGAESVGNVEVKLWQVPEILSDEVLGKMWAPPK-SDVPVITPDELVEADGIIFGFP 79
ShQR2            MAAKVYIVYYSTYGHVERLAQEIKKGVESIPEVEAKLWQVPETLTNDILAKMGAPPKNNDVPVISPNELVDADGIIFGFP 80
PaQR2            MATTLSIVYYSTYGHVEKLAQEIKKGAESVEGVEAKLWQVPETLSDEILGKMGAPPKSSEVPVITPDELAEADGFIFGFP 80
Migut.M01245     MATKVYIVYYSTYGHVERLAKEIKKGAESVEGVEAKLWQVPETLSDEILSKMGGPPK-SDVPIITPNELVEADGILFGFP 79

Potri.011G033100 TRFGMMAAQFKAFLDATGGLWGTQQLAGKPAGIFFSTASQGGGQETTALTAITQLVHHGMIFVPIGYTFGAGMFEMEQVK 159
Migut.M01839     TRFGMMAAQFKAFFDATGGLWRAQQLAGKPAGIFYSTGSQGGGQETTALTAITQLVHHGMIFVPIGYTFGAGMFEMEKVK 159
AT5G54500        TRFGMMAAQFKAFLDATGGLWRAQALAGKPAGIFYSTGSQGGGQETTALTAITQLVHHGMLFVPIGYTFGAGMFEMENVK 159
AT4G27270        TRFGMMAAQFKAFLDATGGLWRTQQLAGKPAGIFYSTGSQGGGQETTALTAITQLVHHGMIFVPIGYTFGAGMFEMENVK 159
Potri.001G410700 TRFGMMAAQFKAFLDATGGLWKTQQLAGKPAGIFFSTGSQGGGQETTALTAITQLVHHGMIFVPIGYTFGAGMFEMEKVK 159
Potri.011G129400 TRFGMMAAQFKAFLDATGGLWKTQQLAGKPAGMFFSTGSQGGGQETTALTAITQLVHHGMIFVPIGYTFGAGMFEMEKVK 159
TvQR2            TRFGMMAAQFKAFFDSTGGLWKTQALAGKPAGIFFSTGTQGGGQETTALTAITQLTHHGMIYVPIGYTFGADMFNMEKIK 159
ShQR2            TRFGMMAAQFKAFFDSTGGLWRTQALAGKPAGIFYSTGSQGGGQETTPLTAITQLTHHGMIFVPIGYTFGAGMFEMGEIK 160
PaQR2            TRFGMMASQFKAFFDSTGGLWRTQALAGKPAGIFYSTGSQGGGQETTALTAITQLTHHGMIFVPIGYTFGAGMFEMEKIK 160
Migut.M01245     TRFGMMAAQFKAFIDSTGGLWRTQALAGKPAGIFYSTGSQGGGQETTPLTAITQLTHHGMIFVPIGYTFGAGMFEMGEIK 159

Potri.011G033100 GGSPYGAGTFAG-DGTRQPTELELGQAFHQGKYFAGIAKKFKGTT-- 203
Migut.M01839     GGSPYGAGTFAG-DGLRQPSDLELEQAFHQGKYIATITKKLKGSA-- 203
AT5G54500        GGSPYGAGTFAG-DGSRQPTELELQQAFHQGQYIASITKKLKGSTA- 204
AT4G27270        GGSPYGAGTFAG-DGSRQPTELELGQAFHQGKYIAAISKKLKGPAAA 205
Potri.001G410700 GGSPYGAGTFAG-DGSRQPTELELEQAFHQGKYIAAITKKLKGAA-- 203
Potri.011G129400 GGSPYGAGTFAG-DGSRQPTELELEQAFHQGKHIAAITKKLKGAA-- 203
TvQR2            GGSPYGAGTFAGADGSRQPSDIELKQAFHQGMYIAGITKKIKQTSA- 205
ShQR2            GGSPYGAGTYAG-DGSRQPSGIELEQAFHQGKHIAAITKKLKQTT-- 204
PaQR2            GGSPYGAGTYAG-DGSRQPSDIELAQAFHQGKYIAAITKKLKKSV-- 204
Migut.M01245     GGSPYGAGTYAG-DGSRQPSDLELAQAFHQGKYIAGITKKLKTSAA- 204
```

**Supplemental Figure 6. QR2 sequence alignment.**

QR2 sequences from the three parasitic plants and representative non-parasitic plants from Phytozome are shown.

**Supplementary Dataset 1.** PLAS-assembled or 1KP-derived and manually curated transcript sequences of phylloquinone biosynthetic and coexpressed genes described in the manuscript.

```
>PaICS Phelipanche aegyptiaca Pa.c23458_g1_i4.7758
GTTACTACTACTCACATTCCCATAACATATGCAACCCAAATAGAAAAGAAAAAAGATATACACAACTCACACTAGCGAGCGTACAT
GGCTACTACTCTTAAGCAATACTATTTCAATGCCGGCAGTTATAAGGACATTGGATCCAAGAAACGCTTTCCCTCATCCCCCACTG
CAATTGCCAACCCGTCGGTTAATTTCTACAACCATAAACATCAAGAACTCGGTTCGTTGTCCATGAATGGTTGTGGAGGTGATCCA
AGAGCTCCCATTGGCACCATCGAGACTCGGACACTTCCGACAGAACCGACAATGGCACTGGCGGCCGACCGCCTCAACTCCGCCAT
ATATGGCCTCAAATCCGATGCTCCGTCGTTCGACTCAGGGATTATTCGAATCGAGGTACCAATTGGAGAGCAGATAGAGGCCCTTG
ATTGGCTTCGTTCACAGAGCCATAGCCATCTTCTTCCCCGCTGCTTTTTCTCAGGCCGAGATTCCAACAATATCACACCACAAATT
AATGGAAACGGAAATGGGGTTATTAACGGCTATCACTCGTCTTCTTCTCAACAAAAACAGAAGCTTGTTAGTGTTGCCGGCCTGGG
CTCAGCCGTCTCCTTCCGCCATCTTCATCCTTTCTCTTTGGACGATTGGCATTCCATCAAAAGGTTCTTATCCAAAAGGTGTCCAT
TGATTCGTGCTTATGGCGCTATGCGATTTGACGCGAGATCCGTTATAGCCCCTGAGTGGAGGGGCTTTGGTTCATTTTATTTTATG
GTTCCTCAGGTTGAGTTTGATGAGTTTGAAGGAAGTTCAATGATAGCTGTAAATGTTGCATGGGACAATCGCCTGTCACAGTCCTA
CGAACAAGCAATTGCTGCGGTTCAAGCTACAATGTCTAAGATTTCAAGAGTTGTCCGGAGAACAAATGGTAGCTCACCTCGTGCAG
TTCTACTGCATCAGACTCATGTTCCCAACAAAGCGTCATGGGATTCATCTGTCAAGCAAGCTTTGGACTTGATAAGCAGGAAAGAC
TCGAGGCTTATTAAGGTTGTTCTAGCACGTAGCAGCAGAGCACTAACCACCGTTGAGATCGACCCTTTAGAGTGGTTATCAAGCTT
GCAGGTTGAAGGGGTTAATTCTTACCAGTTTTGTCTTCAGCCACCTGAATCCCCTTCATTCATCGGGAACACTCCTGAGCGACTAT
TCTACCGAGACCGACTTAGCGTCTGCAGTGAGGCTTTGGCTGGAACACGAGCTAGAGGAGGAACCGAGTCACTTGATCTGCAGATA
GGAAATGATTTACTTTCAAGTGCTAAAGACCATCATGAATTTGCTGTTGTAAGAGAGAGCATAAGAAGAAAGTTAGAGAATGTATG
CTCTAGCACAATAGTTGAACCAAGCAAAACTCTACGAAAACTCCCACGTGTTCAACATCTTTATGCTAAGCTGACGGGGACATTGC
AGAAAGAAGAAGACGAGTTTAAGATTCTGTCTTCTCTTCATCCGACTCCTGCTGTTTGTGGGCTACCTACGGAAGATGCACGGATT
TTAATTTCCGAAACCGAAAGGTTTGACCGAGGAATGTATGCTGGTCCTGTTGGGTGGTTTGGTGGTGCAGAGAGTGAGTTTGCTGT
TGGGATAAGATCGGCATTAGTTGGGAAGGGTGTTGGTGCATTACTGTATGCTGGAACTGGGATAGTAGAAGGAAGCGACTCGGGCC
TCGAATGGCAGGAGCTCGAACTCAAGACTTCACAGTTCACCAGATTGATGAAACTCGAGGCAACACCTCTTCCGGCAATGGGGAAG
AAAAATTGAAACGTGATGTAAAATAAGGGCTTAAAAACCGGGCGAGCTTACTTGCGAAAATCTCAATGTCCAGTTGCTTCATATGC
TGTCCGGTGCACTGTCTACCATCATCAATATATATTTCTAAGAAATAAAATTAATACTACTACCATGTAATTAGGCATAAGGGAAA
ATAGGTCCTCAATCTCAATGTAAAATATCTGTGTTTTGTACAAAAATATTATTTGTTCAGTAAAGGTTTTCTTCTTGAGCAATAGC
CAATACGGTTTACGTTGTTGGTTGAACGTTTAAGGATTAAATTAAAGTTTTGTGTGATAGGATTAAATTTTGGCTTTAAAATTTCT
GTGGCTCAGTATGGGAAATTGACCAATAAATATAAATTGTTTCCCACCAAAAAAAAAAA
>ShICS Striga hermonthica Sh.c17992_g1_i8.612
CCACACACACACACACACACACACATTAAAGCACAAATGTCCATTATAAGGCATTTCAATGCAAGCTTCAAGGACATTAAGTCCAAGA
AGTGCTTTTTCTCAACCCCAACTGCAATTGCAAATGCATCACTTCAATTCTCCAATTATAAACCTCAAGAGCTTTGTTCCTTGTCA
TTGGATGGGTGTCGAGGCCTCGACCCGCGGGCCCCACTTGGGACGATTGAGACTCGAACGCTCCCCAAGGCCCCGACTCCCGCTCT
GGCGGCCGACCGACTCAACTCTGCAGTCTATCGTCTCAAATCCGACGAGCCGCCTTCGGAATCCGGGATTATTCGGATTGAGGTAC
CCATCGCAGAGCCGATAGAGGCGCTTGACTGGCTTCGTTCGCAGAGCCACGGCCATCTTCTTCCTCGCTGCTTTTTCTCCGGCAGA
GATACGGCCAACACCTCACTAATCGAGCATGTTAACGGCAACGGCAATGGGATTAACGGCCACTCATCTCCTCAACGGAAACTTGT
CAGTGTTGCTGGCCTCGGCTCAGCCGTCTCCTTCCGCCATCTCCGACCGTTCTCTTTGGACGATTGGCATTCGATTAAAAGGTTTG
TATCCAAAAACTGTCCGTTGATTCGTGCTTATGGTGCAATGCGCTTTGATGCGAGATCTAAGATATCCCCTGAGTGGAATGGTTTT
GGTTCCTTCTACTTTATGGTCCCTCAGATCGAGTTTGACGAGTTTGAAGGAAGTTCAATGATAGCTGCAACTGTGGCGTGGGATAA
TCGTCTTTCACTTCCCTACGAACAAGCAGTTGCAACACTCGAAGCTACACTGTCCGAGGTTTCAACAGTCGTTTGGAAAATAAATG
ATAGCTCTCATCGTGCTCCTGTCCTTCACCAGACTCATGTTCCCAACAAAGCCTCGTGGGATCAAGCTGTTAAACGAGCTCTTGAC
TTGATAAGCAGTAAAAACTCCTCGCTTGTTAAGGTTGTGCTTGCGCGTAGCAGCAGAATTCTGACCACGGTTGAGATTGATCCTTT
AGAATGGTTATCGAGCTTGAAGGTTGAAGGGGCTAACTCGTACCAATTTGTTCTTCAGCCACCTGAATCCCCTGCATTCATTGGGA
ACACACCTGAGAGACTGTTTTACCGAGACCGTCTAAGCGTATACAGTGAGGCCTTGGCCGGAACTCGAGCTAGAGGAGGAACAGAA
TCACTAGATTTACAAATAGGACATGATCTACTCACCAGCCCTAAAGATCACCATGAATTCGCTGTCGTAAGAGAGAGTATAAGAAG
AAAATTCGAGAACGTGTGCACAAGCACTGTAGTTGAACCGAGCAAAGCTCTAAGGAAACTGCCACGTGTCCAACATCTTTATGCTA
AGCTAACGGGCACATTGCAGAAAGAAGATGACGAGTTTAAGATTCTGTCTTCACTTCATCCAACCCCTGCTGTTTGTGGGCTTCCA
ACTGAAGATGCACGAATTCTAATATCACAAACTGAAATGTTTGACCGAGGAATGTATGCTGGCCCGGTTGGATGGTTTGGTGGTGC
TGAGAGTGAATTTGCCGTCGGAATAAGATCAGCATTAGTAGGAAAGGATATCGGGGCATTACTTTATGCTGGGACCGGAATAGTAG
AAGGAAGCAACTCTTCCCTCGAGTGGAAGGAACTCGAACTCAAGACATCTCAGTTTACGAAATTGATGAAACTCGAGGGGCCTCTA
CCAACAATAAGGGAGAAAGTTGAAATGTAACTCAGAAAAGAAAAGTATATACTTTTCTTCCAGAAGTATATATAGATTTGCTCAGA
GTGCAGCTTGTTTAAGTGATGCAGACCCTAGTAAGATATTGTCCGCTTTGGGCTACTTCCGCCACCCGCACGGCTTTGTTCTTGGC
TGGGCTTCAATACACACCCACCAAAACGCGTCTTACTAGTGAAGGTATCCACATCCTCATATAAGGCCTGCTTCGTTTCCTTTCAA
ACCCGATGTGGGATGCGCTGCATCATCCACCCCCCCTTGTGGGCCCAGCGGTCACCGCTGGCACATGGGCTGGCTCTGATACCAAC
>TvICS Triphysaria versicolor Tv.c7349_g1_i2.6356
CTCAAATATTATAGTACAAAACAAGAGAAGAAAATAATCACACACTAGACTACACATGTCTACTACTAAGTATTTCAATGCTGGTT
TCAAAGACATTGAGTCCAAGAAATTCATTTTCTCATCCCCGATCGCAATTGCAAAACCATCACTTCATTTCTCTAATCATAAATAT
GAACTTGGTTCATTGTCCATGAATGGATGTGGAGGCGATCCACGAGCCCCCATTGAAAAAATCGAGACTCGAACGCTTCCGATTGC
TCCGACGCCGGCATTGGCGGCTGACTGCCTCAACTCCGCCATCTACCACCTCAAGTCGAATGCTCCGGCGGCCGATTCCGGGATTA
TTCGGATTGAGGTACCAATCCAGGAGCAGATAGAGGCGCTTGATTGGCTTCGTTCACAAAGTCACACCCATCTTCTTCCTCGCTGT
TTCTTCTCGGGCCGAGATTCCAACAATATCCCACCACTCAATCAACACATCAATGGAAATGGAAATGGGAATGGGATTAATGGCCA
```

TTCTTCTGCTAAAGAACATCAACAGAAGCTTGTCGGTGTTGCCGGGCTGGGCTCAGCTGTCTCCTTTAGCCATCTCCACCCTTTCT
CTTTGGACGATTGGCATTCCATCAAAAGGTTTTTATCCAAAAATTGTCCATTGATTCGTGCTTATGGCTCAATTCGGTTCGACGCG
AGATCCAATATAGCCTCCGAGTGGAATGGTTTTGGATCCTTTTATTTTATGGTCCCTCAGATTGAGTTCGATGAATTTGAAGGAAG
TTCAATGATAGCTGCAACAGTTGCATGGGACAATCGTTTATCACGGTCCTACGAACAGGCAATGGCTGCACTCGAGGCTACAATGT
CTAAGGTATCATCGGTTATCCGGAGATTAAATGGTAGCTCACATCAGGCTGTTGTGCTCCACCAGACTCATGTTCCCAACAAAGCC
TCCTGGGATTTGGCTGTTAAACGAGCTTTGGATTCGATAAGAAGTAAAAACTCACCGCTTGTTAAGGTTGTTCTTGCACGTAGCAG
CAGAAATTTGACCACGGTCGAGATTGACCCTTTAGAGTGGTTGTCGAGATTGCAGGTCGAAGGGGTTAATGCCTACCAGTTTTGTC
TTCAACCACCTGAATCCCCTGCATTCATCGGAAACACTCCAGAGCGACTATTCTATCGAGACCGACTAAGTATATGGAGTGAGGCT
TTAGCCGCAACACGAGGTAGAGGTGCAACCGAGTCACTCGATTTACAGATCGGAAACGATTTACTTTCAAGTGCTAAAGACCATCA
CGAATTTGCTGTTGTACGAGAGAGTATAAGGAGAAAATTTGAGAAGGTGTGCTCAAGCACAGTAGTAGAACCGAGCAAAGCTTTAA
GAAAACTACCACGTGTTCAACACCTTTATGCTAAGCTCTCGGGCACATTACATAAAGAAGATGACGAGTTTAAGATTTTGTCTTCT
CTTCATCCAACTCCTGCTGTTTGTGGGCTTCCTACAGAAGATGCGCGGGTTTTAATATCGGAAACTGAAATGTTTGACCGAGGAAT
GTATGCTGGCCCGGTTGGTTGGTTTGGTGGTGCCGAGAGTGAGTTTGCTGTCGGAATAAGATCCGCATTAGTTGGGAAGGGTGTTG
GCGCGTTACTGTATGCGGGAACGGGAATAGTAGAAGGAAGCAATTCGTCCCTGGAATGGCAAGAACTCGAACTTAAGACATCACAG
TTTGCAAAATTGATGAAACTTGAGGCTCCTCTTCAAGTGACGAGGGGAGAAAAATGATCTGTGACTTAAGGTTAGTATAAGCAATG
AGTTACTCATCTGCATTTGTGTAGCTTATTCTACAGTGCCAAAACCCAATATATATATTTCTCACTTGTGATGAAAAGTCAGATGT
CAATTCTTGTATACAGCTGGTTTGTAATCATCCATTGTTAGATAAACTCTCTATATTGTATATATAGAGAGAGATGGGCCTGAGGA
AACAATATTTTTTATTATCATTCTTTTTTTTTTTCGTTACCAATGCTTTGTACATAATTTTGTTTGGTTCCTATCCTGTTGTGATA
TAATATGTAGAAATTATAAAGGGGACTCACG

>PaPHYLLO Phelipanche aegyptiaca Pa.c19994_g1_i1.144
GCGAGATAAGTATGTATTATATATTGAACCAGCAACTGTCATTTCTTTAGGACTTAGCGCCTTCTCCTCTTTGAACCTCCATTACC
AATGATTCCCTTCACCTTATACTCTCAAATTACTCCACCTTTTCCATTATCTCCAATCCAAACCCACCAAAAGACCATAATAACAA
TTACTAAGCGCGCCCATTTTCTGAAACCTGCTTTGCCATGTCTAGCTCGCTTTGGCCACCGTCAAAATCCCATTTCCAAGGTTGTG
AGTAGCTCAATGGGAAAAGATAAAGTGTTGGATGCTAAACATGCTGCATTGCTGATCAATACTTGCATTACGCGTAATTTGCCGCC
GGTTTTGAGTTTAGAGCAAGGACTGGATAGGATTAAGGAGGATTTGGAGGAGTTAAAGGCTAATCGTCCTCCTTGTTCAAGTGGGA
TGTACAGATTCCAGCTTGCAGTGCCTCCAAGTGCAAAAGCGTTGAACTGGTTTTGCTCTCAACCGGAGTCATCAAACGTCTTTCCT
CTATGCTTCCTCTCCAATGAGGACAATCCAACATATAATTCACTTTCTCTTGGAAGAACCAGGGGTGTTTTTGGAATTGGTTCTGC
TGTTGTTTTCAAGGATCGTTCTCATGCTTCAGAAAATAGCAGAGCTGTACGAAGATGTCTTTCGGCTGAACCAACATCTTCTAAGG
CTTATGGTTTCTTGGATATCGAGTTTGACAGTAACATGTCTACTACAAAGCATCAGATCGGTTCATATTACCTTTTCATTCCCCAG
ATTGAGTTGGATGAATTTGAAGATATCCCTTTCTTGATTGCAACATTGGCATGGGATGACTCTTCAATTTGTACATTTAGTGAAGC
TGTTCAAAGATTTGAGCTCGCTTTTGATCAGGCCAGATACACCTGTGGAAACGGTAGCCAATTGATTAGATCTTCTCTTTTAAAGT
TCAGCAATGCAGAGAAACATACGGAAATGGTTCGTGCAAATGGTCTACTATTGGATGGGCAACATCTTACAGCCAGCACCATGGAA
ACGGAAGATGCTTCGTCTTGTTGCAGACAGTTTGTTGCAAGACTTTCATCAACTTTATCGATTTCAAATAACATGCTCCCAAGAGA
CGAAACCAAATTAATCGAAAATGTCACTCAAGATTTTCCCAATATCAATACTTTGTGGGCATACCTTATAGTTGAAGAATGCACCC
GTCTTGGTTTGACATATTTCTGTATAGCTCCCGGATCAAGGTCATCTCCACTGACGATTGCTGCAGCCAGTCACCCCCTTACGACT
TGCATTGCATGCATTGATGAACGATCCCTAGCATTTCATGCTCTCGGTTATGCCAAAGGTTCCGGAAAACCAGCAGTTGTTATAAC
ATCATCAGGGACAGCTGTCTCAAATCTTTTTCCGGCCGTTGTGGAGGCTAGCCAGAGTTTCGTACCAATGCTGTTGCTTACTGCCG
ACCGTCCTCTTGAGCTCGTAGATGTTGGGGCGAACCAAGCAATCAATCAGGTCAATCATTACGGATCATTCGTGAGGCACTTTTTC
AGCCTTCCTCCACCTGCCGATGACATATCCGCAAAATTCGTTCTTACCACAATCGACTCAGCAGTATGCAAATCAACATCTTCACC
AAACGGTCCTATACATATTAACTGCCCCTTTCGAGAACCGCTAGCACACAGTCCAAGAAATTGGGACCTTAAATGTTTAAGTGGAT
TAGACGTTTGGATTTCAAATGCCAAGCCGTTTACCAGTTACATTCCATTACAACATTCCCTAACGTGTAATAACCCGAACGGGAAT
ATGATTGAAGTGTTGAAACTAGTCCAAGGGGCCAATAATGGGATTTTAGTTTTGGGTTCGATTCACAAAGAGGATGATATGTGGGC
AGGTCTTTTGTTGGCTAAGCACTTGTCGTGGCCGGTTGTTGTTGATATCCAGTCGGGTTTAAGATTGAGGAAGCACTTGTCGTCGT
TTCATGAGAGAAAAGATATATTGTTCATTGATCAGCTTGATCAACTGTTGATGTCGGATTCTGTTAGAGATTGGATGCAAGCTGAT
GTTATAATACAGATTGGAAGTCGGATAACGGGAAGGCGCATTTCTCAGATGATAGAGCAATGCACTCCGTGTGCGTACATCATGGT
CGACGATCATCCAGGTCGTCATGATCCTTCAAATATCGTGACACATAGGATACAAAGTACCATCAGTGAATTCAGTGATTGCTTGA
TTAAATGTTCCATCTCTCACGTAAGCACGAAAGGGTCGGAATTTATACGAGGACTGGATATGATGGCTGCCTGGGAAACATCTTTT
TTGATTAATTCCGAACAATCGTTGACTGAACCTTATGTCGCGCGAAAAATCTTTGAGACGATTCGCTGTGGATCCGCTTTGTTTTA
TGGAAATAGCATGCCAATACGTGATGCGGACATGTATGGGAGTAACAGGGTGCAGTGCACCCACAGTGCTTCTCTGATGTTGAGTT
CTGGTTTACCGTGTCATCCTGTGCATGTTACCGGAAATAGAGGTGCTAGTGGTATTGATGGTTTGATTAGCACAGCTATTGGGTTT
GCAGTTGGCTGCAATAAAAGAGTGCTTCTTGTAATCGGAGATATTTCATCTCTGCATGATACCAATGGGTTGGCACTACTTAAACA
ACGGACCTTCCGGAAACCGATGGTCATACTTGTGCTTAACAATCATGGTGGTGCTATCTTTAGTCAACTACCTGTTGCAAATACGA
CAGACAGAAGTATACTCGACCAATTTTTCTACACGTCTCACAATGTTTCAATACAAAACCTATGTCTGGCACATGGTGTGAAGCAT
GTACAAGTGCGAACAAAAAAGGAGTTGCAAGACGCATTGTTCACATCTCAAAGGGAAGACGTTGATTGTGTAGTGGAAGTTGAGAG
TGAAATTGACACCAATGTTGCTATTCATAGTAACTTGAGAAATTTTACTCGCAAAGCCTCGGATCATGCTTTCAACATCCTCTCGA
AGCTGTCAGTTGCAGATTCCAACTCGCAACATTACAAAATAAATAAAATGGATTACTCTATGTACCGGGTTCAGCTAAATGCTCCA
CCTACATCAGCTCTTACGGACTCCAAGACTAGCACATCCTACAGAGAAGGTTTTGTTATAAGTCTGTCTCTTGAAGATGGTAGTAT
TGGGTTTGGCGAGGTCGCCCCTCTTGAAATCCACAATGAAAATTTGCTTGATGTGGAAGAGCAACTTCGGTTTCTTATTCATGCCA
TAGAAGGACAAACAATCAATAACGCCCTAGCTCTGTTGAATTGTTCATTTTCTTCTTGGATATGGAACAGTTTAGGAATTCCGCCA
GGTTCAGTCTTTCCCAGTGTCAGATGCGGATTAGAGACGGCTCTTCTCAGTGCACTTGCAAGTAGACAAAGTAGCACTTTATTAAA
TATACTTAATCCCGCAAGCGAAAAATCGTCCAAGGAATCATCTGCCATTCAAATTTGCGCCCTAATTGACTCCTATGGAAGTCCAA
TGGACACAGCTTTCGTAGCATCCAATCTTGTTGCCGAAGGATTTAGAGCAATAAAATTAAAGGTTGCGCGTCGAGCTGATACCAAT

GAAGATATTGCTACTATACAAGAGGTGAGAAGGAAAGTGGGAAAAGATATTGTTCTCCGTGCAGATGCAAATAGGAAATGGAATTA
TGATGAAGCTGTCAAGTTTGCTCACTCGGTCAAAGATTGTGGCTTGCAATATATCGAGGAGCCTGTTAATAACGAGTATGATATAG
TGAAATTTTGCCAAGAAACTGGCCTACCAGTGGCATTGGATGAAACAATCAACTCGATCGGGGAGAATCCTCTTGAGGTCCTTCAG
AAATACAGCCACTCAGGAATAACTGCTGTTGTAATCAAACCAAGTGTTATCGGAGGTTTCGAAAGAGCAGCATTGATTGCAAGATG
GGCCCAACAGCACGGAAAGACGGTTGTAATTAGTGCTGCATTTGAAAGTTCACTCGGTTTGTCGGCCTATATCCAATTTGCTCATT
ACCTCGACCTGCAAAACGCCGAGATACAGAATTTGATGAATAAAGAACCTGCACCAGTCACAGCACACGGTTTCGGAACTTACAAA
TGGTTTAAGGAAGACGTGACTGCAGAGAATCTAAATATCCGTTATGATTCAGACTGTGATTTTGTTAAGGCTGATGCTGTTGATGC
CGGTCGATTTCTCCAGAATTGTCGGTTAAATCCTGATAAGGTTGTTAGAGTTTTTAATCAAGAAAAAGTGCGGGAATATCAGTTGG
CAGTTGATACAGAGGGCGTCTCATTTACTACAAATGTGCTAGAGATGGGAGAAAGCATTGATGGTACTGCAGTTGTGTTTCTTCAT
GGATTTCTTGGAACTGGAGAAGATTGGATCCCGACCATGAAAGCCATCTCAAGCTCAACCAGATGCCTTGCAATAGATCTTCCTGG
TCATGGTGGATCAAAGTTGCAATATAAGGGTACCAAGGGTTCAGATCAATCCGATTTATCGATTGATGTGGTGGTTGATATCTTAT
GCAAGGTGTTGAATAATATTACTACTCAGAAGGTTATACTCGTCGGGTACTCGATGGGAGCTAGGATATCTTTATACACAACACTA
AAACGCAGTGATAAGGTTGAAAGAGCAGTGATTATATCTGGAAGCCCGGGTTTAGTTGATAACGGTGCAAGAGAGATCCGTAGAGC
TAAAGATGACTTCAGAGCTAGCACACTCATGTCAAACGGTTTAGAATTTTTTACGGAGGCTTGGTATGCTGAAGAACTCTGGGCTA
GCTTAAAAACCCATCCACACTTCAAACAAATAGTTGCCAGTCGTTTGCAGCATGACGACTTGCATACTCTTGGCAAAGTTCTGTCT
GACTTAAGCATCGGAAGGCAGCCATCACTGTGGGAAGATCTGAAGCACTGCAAGGTGCCCCTCCAGATTATAGTGGGAGAAAAGGA
TGTCAAGTTCAAGAAAATCGCTCATGAGATGTATACAAAGATTGAACACGGAAACGGAAGTAGTAATTCACTGCCAGTTGCTGAAA
TTTTGAATGCTGGACATGCTGTTCATCTTGAGAATCCTCTTGGTCTTATTACTGCTTTAAGGCAGTTCATAAAGAGAGAAAAGAAT
ACCTAGTTACTTTTGTTTTTGTTTTTTTATTTTTTAAAGAAAAAAATAATACACAAGGAGTTGCATTGTTGTCCACAGACAGAAAA
ATAAAGGTGTGTTCATTTTTAGGACGAAAACAGTTCTGTTAACTGAATAGAGATTGGATTTGATTTATTCATTTGTTATGAATATT
AAAGGAGGCTTATGTAATTTTGAAATTTCTCTGGTTTTCAGGGTACTTTTAGAAGAAAAAAAAAAAAAAAAA
>ShPHYLLO Striga hermonthica Sh.c19093_g2_i7.6567
CCGAAACCCCAGCAACTGTCATAGTTCTTCTCTGGGACTTAGCGCCCTCCTACTCTTTGAACCTCCATTTAATTTATCAACAAAAT
GAGCTCCTTCACCTTATTCTACTCTCAATCTACGCTCCCTTTTCCACTTTCTCCACTCCAACCCAAACAGAAGACCATAATCACAG
TAACAAAGCGCGCTTTTCTAATGAAACCTGCTTTGCGATGTCTAACTCGCTTTAGCCACCACCAAAATCCCATTTCCCAGGTTTTG
AGCAGTTCATCGGAAAAGAATGAAGTCTTGGATGCTAAAGATGCGGCATTGCTGGTCAATACTTGCATTACGCGTAATTTGCCGCC
GGTTTTGAGTTTGGAACAAGGGCTGGAGAGGATTAAGGGGTCCGTGGAGGAATTGAGGGTTAATCCTCCTTGTTGCTCAAGTGGGA
TGTACAGATTTCAGCTGGCAGTGCCACCAAGTGCAAAAGCATTGAACTGGTTTTGCTCCCAACCGGATAAGTCAAACATTTTTCCT
CTATTTTTCCTTTCCAATGAGGAGAATCCGACATATAATTCGCTCTCTCTTGGAAGAACCAGGGGTGTTTTCGGTATTGGTTCAGC
CGTTGTCTTCAAGGATTGTTCTCCTCATAGTGCAGGAAAAGGCAGTGATATTAGAAGACATCTTTTGGCTGAACCAACAAGTGCAA
AGGTTTATGGTTTCTTGGATATTGAGTTCGTCACCGACATGTCTGCTATAAAGCATCAGAGTGGTTCGTATTACCTTTTCATCCCT
CAGATTGAGTTGGATGAGTTTGAAGATATCCCTTCCTTGACCGCGACATTGGCATGGGACGACTCTTCAATTTGTACCTTTAGTGA
AGCTGTTCGAAGATTTGAGCTTGCATTAGACCAGGCCATGCATACCTGTGCAAATAGAAGCCAACTTATTCGGTCTTCTCTTTCAA
AGTATCGCAATGCAGAGAAACATAAGGCAATGGTACGTGCAAATGCTTTGTTATTGGATGGGAAGCACACAGCAGCCAGCGCCCTG
GGGCTGGGAGATTCTTTGTCTTGTTGCAGTCAGTTTGTTGCTAGGCTTTCATCAACTTTATCAATTGCAGATAACATGATAGATGA
AACAAAATCAGTCAACAATGTGATTCAAGATTTCCCCAACATCAATGCTTTGTGGGCGTACCTTATAGTTGAGGAATGCACTCGAC
TTGGTTTGACGTATTTTTGTGTGGCCCCTGGATCAAGGTCATCTCCCTTGACAATTGCTGCCGCCAGTCATCCTCTTACCACTTGT
GTTGCATGCATTGATGAAAGATCTCTTGCATTTCATGCGCTCGGTTATGCAAAAGGTTCAGGAAAACCGGCAGTTGTCATAACATC
ATCAGGCACAGCTGTCACGAATCTTTTTCCGGCTGTGGTTGAGGCTTGCCAGAGTTTTGTACCAATGCTGTTGCTAACGGCTGATC
GCCCTCCCGAGCTTGTAGATGTTGGGGCTAACCAAGCTATTGATCAGGTGAAGCATTATGGGTCGTTCGTGAGGCACTTTTTCAGT
CTCCCTCCTCCTACAGACGAAATATCGGCAAAATACGTACTAACTACAGTAGACTCAGCTGTTGTTAAATCAACATCATCCCCAGT
GGGCCCTGTACACATCAACTGCCCTTTCCGGGAGCCACTTGCAAGTACCTCAAAAACTTGGAGTCCCAAGTGTTTAAGTGGATTGG
ACATTTGGATTTCAAATGCCGAGCCATTAACCAGCCATTCTTTGACTTGTTATAGTAACGTGAGAGAGCAGATGGTTGAGGTTGTT
AGGGTGGTCCAAAGGGCTAATCATGGGATTTTGGTGTTGGGTTCTATTCAGAAGGAAGACGATATGTGGGCAGCTCTTTTGCTGGC
CAAGCACTTGTTGTGGCCGGTTGCTGTTGATATCCAGTCGGGTTTGAGGTTGAGGAAATACTTGTCGTCTTTTCTTGACAGCAAGG
ATATATTGTTCGTTGATCAGCTCGACCAGCTGTTGCTGTCAGAGTCTGTAAGGGAATGGATGCGGGCAGATGTAATAATACAGGTT
GGGAGTCGGATAACAGGGAGACGCATTGCCCAGATGATGGATCATTGTTCACCTTGCTCCTACATCATGGTCGATGATCATCCAGC
TCGTCATGATCCTTCTAATATCATCACACACAGGATCCAAAGTACCATCCCCGAGTTCACTGATTACTTGATCAAGTGTTGCAGCC
CTCGTGCGAGCAACAAATGGCGGGAGTTAATACGAGGATTGGACATGAGGGCTTCCTGGGAGACCTCATTTCTGATTAATTCGGAA
CAATCCTTGACCGAACCTTACGTAGCACGGAAAATCTTCGAGGCAATCCGCTGTGGGTCAGCTTTGTTTTACGGGAACAGCATGCC
TATACGTGATGCGGACATGTACGGTAGTGACTGGGTGCATTGCACCCACAGTGCTGCTCTCATGTTGAGCTCTGGCTTGCCATGTC
ACCCGGTGCATGTTAGCGGGAATAGAGGGGCTAGCGGTATTGATGGGTTGATAAGCACTGCTATTGGATTTGCTGTTGGCTGCAAT
AAAAGAGTACTTCTTGTGATTGGCGATATTTCATTTTTGCATGATACGAACGGGTTGGCATTACTAAGACAATGGACATACCGGAA
ACCAATGGTCATACTTGTGCTTAACAATCATGGTGGTGCTATCTTTAGTCAACTACCAGTTGCAAATACCACAGATAGAAGCATAC
TAGACCAGTTTTTCTACACGTCTCACAATGTTTCGATACGCAATCTATGTTTAGCACATGGTGTGAAGCATGTTCAAGTGCAAACA
AAAAGGGAGTTAGAAGATGCATTGTCCACATCTCAAAAAGAAGACATTGATTGTGTGGTGGAAGTCGAGAGTGAAATTGATATGAA
CGTTGCTATTCATAGTAAGTTGAGGAACTTTTCTCGGAAAGCCTCGGATAATGCTCTTAACATCCTCTCAAAGCTGTCAGTTGAAG
ATTCCCACTCGCAAGATTACAAGATCCATAAAATGGATTACTCTCTCTACCGGGTTGAGCTTAATGCCCCACCTACTTCAGCTTCA
ATGGAGTCGAAGGCTCCCACATCCTATAGGGAGGGTTTTGTCATAAGTCTAGCTCTTGAAGATGGTAGTACTGGTTTTGGTGAGGT
GGCACCTCTTGAAATCCACAAAGAGAACTTGCTTGATGTGGAAGAGCAACTTCGGTTTCTTGTTCACGCTATAGAAGGAAAGACAA
TCAGTGGCATACTACCTCTGTTGAGTTGTTCAATTTCTTCTTGGATATGGAAGAATTTAGGAATTCCGCCAGGTTCAATCTTTCCC
AGTGTTCGATGTGGATTAGAGACAGCTCTCCTGAGTGCTATTGCAAGTAGACAAAATAGCACTTTACTGGATATAATCAGCCCCAC
AACCGAAAAAACGTCCGAGAAATCATCCCCTGTTCAAATTTGTGCCCTCATTGACTCCTACGGGACTCCAATGGACACAGCTCTTG

TAGCATCCAAGCTGGTTGCTGAAGGATTTACGGCTATAAAAATAAAAGTTGCACGTCGAGCAGATCCTGATGAAGACGTAGTTACA
ATACAAGAGGTGAGAAGGAAAGTGGGAAAAGATATTGTACTCCGTGCGGATGCTAATAGGAAATGGAATTATGATGAAGCCGTTAA
GTTTGCTCTCTCGACCAAAGACTGCTGCCTGCAATATATCGAGGAACCAGTGAATGATGAGAATGATATAGTGAAATTCTGTGAAG
AAACTGGTGTGCCAGTGGCTTTGGATGAAACAATCAACTCTATAGGGGAAAACCCTCTGGAGGTCCTCGGGAAGTACAGCCACTCG
GGAATTACTTCTGTTGTAATCAAACCTAGTGTCATCGGCGGGTTCGAAAAGGCAGCTCTGGTAGCAAGGTGGGCCCATCAGCACGG
GAAAATGGTTGTAGTTAGTGCTGCATACGAGAGTTCACTTGGTTTGTCAGCCTTCATCCAGTTCGCCCGTTTCATCGACCTGCAAA
ATGCCGAAATTCGAAGCTTGACGAGCAAGGAGCCCGGACCAATCACAGTGCACGGATTGGGGACATACAAATGGTTCAAAGAGGAT
GTGACGTTGGAGCATTTAAATATTCATTATAGTCCTGAGCACAGATCAGTCGAGGTCGATGCTGTTGATGCCGGTCGATTTCTCCA
GGATTTGCGGGTAAATAATGACGTAGTTGTCAGAAAGTTTATTGGTGAACAAGTGCGCAAGTACCAAGTAGCGGTTGATACAGACG
GTTTCTCGTTTACTACAAATGTGGTCGAGGCTGGAGAAAGCATTGATGGTAGTGCTGTTGTGTTCCTCCATGGGTTTCTTGGAAGC
GGAGATGATTGGGTCCCGATCATGAAATCCCTCTCAACCTCCACTAGATGCATTGCAATCGATCTACCCGGCCACGCTGAATCAAA
GTTGCAGTTTAACGGTTCCGATTTGTCAATCGATATGGTAGTTCATATCTTGTGCAAGGTGTTAAATAATCTCACCCCTCATAAGG
TCACTCTTGTGGGCTACTCAATGGGAGCTCGGATATCTTTATACACAGCCCTAAAATGTTCTCACAAGGTTGAAAAGGCTGTTATA
ATCTCGGGAAGCCCGGGTTTGATAGACAAGGATGCAAGGGAAATCCGTAAGGCTAAAGATGACTTTCGAGCAAGCACACTGGTGTC
AAATGGGTTACAGTTTTTTACAGAGGCTTGGTATGCTGAAGAACTCTGGGCAAGCTTAAGAACCCATCCACACTTTAAAATGATAG
TTAACAGCCGTTTGCAGCATGATGACTTGCCGACTCTTGGTAAAGTTCTCTCTGGCTTAAGCATCGGAAGGCAACCATCTTTGTGG
GAGGATCTGAAGCATTGCAAGGTCCCACTGCATATTTTAGTGGGAGAAAAAGATGCCAAGTTTAGAAGAATCGCTCATGAAATGTA
CGCCAAACTCGGCCATGAAAATGGAAGTTCTAATTATTCACCGCCAGTGACTGAAATTCCGAATGCTGGACATGCAGTTCATCTCG
AGAATCCTCTTGCTCTGATTACCGCTCTACGACAGTTCATATAGAGAGGGAAGACTAGTTAGATCCCTTGATTTATGTACAATAAA
ATAAAATTAAAAAAAAAAAAGAAAAGA
>TvPHYLLO Triphysaria versicolor Tv.c112198_g1_i1.18287
ATGATATTCATAGTTAAAATTTAGCTCACTCTACTCTTGAACCTCCATTAACAATGAGCCCTTTCACCTTACACTCTCAAATCACA
CTCCCTTTTGCACTATCTCCACTCCAACTCCACCAAAAAACCATTATCACAATAACTAAGCGCACCCATTTCCTCCTCAAACCTCC
TCCGCCCTGTTTAACTCGCTTTAGCTGCCGACAAAATCCCATCTCCAAGGTAGTTAGAAAATCATCAATGGGAAAAGATCAAATCT
TGGATGCTGCATTGCTGGTCAATACTTGCATAAAGCGTAATTTGGGGCCGGTTTTGAGTTTAGAGCAAGGATTGGACCGGATTAAG
GAGGCTGTGGAGGAGTTAAAGGCTAATCATCCTTCTTGTTCAAGTGGGATGTATAGATTTCAGCTTGCAGTGCCTCCGAGTGCGAA
AGCGTTGAACTGGTTTTGTTCTCAACCGGAATTATCAAACGTGTTTCCATTATTCTTTCTTTCTAATGAGGAGAACCCAATATATA
ACTCACTTTCTCTTGGAAGAACGAGGGGCGTTTTTGGTATTGGTTCAGCTGTCGTCTTCGAGGATTGTTCTTCTCATGGTTCGAAG
AAAACCAGTGCTATACGAACATCTCCAAAGGTGTATGGTTTCTTGAATATCGAGCTCGACACAAATATGTCCACTATAAAGCATCA
GAGTGGTTCGACTTACCTTTTCATTCCTCAGATTGAGTTGGATGAATTCGAAGATATCCCTTTCTTGGCTGCCACGTTGGCATGGG
ACGACTCATCGATATGTACTTTTAGTGAAGCTGTTCAAAGATTTGAGCTCGCTTTTGACCAGGCCAGATACACATGTGGAAACGGT
AATCAATTGATACGATCTTCTCTTTTAAAGTTCAGCAATGCGGAGAAACATGGTGAAATGGTACGTGCAAATTCTCTGTTATTCGA
TGGGAAGCATATTGCAGCCAGCACCTTGGAATTGGGAGATGCTTCGTCTTGTTCTCGTCAGTTCGTTGCTAGGCTTTCGTCAACTT
TATCAATAGCAAACAACATGCACCCGAGTGACGAAACCAAATTAGTCAGCAATATGACTCAAGATTTTCCAAATATCAATGCTTTA
TGGGCATACCTTATAGTTGAAGAATGCACTCGACTCGGTTTGACATATTTTTGTGTAGCGCCTGGATCAAGATCATCTCCTTTAAC
TATAGCTGCAGCTAGTCACCCTCTTACGACTTGTATCGCATGTATCGACGAACGATCACTCGCGTTTCATGCTCTCGGTTATGCCA
AAGGCTCCCAAAAGCCAGCTATCATTATAACATCTTCAGGCACAGCCGTCTCAAATCTTTTTCCCGCCGTTGTAGAGGCTAGCCAG
AGTTTTGTACCAATGCTATTGCTAACTGCCGATCGTCCTCCCGAGCTCGTAGATGTTGGGTCAAACCAAGCAATCAATCAGATAAA
TCATTACGGATCATTCGTGAGGCAATTTTTCAGTCTTCCCTCTCCTTCTGACGATATATCCGCAAGATATATTCTTACCACTATCG
ACTCGGCTGTATTTAAAGCAACTTCTTCGCCAACCGGTCCAATACACATAAACTGCCCTTTTAAAGAGCCACTAGCAAACAGTCTA
AAAAATTGGAACCGTAACTGTTTAAACGGACTAGACGTTTGGATTTCGAACGCCAAACCGTTTACGAGCTACATTCCGTTAAAAAA
TGCATTGACGTTTGATATGATTGAAGTGACGAGGCTGGTCCAAGGGGCCGATCGAGGGATTTTAGTGTTGGGTTCGATTCAGAAAG
AGGATGATATGTGGGCTGCTCTTTTATTGGCTAAGCATTTGTTATGGCCAGTTGTTGTCGATATTCAGTCGGGTTTGCGGGTGAGA
AAGTACGTTTCGTCGGTTCTTGACAGCAAGGATATATTGTTCGTTGATCATCTCGATCAGCTGTTATTGTCGGATTCTGTCAAGAA
TTGGATGCGGGCCGATGTTGTAATACAGATTGGGAGTCGGATAACCGGGAGACGAATTTCTCAAATGATAGAGAAATGCACTCCGT
GTCCGTACATCTTAGTCGACGATCATCCGGGCCGTCACGATCCTTCTCATATCATCACACACAGGATACAAAGCACCATCTCCGAG
TTCAGCTTTTGCTTGATCAAATCTTACACACCCGATATAAGCAAGAAATGGACGGATCTTATTCGAGGATTGGACACAATGGTTGC
CTGGGAAACGTCATTTTTGATTAATTCCGAGCAATCATTGACCGAACCTTACGTAGCGCGAAAAATCTCCGAGATGATCCGATGTG
GGTCCGCTTTATTTTATGGGAATAGCATGCCGATACGTGACGGGGACATGTACGCGAGTAACTTGGTGCAATGCACCCATAGCGAT
TCTCTAATGTTGAACTCCGGTTTAGAGTGTCACCCGGTGCATGTTAGTGGAAATAGAGGAGCTAGTGGTATAGATGGTTTGATTAG
CACTGCTATTGGATTTGCTGTCGGCTGTAACAAGAGAGTGCTTCTTGTGATCGGAGATATTTCGTTTCTGCATGATACTAACGGGC
TGGCATTACTGAGACAACGGACATCTCGGAAACCGATGGTCATACTTGTGCTTAACAATCACGGTGGGGCCATATTTAGTCAACTG
CCGGTTGCAAGTACGATAGACAGAAGTATACTCGATCAGTTTTTCTACACGTCTCACAATGTTTCGATACGCGATCTATGTCTCGC
ACATGGTGTGAAGCATATATCAGTTCAAACTAAAAGTGAATTGCAAGACACATTGTTCAAATCTCAAAGAGAAGAAGTTGATTGTG
TAGTGGAAGTCGATAGCGAAATTGATACGAACGTTGCTATTCATAGTACTTTGAGGGATTTTAATCGCAAGGCCTCGGATCAAGCT
TTAAACATCCTCTCAAATCTTTCAGTTTCAGTTGAAGATGACAACTTTCAAGATTACAAGATTAATAAAATGGATTATTCTCTGTA
CCGGGTTCAGCTTAATGCTCCACCTACTTCAGCTTCTTCGACGAATTCCAAGAATAGCGCATCTTATAGAGAAGGCTTTGTTATAA
CTCTGTCTCTTGAAGATGGTAGTATCGGATTTGGCGAGATTGCACCTCTTGAGATTCACAAAGAAAACTTACTTGATGTCGAAGAG
CAACTTCGTTTTCTTGTTCACGCCATACAAGGAAAGACGATCAGTAACATCATACCTTTATTGAAATGCTCGTTTTCTTCTTGGAT
ATGGAACAATTTAGGAATTCCGCCAGGTTCGATTTTTCCAAGTGTACGGTGCGGATTAGAGATGGCTCTCCTCAGTGTAATTGCAA
GTAGACAAAATAGCACATTGTTGGATATAATTAATCCAACAAGCGAAGAATCGTCCCCCGTTCAAATTTGTGCTCTTATCGACTCT
TATGGGAGTCCAACGGAAACAGCTTTTGTTGCCTCCAACCTTGTTGCCGAAGGATTTACGGCTATAAAAATAAAAGTAGCGCGCCG
AGCAAATCCCGATGAAGATATCGCTACAATACAAGAGGTGAGAAGGAAAGTGGGACCAGATATTGTACTCCGTGTCGATGCAAATA

GAAAATGGACATATGAGGAAGCTGTTAAGTTTGCTCGATCCGTCAAAGATTGCTGCTTGCAGTATATTGAGGAGCCTGTTAACAAC
AAGGATGATATAGTGCGATTCTGTAAAGAAACGGGCTTACCAGTTGCATTGGATGAAACAATAAATTCGATCGGAGATAGTATTTC
TCTTGAGGTCCTTCAGAAGTACAGCCACTCTGGAGTAGCTGCTGTTGTAATCAAACCAAGTGTAATTGGAGGTTTCGAAAAGGCGG
CATATGTTGCGAGATGGGCCCACCAGAACGGAAAAACAGCTGTTATTAGTGCTGCATTCGAAAGCTCGCTCGGTTTATCAGCCTAT
ATCCAATTCGCTCGTTACCTCGACCTGCAAAATGCCGAGATACATAGTCTGATGAACAAGGAGCCTGCACCAATCACAGCTCACGG
TTTCGGAACTTACAAGTGGTTTAAAGAAAACGTAACGGATGAGAATCTGAATATCCGTTATGATCCGGACCGTCGGTTTGTCGTTG
CTGATGCTAATGATGCCGGTCGATTTCTCCAGAATTGTCGGATAAATCGAGACTCGATTGTTCGGAGCTTCGTTCATGAAGAATTA
AAGGAATATCAGTTGACAGTTGATACGGATTTTGTCTCGTTTACTACAAATGTGGTCGAGATCGGAGAAAGCACCGATGGTACTAC
AGTTGTGTTTCTTCATGGATTTCTCGGAAACGGCAAAGATTGGGTTCCGATTATGAAGGCAATCTCAAGCTCAACTAGATGCATTG
CAATTGATATTCCTGGTCACGGCGGATCAAAACTGCAATATAAGGGTATCAACGGTTCAGATCAACCCGATTTATCTATTGACTTA
ATAGTTGATATATTATGTAAGCTGTTGGATAATATTACTCCTCGGAAAGTAATACTCGTCGGTTATTCAATGGGAGCCAGAATCTC
ATTATACACAGCCCTGAAACACAGTAATAAGGTTGAAAGTGCTGTGATAATATCTGGAAGTCCGGGTTTGGTAGATAACGAAGATA
GAGCAATCCGAAAGGCTAAAGATGACTTTAGGGCTAGCACGCTCGTCTCGAACGGGTTAGAATTTTTTACGGAAGCTTGGTATGCT
GAACAAATGTGGGCTAGCTTAAGAAGCCATCCACATTTCAAGCAGATAGTTACAAGTCGTTTGCAGCAAGGCGACTTGCAAATGCT
CGGAAAAGTTCTGTCTGAATTGAGCATCGGAAGGCAACCGTCGTTGTGGGAAGATTTAAAGCATTGTGAAGTGCCGCTTCAGATAA
TCGTGGGCGAAAAGGATGACAAGTTCAAGAAAATTGCTCGTGAGATGTACGAAAAAATCGGCTTTGAAAATGGAAATAAAAGCTCG
CCTCCGGTTGCTGAAATTCCAAATGCTGGACATGCTGTCCATCTTGAGAATCCCCTTGCTCTTATCACTGTTTTAAGGCAGTTTAT
AAAAAGAGAAAATAGTACTTAATAAAAAAAATTAAAACAAAAAAAAAATTATGTTCAGTTTTTCAAAATAAAATAATACAGATTAC
AGAAATATATTCATATTTCTTGTAATAAAGATTGAACAAGTCCTTTGCTTCAAAAACAGAGAATAAAAAGATATTGAATAAGATGT
GTTTTCATTTTGTTTGGCATGACTTAT

>PaMenE Phelipanche aegyptiaca Pa.c23477_g1_i1.14490
TCTCTCTCTCTCTCTCTCTCTCTTTTGCTATATACTTCTGTAAGTTTCCCCTTCTGCCCTCCGCCGTAATGGCTAATTACTCGGAG
TCCCACATCTGCCAGTGCTTGAGCCGCCTCGCCGCCGTCAGCCGCATCTCCACCGTCACCATATATGGAGACCGCCGGAAAACCGG
AATGCAATTTGTCGAAGAAGTTATGGGCCTGGCACATGGACTTCTCCAACTCGGTATCAAGCCCGGTGACGTTGTCTCCATTTCTG
CTCTCAACAGTGATTTGTATTTGGAATGGATGCTTGCTATTACTTATGTTGGAGGAATTGCTGCTCCACTAAATTATCGATGGAGC
TTGGAAGAGGCTAAGTCAGCATTGGAGGTAGCAAGACCCGTATTATTAGTAACCGACTCAAGCCCGGGCTATTGGCATTCCAAATT
TCAGATTGATTATGTCCCGTCTCTGAGGTGGCATGTCTTGATGGATATGCCTGTCAAAGCCGACAGTACCAGGACAATTTTTGCGG
CCGAATTTCTTAAGGAGCCTGCCGGAAGATCTGTAAAAGTGGACTATCTTTGGGCACCTGAAAGAGCTGCAATTATATGCTTCACC
TCAGGAACCACTGGAAGGCCTAAGGGAGCTACTATAAGTCACTCGGCTTTAATTGTGCAATCCCTTGCAAAAATTGCAATTGTTCG
CTATAATGAGGATGACGTATATCTTCACACTGCTCCCCTGTGCCATATAGGCGGAATATCATCAGCCTTGGCCATGCTAATGGCAG
GAGGTTGTCATGTTATATTACCAAAGTTTGAGGCTAGCTTAGCAATTGAAGCCATTAGGGAACACAGCGTCACTTCCTTGATCACT
GTACCCACCATGATGGCTGATCTAATCTCTTCTCATAGAATGGATCAAACATCTACAAGTTTCGAATCTGTGAAGAAGATCCTGAA
TGGAGGTGGTGGTCTATCAGTTGAGCTCATAAAAGATGCAACCAAATTATTTCCACTAGCCACGCTTCTCTCAGCTTATGGGATGA
CAGAGGCATGTTCTTCTCTTACCTTCATGACTCTTTACGAGCCGACAAAAGAAGGCCATCACTTGCAGCCACATGATATACAAAAG
TCCAACTTAATTAGTTGTCAAGGAGGTATATGTGTAGGAAAACCTGCTCCACATGTTGAACTACAAGTAAATGCCGAGGAATCTTG
TAATACCGGGAGAATATTGATGAGGGGTCCCCACGTAATGCTTCGTTACTGGGGTCAAAGTCCATCAAAACATTTGAGTCCTGTTT
ATGAAGGTTGGCTTGATACTGGGGATATAGGGCAGGTAGATAATCATGGTAATTTGTGGCTTATTGGACGAGCAAAGGATAGAATC
AAGAGTGGAGGGGAAAACATTTATCCAGAAGAGGTAGAGGCTGTCTTATCCCAACATCTTGGAATTTCTAGGATTGTTGTTGTTGG
AATTCCAGATTCTCGGTTGACGGAGATGGTATTTGCCTGTATTAAGCTAAACGACGGTTGGAGATGGACTGATTTTGTTGGCAATC
ACTCAGCAGGAGAACATATACAGTGTTTGTCCAGTGAGATACTCAAACGCTTTTGTAGAGAAAAAAATTTAACAGGGTTTAAAATT
CCCAAAAGATTTGTTTTGTGGAAGAACGATTTTCCAATAACAACAACTGGAAAATTAAGGAGAGACCAAGTCAGAGCAGAAGTTAT
GTCTCATACTCAGTTCCTCACCAGCAAACTTTAATTGCCACAAAATATTTTCTGGGAAATAATGATTGTACAGAGTTTGTGCACAT
TTTCTTTAAATTGATCAAATCTGGACAACAAAAGGGTGCGTGTTCATTTTGAATTTTGATAAGAATAAAAGTTGCTATGTATTTAA
AATGGGGTTGGTTGAGTTATTGAATAATCTAATAATTAATACCTCATCCGTTTCAGGAATATTGCAAAAAAAAAAAAAAA
>ShMenE Striga hermonthica Sh.c16227_g1_i2.0
CGGGGTCGCAGGCACTGAAAAAAAAACCAGTGTACAAACGCCTAGGCCCTGCGTATTAGGGTGCTCGCTTAGCGCCTAGCGCCTAAT
TTTCCATTGGCCACGTCGCCTCACTTCGAAACGACGTCGTATATCAGGCAAAGCGTTTTCGTCTCGGATAGAGCAAACGGTGTTGA
TATTACTCATTCGCAATTCTACAAAGACGATGTCATCTCCCTTTTTGTGTCGGTCGATTCCTCTTGAAATTTCAATCCTGTGTGCA
TATTTCCTTTGCAAAAGATCTGTTTATTTTTTCTTTTCTTTCCCATTTGGGTCTGATTTTTCCATGGTCGTCTAGGGCGAGCTCTT
TGGTGTGAACGGCGAGCGCAATTTCCTTCAGCTCTCGAGCAGGCCAGAAAGATTGAAGCACGACGACCTCGAGGCCATGGATTGGA
GGGGAACCGATTTCAATTTCATCATATCCGATTCTGGGCTCTGATGGTTCAGATCCATTTCTGCGGAATTTCGCTCATGGTATCGA
AGGAGCTGGCGGCCGCGTGAGGTTTTTGCGAGAGGAAAAATCTCATAATCGTTGGAATCACGCTTCTTTTTAAACGGAATCTTAAG
CCTCTTCGCCAGAATCTTGATCCTATATAATTGGTTCGTTGAATCAACGAGTTTTGGAGCATTTGTCGACCTTGCTGAAGTATTCT
GATGCTTAAAACCTGTTCGGCCCCTGATCACTTTGTACTGGCTATCAAAGTTGTGTGTGCCCGTAGTAGGGGAATCGAAGATATCA
ACCATCGACGGCCTCTGCTCAGCTCAAAACGCACAATTCAGCCGTTTCTGGCCCAGAGAATAAGCCTTTACCGTTGCCGGCGAATT
CGGGATTTTGGTTGGCATAACACATGTACTTACAGCAGAGCTGGAACCTCCTGGTGATCAGCGGCGGATTGAATGATGCTGCTCGA
TTGCATTTGATTTTCGTTGGACAAAAAAGGGAATTGCTAAAGTTGAAGAAACTTATAGTTCTCACAAAAGAAGGAAAATGTATGAC
CTTGACTATGTTTTCTCCTCAGAGTGACTTGTATTTGGAATGGATGCTTGCTATCACTTATGTTGGAGGAATTGCAGCTCCACTGA
ACTATTGATGGAGTTTGGAAGAGGCTCGTTCGGCGATGGAGTTAGCAAGAACCGTTTTGTTAGTAACCGATTCAAGCCTGGTTATT
GGCATTTTATGTTCCGTTTTTACGATGGCACGTTTTGATGGATACGAAAATTGAAGCTGAGAGTAACAGGACAATTTTCGCATCTG
AATTAGTTAAGGAGATTGGTGGAGGATTTGCTGATGTAGACTATAGATGGGCACCTGAAAGAGCCGCGATTATATGCTTCACGTCA
GGGACAACTGGGAAGCTGAAAGGAGCTACAATAAGTCACTCGGCTCTCGTTGTGCAATCCCTTGCGGAAGTTGCAATTGTTCGTTA

TGACGAAAATGATGTATATCTTCACACAGCTCCACTGTGCCACATTGGCGGAATATCATCAGCAATGGCCGTTTAGATGGCCGGGG
GTTCTCATGTTATATTACCAAAGTTCGAGGCTAATTTAGCAATTGAAGTCATTAAGAAACACAATGTGACGCCCTTGATCACTGTA
CCGGCACTGATGGCCGATTTAATCTCTTTTAATCGATTAAACAGAACATCAGTGACTTACGAGTCGGTTAAAAAGATTCTGAATGG
AGGTGGTGGCCTGTCGACCGAGCTCACTAAAGACACAATCGAAATTTTTCTGAGTGCCACACTTATGTCAGCTTATGGGATGACAG
AGGCATGTTCGTCTCTTACCTTCATGACTCTTTTCGACCCAACCAAAACGACCCATCAGAAATACCACTCGAGTTTCCAAGGAGGC
ATTTGTGTAGGACAATCGGCTACACATGTCAAACTAAAAATAAGCCCGGACGAAAAATCTTTAAACACCGGGAAAATACTAACGAG
GGGTCCACACGTAATGCTCGGTTACTGGGGTCAAAGGACATCCAAATCCAATCATTCGAAACCAGGCCTCGAAGGCTGGCTCGATA
CTGGAGACATAGGGCAGATAGACGATCACGGTAATCTATGGCTTGCCGGACGAGCAAAGGATAGAATCAAGAGTGGCGGGGAAAAT
GTTCATCCTGAAGAGGTTGAGTGTGTCCTATCCCAACATCCTGGGATTTCGAGGATTGTTGTTATTGGGATTCCAGATGCTCGGTT
GACAGAGATTGTAACCGCATGTGTTCAACTAAAAAGCGACTGGCAATGGGTTGATTTTGGCGCCGGTCGGTCAGCAGGAGTCGAAC
AAGTCAAGTGTTTGTCGAGTGAGATCCTTAGGCAATTTTGCAGACAAAAGAACTTAACAAGATTTAAAATTCCAAAAAGGTTTATT
CTGTGGGAGAACGATTTTCCAACGACGACGACTGGGAAGTTAAGGAGAGACCAATTAATGGTAGAAGTTATGTCTCATACTCAGTT
TGTGTCCAGCAAACTATAGACAGTTCACACATAATCACAAATATGTGAAAATTAGTGTATAGAGTTTGTAGCTTAATGTGTGGCCA
TTTCTTGTAAGATAATAATAAGTACAGGGAGATTAATGTGAAATTTGGTGATCAAATTGAAGGATGTGAATGTTGAGAACTGATGT
CCTGAAGCATTTGTGTAATAAAATTAATCCTTTGGGGTTTGTGTAATAAAATTCTGCATTTTTTTTTTTAATATATAGAAGGTTGA
GTATGTTTTGATTCTTACACAAATTTTGTGTGGGTCAGCCTAACCAGCAGACAAATAAATTG
>TvMenE Triphysaria versicolor Tv.c6492_g1_i1.12968
ATTTGAAACTCAGTTAGACCCCAAAGCCTACTCAAATGATTTTGCCAATTTTAAAACGATTTTATTATTTTAATGGGAAAAGAAAA
CACTATAAAATGGTTCAATAGCATATCATCAACTCTCTGCCTCTCCCTCCATATCTTCCACTATCCCTCTCTAAACTCCTAATATC
TTCGTACATCTCTGAATTTCCCAATTTGCCCTCCGCCGTAATGGCAAATTACTCGGAGCCCCACATCTGCCAGTGCTTGAGCCGCC
TCGCCGCAGTTAGCCGCGATCCAACCGTCACCATTTGCGGCGATCGCCGGAAAACAGGAACGCGATTCGTCGGAGGAGTTATGGGC
CTGGCCCATGGGCTTCTCCAACTCGGAATCAAGCCCGGTGACGTTGTCTCCATTTCTGCTCTCAACAGTGATTTGTATTTGGAATG
GATGCTTGCTATTACTTATGTTGGAGGAATTGCTGCCCCTCTCAACTACAGATGGAGCTTGGAAGAGGCTAAATCGGCAATGGAGT
TAGTCCGGCCCGTATTATTAGTAACCGACTCGAGCCGTGGTTACTGGCATTCGAAGTTTCAAATCGACTCTGTTCCGTTTCTAAGG
TGGCATGCCTTGATGGACATGCCTCTTAAATCCCATAGTAGCGGGACAACTGAATTGCTGAAGGAGCCTGCTGTACTAGACTATCT
TTGGGCACCTGAAAGAGCTGCAATTATATGCTTCACCTCAGGGACCACCGGAAGGCCAAAGGGAGCTACCATAAGCCACTCAGCCT
TAATCGTTCAGTCCCTAGCAAAACTTGCAATCGTTCGCTATGACGAGGATGATGTGTATCTTCACACAGCTCCATTGTGCCACATA
GGCGGGATATCGTCAGCGCTAGCCATGCTAATGGCAGGGGGTTGTCATGTTATATTACCAAAGTTTGAGGCCAATTTAGCAATTAA
AGCCATTAGGGAACACAATGTGACGTCTTTGATTACTGTACCCACGATGATGGCTGATCTAATCTCAATTAATAGGATGAATCAAA
CATTTGAGACTTTCGAGTCTGTAAAGAAGGTTCTGAATGGAGGTGGTGGTCTTTCAGTTGATCTCATAAAAAATGCAACCGAAATT
TTCCCAAGAGCCACACTTATCTCAGCTTATGGAATGACAGAGGCATGTTCCTCGCTAACCTTCATGACTCTTTACGATCCAACAAA
GGAGAATCCACATAAACAGTCGTCGTCATATTACAATGATGTCCGAAAATCCAACTTAAGTCGTGAAGGAGGTACTTGTGTAGGAA
AACCTGCTCCTCATGTTGAGCTAAAAGTATCCGGCGATGGGTCTCCTAATAATACCGGGAGAATATTAATGAGGGGTCCACATGTA
ATGCTCCATTACTGGGGTCAAAGTCCATCGGATCATTTGGATCCTGTTTATGGAAGTTGGCTCGATACCGGAGATATAGGACAGTT
AGATGGTCACGGTAATTTGTGGCTTGTCGGACGAGCAAAAGATAGAATCAAGAGCGGAGGGGAGAACATTTATCCAGAAGAGGTAG
AGGCTGTCTTGGCCCAACATTCTGGAATTTCCAAAATTGTGGTTGTTGGAATACCGGATTCTCGGTTGACGGAGATGTTAATAGCC
TGTGTTAGACTAAAAGACGGTTGGCGATGGGCTGAGTTTGGTGCCAATCGTTCGGCAGGAGACCAAATCCAGTGTTTGTCCAGTGA
GATGCTTAGACACTTTTGTAGAGAGAAAAACTTAACAGGGTTTAAGATTCCAAGGAGATTTGCTTTGTGGACGAATGATTTTCCGA
TGACAACAACTGGGAAATTAAAGAGAGATCGAGTCAGAGAAGAAGTTGTTATGTCTCATACTCGCTTTCTACCCAGCAAACTATAA
TTCCCTCGATATATTTTATCAGGGATGAAACACATTCAAATGTAATGATGTGATTTGTATTTTCAGTATCCCCTACTGTGATTAGC
AAGAATATAAGCCTGAGTGGCTTTGGAGACTTGAAGAGGCAAAAATGTAATTAGGGCTATACTTAAAAGAATTTGTGTTCACTATT
CATATCATTTTTAGTATTAGATTGTGTTGTAAAAAAAAAAAAAAAAAAAAAAACCAAAAAAAAAAAAAAAAAAAAAAAC

>PaMenB Phelipanche aegyptiaca Pa.c21895_g1_i1.5023
CCGAAACCCCGTGACCCAAGAAAACTACTTCCCCAAACTACAAAACAACATTATAAATACTTCTTCCCATTATTTCCATCTCCACT
AATTTCAATACTCCCCTTAAAGTCAAAACATAATTGCCTTTTACAACTGATTATCACTCTAAAACAATGGCGGTCACTATGACAGG
AAAAGATGCTGAGATAATCAACAGAAGAATGGCCTCAGTTGCCAGACATCTCATCCCAGCTCAAAACCCAGACACAAATAATAATA
CTTACAATTTCATTTCCGGATCGAATTGTAGCTCCAAATTTAACGACACGTATCACCGGGTCCAGGGAGAAGTCCCGACGCATAAC
CCGGAATGGAAACCCGCCCTGGATGAGTCCGGCAAGGAGTTTACGGACATTATCTATGAAAAAGCTGTCGACGAGGGCATCGCTAA
GATAATGATTAATAGGCCGGAGAGAAGAAATGCCTTCCGGCCACAAACGGTGAAGGAGCTGATGCGGGCATTTAACGACGCCAGGG
ATGATAACTCGATCGGAGTTATTATTTTCACTGGGAAGGGCACCAAGGCTTTCTGCAGTGGAGGGGATCAATCTCTAAGAGGCAAG
GAAGGTTACGTCGATTACGATAATTTTGGTCGCCTTAACGTTTTAGATCTTCAGGTGCAAATTCGCCGCCTTCCGAAGCCGGTGAT
AGCAATGGTTGCGGGGTATGCAGTAGGGGGTGGACATGTGCTCCACATGGTGTGCGATTTAACAATTGCAGCTGATAATGCTATTT
TTGGTCAAACAGGACCTAAGGTTGGAAGCTTCGATGCTGGTTATGGGGCTTCAATAATGTCTCGTTTGGTTGGGCCCAAACGGGCA
CGCGAAATGTGGTACACGGCAAGGTTTTATAATGCAACGGAAGCGGAGAAAATGGGACTTGTCAACACTGTTGTTCCTCTGGAAAA
GCTTGAAGAGGAAACAATCAAATGGTGCAGAGAAATTCTGAGGAACAGTCCGATGGCTATTCGTCTGTGTAAATCAGCTATCAACG
CAGCCGATGATGGTCATGCCGGACTTCAGCAAATCGGAGGAGATGCAACGCTTCTATTTTACGGGTCAGAGGAAGGCACGGAAGGG
AAGAATGCATACTTGCAACGCAGAAAACCAGACTTCTCAAGATTTCCTAAGCTTCCATGATATTAATATATATGTGGAGTAGCAAA
GTATTTTTCATATGGTTATGCATGTTTGAGATGAGAAATAATGGGTTGGGTGAGAAAGAATAAAGTTGTGTGAAGATTTGATTTGA
CTACCTTTTTTTCTTTTACGTATCTTTTCTGTTGTATCACAAAATATTGGTTATCAATAATCTTTTACTTCGTTTACTTGATGTTC
AATTCCACGAGTTTCGTATAGTTTTATGTTCTGAACTTAATTAGCAATGGAATAAGGTTTTTCAATATTTCAAATCAAAGATATTG
TACATCCTTTTATATCACAAAAAAAAAAAAA
>ShMenB Striga hermonthica Sh.c17026_g1_i1.3451

GAGTAATTTAATCAAACCATCAACTTCACAATTACAGCTCTTGTTAAACTAAGTCCTATAAATACAACCTTGAGTTATCCTCATTT
GTGCACCAAATTTCAGTTCTAGTTAATTTCGACAGTTCGATTAAATTACTCCGACGAGATGAACGGGAAAGACGCCGAGATAGTCA
ACCGGAGAATGGCGTCCGTCGCCAGACATTTAATTGGGCCTCAAGGCCCGAGCCCAAATAACGCCCTCATATCCGGGTCGGGGTGC
AGCTCCGGATTCAACGACACGTACCATCGGGTCCACGGGGACGTCCCGACCCACGACCCGACTTGGAAACCCGCTTTGGATGAGTC
GGGCAAGGAATTTACCGACATTATTTACGAGAAAGCCGTCGGGGAAGGCATCGCTAAGATAACGATAAATAGGCCAGAAAGGAGAA
ATGCTTTTCGGCCACAAACGGTGAAAGAGCTAATGCGCGCGTTCAACGACGCAAGAGATGATAACTCTATTGGGGTTATTATTTTC
ACTGGGAAGGGCACGCAAGCCTTTTGCAGTGGAGGCGATCAATCTCTTAGAGGCAAAGAAGGCTACGTAGATTATGATAATTTTGG
ACGTCTCAATGTTTTAGATCTCCAGGTGCAAATTCGACGCCTACCTAAGCCAGTGATAGCTATGGTTGCTGGGTATGCAGTTGGGG
GTGGACATGTGCTTCACATGATTTGTGACTTAACAATTGCAGCTGATAATGCAGTGTTTGGCCAAACAGGGCCTAAGGTTGGAAGC
TTTGATGCTGGTTATGGAGCTTCGATAATGTCCCGTTTGGTTGGGCCGAAGCGCGCACGCGAAATGTGGTACATGACGAGGTTTTA
CAATGCTGCCGAAGCAGAGAAAATGGGACTCGTCAACACCGTTGTTCCTCTCGAGAAGCTTGAAGAGGAAACAATCAAATGGTGCA
GAGAGATTCTGAGGAACAGCCCAATGGCTATTCGTCTTTGCAAATCGGCCATTAATGCGGTCGATGATGGCCATGCCGGACTTCAG
CAAATCGGGGGAGACGCTACGCTTCTTTTTTACGGGACAGAGGAAGGTACGGAGGGAAAGAATGCGTACTTGCAGCGTAGAAAACC
CGATTTCTCCAGATTTCCAAAGCTCCCATGATTTCATATTCATATGTGGAGTTCTTAATAATAATTGATGGAAATAAAAATAAAAA
AAATAAAAATAAAATGAATAAAGTTGTTTGTTGAGTGATGATTTTAAGTAACTTATACTAATTTCATCATTTTGTATCAGAAAGAA
ACAGATGCATGCTGCTTATTATCAATAAACTTGTAATTTTCTTATGTTTATTTCAGTGCTTCTGAGAATTTTGTGACATTCATAAA
TTTTTATGTTGGCAGTAGATTAATTATACGGGAAATGCTACAATCCTAGATCATCATATAACCATCGTGGATAGTTGATCTAACGG
TT
>TvMenB Triphysaria versicolor Tv.c8078_g2_i2.0
CTCCGGCCGACGGAGATATCCCTTCACACTCATCCATCACCTTCTCTTCACACTTCACACTCTAAACCCATTATAAATACAACGTC
CCATTAACCCAATTTACCCAATAACCCCACTCAAATTCAAGAAATTCATTTGCCTTAATCACCTGATTACGTCATCCAAAACCAAT
GGCTAAATTAACATTCGAAGATGCTGACATAATCAACCGGCGAATTGCCTCTATTTCCCGGCATCTCAGTCCACCTCAAAACCCGG
ACCCGAATAACCCAAATCTCATTTCCGGGTCAAACTGCAGCTCCAAATTTAACGACACTTTCCACCGGGTCAACGGAGAAGTTCCG
ACCCATATCCCGGAATGGAAACCCGCTTTGGATGAGTCCGGCAAGGAGTATACCGACATTTTATACGAGAAATCCGTCGGAGAAGG
CATCGCTAAGATTACGATTAATCGGCCGGAGAGGAGAAATGCTTTCCGGCCACAAACGGTGAAAGAGCTAATGCGTGCGTTTAATG
ATGCTAGAGATGATAATTCTATCGGAGTTATTATTTTTACCGGGAAGGGCACATTGGCCTTTTGCAGTGGAGGTGACCAATCACTA
AGAGGCAAAGAAGGTTATGTTGATTTTGATAATTTTGGTCGCCTTAATGTTTTAGATCTTCAGGTACAAATCCGTCGCCTTCCCAA
GCCCGTGATAGCAATGGTTGCAGGGTACGCAGTGGGGGGTGGCCATGTGCTTCACATGGTTTGTGATTTAACAATTGCAGCTGATA
ACGCAGTTTTTGGCCAAACAGGACCTAAGGTTGGAAGCTTCGATGCCGGTTATGGGGCTTCTATAATGACTCGTTTGGTTGGGCCG
AAACGAGCACGCGAAATGTGGTACACGACGAGGTTTTATAATGCAGCCGAAGCCGAGAAAATGGGACTAGTCAACACTGTTGTTCC
GCTTGAGAAGCTCGAAGAGGAAACAATCAAAATGTGTAGAGAGATTCTGAGGAACAGTCCGATGGCTATTCGTTTATGCAAATCAG
CGATTAATGCGGTCGATGATGGTCATGCGGGCCTTCAGCAAATCGGAGGAGATGCAACACTTCTATTTTACGGGACAGAGGAAGGG
ACGGAGGGGAAGAATGCGTACTTGGAACGTCGAAAACCGGACTTCTCGAGATTTCCTAAGCTTCCATAATTTAGAGTATGAGTTTA
GGAATAATGTTTGGAGAGAAAATAATAAAGTTGTGTGGTTTTTATTCTTATTTATTTTTATCATGTAGTGTCACAAACAATTGGAA
ACTATAATTTCCTTAATAAAAGATGTGTTATGATTCTTATGAATGAAATCAAGATTTAAAAGATGTTGAGCATATTGATATGCTAT
GAAACACGTTGTTA

>PaDHNAT Phelipanche aegyptiaca Pa.c23042_g1_i1.10619
TTTGCCTCAAGTTTTTATGCTATTAGCTCATAATTTTGACCCGGAGAAAAATTGGTGATCAATATTAGTTCTCTTTTCTGGTAGAG
TTGGTAAAAAAAAAATAGTAGATCACCGCAACAAATTTGATTCAAATTAGTCGAGAGAAGAGATGAACCAACCACCACCTTCCGCC
AGGCCACCGTCACCTCCGTCGAACACAAAGGAATTGGATTTTCCGCTCCACACACTCGGTTTTAAATTCGACTGCCTTTCGCCCGA
TAAGGTTTCGGGACACCTCCTTATCACTTCAGAGTGCTGTCAGCCGTTCAAGGTGTTGCATGGAGGGGTGTCGGCTCTTATAGCGG
AGTCTCTAGCAAGCATAGGAGCTCACCTTGCTTCTGGTCTGCAGAGAGTTGCCGGTGTACACCTTAGCATCAGTCACCTTAAGAGT
GCTAAGTTAGGTGACACTGTTCTTGCCGAAGCTACACCTGTCAACATCGGTAAAACTATTCAGGTCTGGGAGGTGAACTTATCGAA
ATGTGATCCTTCGGATTCCGAGATTAAAACGTTGATCTCATCTTCAAGAGTCACTCTTCTTTGTAACTTGCCTGTGCCAGAATCAC
TCAAGGCTGCTGCTCAAGGTCTCAAAAAGTACGCAAGACTGTAATAATAAGCTCAAAATTTAGTACTGTTATTATAATCTCTGAAA
CAACTCTATGTTGTGAAAAAAACGAAAGAAAGTATCGAAATATAACTAAAATGATGTAATTTACAGAATTTATGTTATACGCAGGT
CAACTTTTCATTCAGATGCTTTAAAAACTCTCAGCATTAAGTTGGGGTCTTTCACAATCTTCACATATTGAAAGAGGCATAATCTT
GGATCCAACACATTTTGTGCGATACATAGTATTTATTCATATCAATTCTTGTTTGGGGATCAACAATGTGGAATTTTTCGGACCCA
TTTACTCTCTTTGAGCATCTCGAGCAGAGCGATGGAGATGCGTTTTTTCGCAAAATGGAAGCAAATTGGGGGTCGCAGTGCGCTAT
TTATTTGCGCCAGTCTTGTGCTTGGAATAGTCCTGCGCCATTTTCAGCGCAGGACTCTAGTGCGCTATTTTTTTTTATATACTTTT
TTGTTGTTTATATTTCTTTTTTTCTTTTAATGTAAACAATTTAATATTAAATTATTATATCTAATATTATAATTATAGAAAAAAAA
AAAAAAAAA
>ShDHNAT1 Striga hermonthica Sh.c83733_g1_i1.18685
GTCCATGCAGTACCATCATCCTGAATTGTCATTATTGTCTAAATGTAAGAGAGCTACAATAACAAACATGTATCACCCACATACAA
TCAGATTCTAGACTACACCACGGTGGCTTTTCTACCCTATCTCAGTTTTTCTTCTTCTCCTGTTCTTGAAATTCGTTAAACTGATC
TACTCAGAGTAGATTGGTCTGCGAATAATCTAAAACCATAAAGACACAAGAAGCGACTCAGCAGAGAGAAGATGAACCGACCGCTG
CCATTAAACACAAAGGATTTGGATATTCCACTCCACACGATTGGGTTTGAAATTGACTGCATTTCGCCTGAAAAGGTCTCGGGCCA
TGTTCTGATCACTGAAAAGTGCTGTCAGCCGTTCAAGGTGCTGCATGGAGGAGTTTCGGCACTAATAGCCGAGGCTCTAGCAAGCA
CTGGGGCCTACCTTGCTTCAGGCCTTCAGAGAGTAGCTGGCGTCCACCTAAGCATCAATCACCTAAAGAGTGCAAAGTTAGGGGAC
TATGTTGTTGCGGAGGCTACTCCCGTGAACATCGGGAAAAGTATTCAGGTCTGGGAAGTGAACTTATCAAAATGCGATCCTTCGAA
CTCAGAGACAAAAACATTGATTTCTTCTTCAAGAGTTATTCTTCTTTGTAACTTGCCTGTGCCAGATTCATTAAAGGAGGCTGCTC
AAGGTTTCAAAAAGTACTCCAAACTCTAGATAAGTTCAAAACCGATTTTCTTCTGTAATTTTCCGAAACTATTGTGTTGTAATAAG

```
ATAAAAGGAAAGCAAAAAGGGAAGGAAAATGAACCCTTTTGCAATTACAGCAAAATGTCACCCTACCAAATATGCAGAATATATGC
TAGGCAGTTGGCACAAGTTATTGTTCGGTTGGTCTGTGATAGCAATTAGCAAGGTGGAATTTGTCTTACCAGCATATGATATATTA
TGCTTCGTATAAACTATGACAAGTCTATTTGAATTTTGTGGTTATGATATATTGGATAACAATATATTGCTCTTAGTTTTGATGTT
GTTGTTTTTGTTCCGAAAATTGCTCTTAAATAGTAATCACATTGCACTACAATTAAGAATACAATACCATGAGGATTTCATTTGAG
ATCATGAGATCATAAACTATGTTATTACACACTTCAAATGAATGACATAACATGACTAT
>ShDHNAT2 Striga hermonthica Sh.c13237_g2_i1.9518
GGGATAATAGATAAAGCTGCCATTTTTTTTTCTTGCTTCAGCTTCTCTACCACATTCCCAGATAAATTGTTTTGTGGTATAGTAGT
GTGTGAATAATATACAAAACCACAAAGGCACGAGAAAAAACTCGATTTCTATCCCAGTCGACCGAGCGGAGATGAACAGAACACCA
TCCGCCACAGGGCCTCCGCCACCACCACCATCAAAGACCAAGGAATTGGATATTCCACTCCACACGATCGGCTTTGAAATCGATTG
CCTTTCGCCTCAAAAAGTTTCGGGCCACGTTTTGATTTCTGAAAAGTGCTGCCAGCCGTTCAAGGTGCTTCATGGAGGTGTTTCGG
CACTGATAGCCGAGTCTCTAGCAAGCATTGGGGCCCACTTAGCTTCAGGCCTTCAGAGAGTGGCCGGCGTACACCTCAGCATCAGT
CACTTAAAGAGTGCAAAATTAGGCGATTTTGTTGTTGCCGAAGCTAAGCCTGTGAACATAGGGAAAAGTATTCAGGTTTGGGAAGT
GAACTTATCGAAGTCCGATTCTTCAAACTCTGAGACCAAAACGTTGATTTCATCGTCAAGAGTTACTCTTCTTTGTAACTTGCCTG
TGCCCGAGTCACTCAAGACTGCTGCTCAAGGTCTCAAAAAGTACGCGAAACTCTAAATAAGCTAAAAACCGGTTTCTTTTGTATTG
CTAAAACTATTTTTTATGTAAATAAAAGACAAAAGGGAATGACAATGAGCCCTTTGCAGCTATAGCAATTATCACATCACCCTACC
CAATTTGCATAATCTATGTTTGGCACAAGTTATTATTGCTCAGATGATTAATAAATGTCTATTTGATGACAATTATCAGTCGATTG
TTCGCTCTATCCATCGAGCTATAATGTAAATTAAATAGAAAAAAAAAATACTGTGTGAATTTTTCCAGTTGAACGCA
>TvDHNAT1 Triphysaria versicolor Tv.c6470_g1_i1.9012
CCTTATTATTTTTTATTATACATATCAATGGAAGATTAAACTGTGCACCCGATTGTTTATTTCGTTTGAATCTTACTGTGCTCTGT
CTTGTATTTTGGTTTTCTGCTGTTTCTTTGCCTGGTTTCTGTTCTCTCGAGTTGTCAAAAACGCCCATTTATTCGAGAGGATTGTG
TTTTGGGCTGTCAAGGTTTCAATCAATTAAAAACGCATAAAGTATGAGTGAATCACTACCACCAGCAGTAAAAAGGATGGAGCTTT
TGGATGCTCCACTTCACTTATTTGGCTTTGAACTCGATGAACTTTCGCCTCATAAAGTTTCCGGCCACCTCTTAATCACTTCAAAG
TGCTGTCAGCCATTCAACATGCTGCATGGAGGAGTTTCGGCTCTGATAGCTGAGGCTTTGGCAAGCATAGGAGCTCATATGGCTTC
AGGGTTTCATAGGGTAGCCGGTATACAACTGAGCATCAATCATCTAAAACCCGCTCAGGCCGGAGATTTTGTAATTGCTGAGGCCA
CACCAGTCAGTGTTGGTAAATCTGTTCAGGTCTGGGAGGTGCGTCTCTTCAAATGCGATCCTTTGAAATCCGATGAGATTAGAACG
TTGATTGCGTCATCGAGAGTTACGGTATTCTGTAACTTGCCTGTACCAGAATCTTCAAGGGATGCTGCTCAAAATCTCAAAAAGTA
TTCGAAGCTATAAATCAGTAATCTTGGTCAATGATTATCGGGTTTTTCACATCTTTCATCTGTATGTTTTGAAACCTGATGTTGCT
TGCACTTCCAGATCCTCCCTGTTCATGTATATATATTTCAGGTCCACAGGTAATGTCTATGGCTCGAACATTCTTCGATCCATCGA
CGATTGAAATGCAGTCGTCGCCTAATTGATACGATATTAATTAACAAACGTGTTACTACAAATTAATTCGAAAGTAATATATATTA
ATGAAGAAAATATCGTACCGGTGCCAATAGTACAGCTAGAAATACGGATATTTTGAGTGGCAGTGACATGTATGCCATCAGTGTTG
GGGCTTTCATCAGGAGCAGTCACAACAAGTTTTGAAGCTTCAACATTTTTGCACCTTTGAAATGACAATTGCATTTGCTGTGCATT
TTTTATCTTCAGATTTTGTACTACCAAGTCTATGCACTTGTAGAAGGTTATAGCCTGCACAATTCATTTTCTTCATTAATATATAA
TCGTAATTATATAAATTGATAAATAACACCATTTTATGCCATGAACAAGATGCCTATATACATAAAAAATTTAAAAAT
>TvDHNAT2 Triphysaria versicolor Tv.c6736_g3_i3.9225
AAATCATTTTTTTATTGGAATTCAATCCTTAGTGTCAGAGATGATAATAATAGACCCATACGAGTTTGCGGGCCATCCATTTACTT
GGATTTTTTTTTCGATTTATCGTCCAAGTGGAAAAACACAACAACAATATGATTTGTGAAGCTGCTATATTTTCTCCGCCATTGTA
GACTAGAACTCTGCTTCTTCATTGCCCAATATACAAATTAAAAGCATCTATAGTAGATTTGTTTCCTAATTGAGAGAAGAGCCGAG
AAGATGACCCAACCACCATCTGCCGTCGGGCCGCCGCCACTACCACCACCAGCGAAGACAGAGGAATTGGATTCTCCGCTTCACTT
GATCGGCTTTGAAATCGATTGCCTTTCGCCGGACAAAGTTTCTGGCCATATTATTATCACTTCAAAGTGTTGCCAACCGTTCAAAG
TGCTTCATGGAGGTGTTTCGGCACTAATAGCCGAGTCGCTAGCAAGCATAGGAGCTCACCTTGCGTCCGGTCAGCAACGAGTTGCC
GGTGTACACCTCAGCATCAATCACCTTAAGAGTGCAAAGTTAGGTGATTTTGTCGTTGCCGAGGCTACCCCTGTCAACATCGGCAA
AACTATTCAGGTGTGGGAGGTTCGTTTATCGAAATGCGACGATCCTTCAAACACTGAGATAAAAACGTTGATTTCGTCATCAAGAG
TTACTCTTATTTGTAACTTGCCTGTGCCTGAATCGCTCAAGAGTGCTGCTCAAGGTCTCAGAAAATACGCCAAACTATAATAAGCT
CTTGTTTTTTGAGCATATAATAAACCCCAAATCGAGTGTTGTTGTAATCTCTGTAACAATTTAATGTTTTTTTTTTTTTGGAAATA
TATAGCAAACAGTGCTGTTTGTGGAACCGGAACCTATGTATGTTACACGCACGTCAAATTTCATTTATATACTTGCGCAAACGGCA
TATTACGAGACTCTGCTTCAATTTCATGTGATGGTATATAAATTTATTTGAATCATCATTTCTGCAAAACAATTTGCATATGACAA
ATACATGAGGATAT
>TvDHNAT3 Triphysaria versicolor Tv.c102567_g1_i6.16017
GAGAAATGGAAATCTCATAGGGTTAAAATCTGGGAAGAAAAAAGGCGTTAAAATAAAAAAAAATCTCATATTATTTGGTTGTCAAG
TAAACTATAATCAAGAATCACACCCCGTTTCCACCCTATTACGTTTTCATGTGCGCAAGTGGTGTAGAAAGAGACACAACAAAGAA
AAAGGCGCAGGTGAAGATGAAAAAGCCCTAAAGCAGCAATTTTTATTAGGATTTTTGATTATTTTATTCCCAAATCCTGTGGATAT
GATTCCGCTTCTGCTCCTATTTCTACTTCACTGCCCAACAATCCCTTCATTTTTCTTCGGTATTGTGCACTGATTTCCAGAACTAT
AACGAATACAAAAACTTGATCCATCTCAGCCGAGAGAGGCGATAAGATGAACCAACCACCACCTAATGCCGGACCGCCACCACAAT
CAATGACGGAGAAACTGGATGTTCCGCTTCACTGGCTCGGTTTTGAGATCGATTGTCTTTCGCCTGACAAAGTTTCAGGACATTTT
ATCATCACTTCAAAGTCGTCTCAGGCGTTCAAAGTGCTACACGGAGGAGTTTCAGCGCTGATAGCCGAGTCACTAGCAAGCATAGG
AGCTCACCTTGCGTCCGGTCAGCAACGAGTGGCCGGTGTACACCTCAGCATCAGTCACCTTAAGAGTGCACAGCTAGGTGATTTTA
TCGTTGCCGAAGCTACCCCTGTCAACATCGGCAAAACTATTCAGGTGTGGGAGGTTCGTTTATCGAAATGCGATGATCCTTTAAAC
ACTGAGACGAAAACGTTAATTTCGTCTTCGAGAGTTACGCTTCTTTGTAACTTGCCTGTGCCTGAATCTTTCAAGACTACTGCTCA
AGGTCTCAAAAAGTATGCAAAACTATAATGGCTATATAAAATTATTTGAACATTATTATTATGATTAAATCTGTCTTGGATTTATT
TATGGGTAAATTGGACATTCAGTTCCTGTAGTATAGTGAAATTGCAAAGTCTGTGTTCAATGTTTTGATAATTGCACTTTCAATAC
CTAAAATTTGGGTAGTGACAGTGAAACAGTGGTTACTGAAATTGTTGTTTTTCTAAAAGTTTAGGTGTTGAGAGTGCAATTATTAA
AATATTTGGTATCGACTTTGTAATTTTAGTAT
```

>PaMenA2 Phelipanche aegyptiaca Pa.c24509_g1_i2.9208
TAAGATTCTATTTATATCACAGCCGCTCAGTTTTTCTCTCTTTTGTTTTTATATTTTTTCATTTACCAAGCTTGTATAACCAGGAC
CAGATCCCAAGCAAATTAAGATTCTATTTATATCACAGCTCATTATTTTCATACACTTGTGGTTTATTTTTTCATACTTTTACTTT
AGTTTGACCCTCGACAATGGCGGAAGCGGCAAACCATCCAAACCAGGCAAACATTGTTAAAAGATTACATAAGAAAGAGAAAAAGG
AAGGTGATATATCTCGAGCAACTTTAATATGGAGAGCTGCCAAATTACCTATGTACACCGTTGCATTAACTCCTCTAACTGTTGGG
ACAGCTGCTGCATATTGGGAGTCGGGATTTTACTCTTTGGAGCGTTATTTCACTCTCTTGGCGTCTTTTGTTCTTGTCAATCTTTG
GATCAATTTAAGCAACGACGTTTACGATTTTGATACTGGAGCTGATAAAAACAAAAAAGAATCTGTTGTCAATATATTTGGCAGTC
GCACGGCCATCCACATTCTTTCGCTGTTAGTACTTGCAATTGGTTTCGCGGGGCTTTGTTGGGTAGCTCTTGAGGCTAAAAACCCG
CGCGCTATTCTATTATTGGCTTCGACAGTATTTTGCGTCTACGCTTACCAGTTCCCACCGTATCGTTTAAGCTATCACGGACTGGG
AGAGCCCTTATGTTTTGCAGCATATGGCCCTTTTGCCACAGTTGCTTTTTATTTGCTCCAAAGCGGCTCTTCAAGTGAGCTACCCA
TATCTAGCACGGTTGTATTTGCATCAATTTTTGTTGGTTTTACATCAGCTTTGATCCTCTTCTGTAGTCATTTTCATCAGATAGAG
GATGATAAAGCTGTTGGGAAAATTTCTCCTTTGGTGAGGCTTGGCACTGAAAAAGCATCAAAAGTTGTGAAAATGAGTGTTTTAGG
GTTCTATTGGCTTGTGTTTGGTTTAGGACTGTCCCAAACACTTCCTTATGCTTGTGTAGTGCTCTGTACTATGACACTGCCCATGG
GAAACTTAGTAGTTAGCTTTGTGCAGGAGAACCACAAGGATAAATCCAAGATCTTCATGGCTAAATACTACTGTGTAAGATTACAC
ACTGTATTCGGAGCTGCTTTGGCTGCTGGACTGGTGGCATCAAGATTAATAATGCATTCTGGAGAGCAACTTCAGCAGAGTATTTT
CGATTATGCCAAGTTTCTTTATCTCTGAGTTTATGCATTTGTATTTCTCAGTCTTTGCACACTAATAAGGAAAACTGATTCTCAGC
TTTTAGGTGGACAAGATGAATCTTCTAGAAGCTCAGAGTCGTTTTTCTTTTTTTCGTAAAATTCGCAAACACTCTCTTTAGAAAGA
GTTGGAATTAGAATTTATTGCTGCCCCAAGTTAAAAGTTTCTTAGAATGATTTCCTTTATATATATTTATATATATATTTTTATAT
TTTCTGTTGCAAAAAAAAAAAAAAAAAAAAA
>ShMenA1 Striga hermonthica Sh.c15001_g1_i2.8280
TTTTTTTTTTTTTTTTTTTCTGCTTTATTCAAGCCATAGAGCTAGATTTATGGCGCCTCTTGCTGTGGCAGCAGCTGTGTATTGTT
CCACAAGCCATGGCTATGGCGTCAAGAAGCTCGACGATTATCTTACCAGAAAACTCAGTATCAGCAGGATTCATCAAGTATTACTT
CTTCCAGATGCCTGTCAAAGGTCACTATGCACAAAATTTAATTTCAAGAAGGCCAGCATGAGGCAATTATATTCTCTACGGGGGCA
TTATATTAATTCATTTAAACAAAGAGCAGAACACTGTGGGAGTAGTAACATTGAAGAAAACAAGGAAGAAAGCATCTCCAGGGTAG
CTTTGATGTGGAGAGCTATCAAATTACCCATTTACTCTGTTGCATTGATCCCTATAACAGTTGGAAGTGCAGCAGCATATTTGCAG
ACGGGCCAATTTTTTGGAAAGCGTTATTTAAAGCTATTGTTATCATCAGTTCTCATCATAACGTGGCTCAACTTAAGCAATGATGT
ATATGATTTCGAAACTGGGGCTGATAAAAACAAGAAAGAATCAGTTGTTAATCTAATTGGCAGCCAAACAGGAACTCATATTTTAG
CTTGGGTACTACTTGAACTTGGTTTCGGGGGCCTTACTTGGGTGTCTATTGAGGCTGGAAGTATACGTTCTATAATTCTACTTGCC
TGTGCCATATTTTGTGGCTATATTTACCAGTGTCCACCTTTTCGGCTGAGTTACTTGGGACTTGGGGAACCTTTGTGCTTTGCGGC
ATTCGGTCCATTTGCTACCACTGCCTTTTACTTGCTCCAAAGCGGGACAAGTGAGCTGTCAATCTCTGCAACTGTAATCTCTTCAT
CAATTCTTGTCGGCTTCACAACGTCCTTAATCCTATTCTGTAGCCATTTTCATCAGATAGAGGATGACAAGGCAGTCGGGAAATTT
TCGCCTGTGGTCAGGCTTGGAACCGAAGGAGGTGCAAATGTAGTGAAAGTGGCCGTACGGATACTTTATTCGCTTCTATTTGTTCT
CGGGCTTGCCCAAATTCTTCCTTTCCCCTCTATAGTTCTCTGTGCTTTAACATTACCAGTTGGAAATTTAGTGGTTAGCTTTGTCG
GGACAAACCACAAGGACAAGATGAAGATATTCATGGCGAAATATTACTGCGTGCGGTTGCACACGATATTCGGGGCTGCATTAGCT
GCTGGGATGGTAGCGGCCAGAATGTTTGCGAGAAAGCAACTCCCGCATGCTATTATTCTTTGAACTATTGTTGGTGCTCATTTGTA
TAAATTTTCGATGTTCTTAAAACTGGACTGACCATTGAAATAAAGCTACAATGGACTGACCATTTAATCAAGAAGATTATATAAGA
ATTGATTTTTCAATAACTCAACCGATTCTATTTTTTTGTTTTTTTTGAATTGTAAACGAAGTTTTCCACAATATCAAAATTAACTA
TTTATCTTTAATACAAATCAATCTTTATCCTTAATAAGTAAAAAAAAAAAAAAA
>TvMenA1 Triphysaria versicolor Tv.c6621_g2_i2.1348
CTCCAGCATTGTGAGGTTGAAATTTGTCCATGTTTTACTTATCTTCTTCGTGACCCAGAATTAGTTACTCTCTGTTCCAGCTAACA
TGGTCTTTTCACATTTCTTAATCCTCACCATTTTTCTTCTTCGATTAGTATATCGTTGATCTACTGTTCACTTTCAAGCTGTAAGG
TTTGGCGCCTGTGGCTATGGCAGCGGCCACGTTCTGTTCTATTAGTATTAGCCATGGCTATGCCGTCCAGAGACTCAACAGACACA
AAATCAACAGGACTTACCAAGTATTACCTCTTGTATGTGGCAGTCGAACCACGAAAGTTCATTTGAACAAGACAATCATAAGACAA
TATTTACATTCTATACAAAGGCGTTACAAAATCTCGCCCAAGTTTAGATCAGAGAACAATGCAGACAACACTCACGTCGAAGAAGA
AGAAGAAGATGAAGATGAGCAAGAAAGTGTATCTAAGGCAACTTTAATTTGGAGAGCCATCAAATTACCAATGTACACTGTTGCAT
TGATTCCTATAACGGTTGGAAGTGCAGCAGCTTATCTACAGACAGGACAATACTTTGGAAAGCGTTATATTATGCTATTGGTTTCT
TCGGTTCTCATCATAGCTTGGCTCAATTTAAGCAATGACGTTTACGATTTCGATGCTGGAGCAGATAAAAACAAGAAAGAATCAGT
TGTTAATCTATTCGGAAGCCGGACAGGAACACATGTTTTTGCATGGCTGTTACTCGCACTTGGTTTCGCGGGCCTTGCTCGGGTTT
CTGTTGAAGCTGGGAGTTTACGTTCTATATTTCTACTTGGGTGTGCCGTATTTTGCGGCTACATTTATCAGTGTCCGCCGTTTCGA
TTAAGTTATATGGGACTTGGAGAGCCCTTATGCTTCGTTGCATTTGGCCCGTTTGCGACCACAGCCTTTTACTTGCTTCAAAGCGG
GACAAGGGAGCTGTCGATTTCTGGCATCGTTATCGTTTCGTCGGTTCTTGTTGGTATCACAACATCCTTAATTCTATTTTGTAGCC
ATTTTCATCAGATTGAGGATGATAAGGCTGTCGGAAAATATTCGCCTTTGGTTAGGCTTGGATCCGAAGGAGGTGCTAACGTTGTG
AAAATGGTTGTTAGGACGATTTATTCGCTTTTATTTATTTTGGGACTGGTCCAAACCCTTCCATTCGCGTCCATTGTTCTTTGTGC
TTTAACATTGCCGGTTGGAAATTTAGTTGTTAGCTTTGTCGAGAAGAACCACAAGGACAAGACGAAGATCTTCATGGCAAAATATT
ACTGTGTGAGATTGCATACAGTATTCGGAGCAGCATTGGCTACTGGGCTGGTTGCAGCTAGAGTGCTTGCAAGAAAGCCAATTCCT
AATGCTATTATTCTTTGAGCTTTTCAGTCCTACAAGGTGGTGATGAGGATTGTTAGTTATTTCGATTACACTTTTCAAAGAAAAAT
AAATAAAATAATCATTCATGTAATTAAATTTGTGTCATTATCACTTGTTAATAACTTGTTGATAAAATACAATCAACAATAAATGT
TTAACAATAATTCCAAATCAATCAAAATCTGAATTTAAACAAAAAAAAAAAAAAAAAAAAACAAATCTACACCGCAAAAATCTACTA
CTCGTTATCGGGCTTAGGGGCTGCTTCCTTGATCTCCTCAGAGTTATCATCCTGCATGTCGGAAGTCCACAAAGTGAGGTTGTCAC
GGAGAAGCTGCATGATCAGAGTGCTATCCTTGTATGATTCCTCCCCAAGTGTGTCTAGCTCAGCAATTGCCTCGTCAAAAGCCTGT
TTGGCGAGATTACAAGCACGATCAGGAGAATTCAAAATCTCGTAGTAAAAGACTGAGAAGTTGAGTGCAAGTCCAAGATCGGAAGA
GCACACGTCTGA
>CaMenA2 Conopholis_americana FAMO-2092133 (1KP)

TTTTATTTTTCCTTTGATAATGGCAGGAGGAGCCAACTTCATCAGTTTGGCAGACAATCCAAGCCTGGAAAAAAATGTTAAAAGAT
CAAAGAAGAAGGGAGAGAAAAAGGAAGAAGAAGATATATCTAGAGCAACTTTAATATGGAGAGCTGCAAAATTACCTATGTATACT
GTTGCACTAATTCCTCTAACTGTTGGAACAGCCACTGCATATTGGGAGTCAGGGTATTACTCTTTGGAGCGTTATTTCATTCTCTT
GGCATCTTTTGTTCTTGTCAATGTTTGGATCAACTTAAGCAACGATGTTTATGATTTTGATACAGGAGCTGATAAAAACAAAAGAG
AGTCTGTTGTGAATATGGTTGGCAGCCGCACGGCCATCTATATTCTTTCATGGTTACTACTTGCACTTGGTTTCGCGGGACTTTCT
TGGGTAGGAATTGAGGCTAAAAACCCGCGGGCTATACTATTATTGGCTTCATCAATCTTTTGCGTCTATATTTACCAGTGTCCACC
TTATCGTTTAAGCTACCATGGACTGGGAGAGCCCATATGCTTCGCCGCATACGGCCCTTTTTCCACCATTGCTTTTTATTTGCTTC
AAAGCACTTCAAAATTTGAGCTACCGATATCCAGCACGATTGTATCGGCAGCAATTCTTGTTGGTTTTACAACAGCCTTGATCCTC
TTTTGTAGTCATTTTCATCAGATAGAGGATGACAAAGCTGTTGGGAAAATTTCTCCTTTGGTGAGGCTTGGTACTGAAAAAGCATC
AAAAGTAGTGAAAATGGCTGTTTTGGGGCTCTATTGGCTTGTTTTTGGTTTTGGACTGACCCAAACACTTCCTTATGCTTGTTTAG
TGCTTTGTACTATGACACTACCCATTGGAAATTTAATAGTTAGCTTTGTTCAGGAGAACCACAAGGATAAATCCAAAATCTTCATG
GCTAAATACTATTGTGTGAGATTACACACCGTATTCGGAAGTGCTTTGGCTGCTGGACTGGTGGCATCAAGAATGCATTCTGGAGA
ATTAACTTCAGAGTTTATTAACGTACTTCGTGCCACGAAGTTTGCGTACTTCTGAGTTTATGCAATTGTTCATGATTTAGTGATAT
TGACGTACTTTGATTGTGCCAAGTTTGCGTACTTCTGAATT
>OfMenA2 Orobanche (syn. Aphyllon) fasciculata VYDM-2034145 (1KP)
CAATACACTTCGTGATTTATTCTTTACTTTTACTTTAATTGGCAATCGACAATGGCGGGAGGCAGCTCTATTAGTTTGGCAGACAA
CCCAACCCAGGAAAAAGTTGTTAGAAGATCAAAGTCAAAAAAGGAAGAAGATATATCTCGAGCAATTTTAATATGGAGAGCTGCCA
AATTACCAATGTATACAGTTGCATTAATCCCACTAACTGTTGGAACAGCCGCTGCATATTTGGAGTCAGGCTTTTACTCTTTGGAG
CGTTATTTCATTCTATTGGCGTCCTTTGTTCTCGTCAATGTTTGGGTCAATTTAAGCAACGATGTTTATGATTTTGATACAGGAGC
TGATAAAAACAAAAAAGAGTCTGTTGTCAATATAGTTGGCAGCCGTACGGCCATCCATATTCTATCATGGTTACTACTTGCACTTG
GTTTCGCGGGTCTTTGTTGGGTAGGACTCGAGGCTAATAACCCGCGCGCTATACTATTATTGGCTTCGGCAATCTTTTGCGGCTAC
ATTTACCAGTGTCCACCGTTTCGTTTAAGTTACCACGGACTGGGAGAGCCCTTATGCTTTGCATCATTTGGCCCTTTTTCCACAAT
TGCTTTTTATTTGCTGCAAAGCAGTTCAAGCGAGCTACCGTTATCGAGCACAATTGTGTCTGCAGCGATTCTTGTTGGCTCTACGT
CAGCCTTGATCCTCTTCTGTAGTCATTTTCATCAGATAGAGGATGATAAAGCTGTTGGCAAAATCTCTCCTTTGGTGAGGCTTGGT
ACTGAAAAAGCATCAGAAATTGTGAAAATGGCTGTTCTTGGGCTCTATTGGCTTGTGTTTGGTTTAGGACTGTCCCAAACACTTCC
TTATGCTTGTATAGTGCTCTGTACTATGACAATACCCATGGGAAACTTAGTAGTTAGCTTTGTTCAGAAAAACCATAAGGATAAAT
CGAAGATCTTCATGGCTAAATACTACTGTGTGAGATTACACACTGTATTCGGAGCTGCTTTAGCTGTTGGACTGGTGGCATCAAGA
TTAATGCATTCTGGGGAGCAATTTCAGAGTCTATTAACGTATTTCGACCGAGCCGAAATTGCGTAATTCTGAGTTTATGCATTTGT
ATTCCATGATTCAGTTCAAACTCTAAATAAGTAAACAGAGAGAGAAGTGATTCTCAGCTTTTAGGTGCGGAAAATGAGAGTTTTGG
AAGCTGAGAGTGGCATTTTCGTAAAATGGGCGAACACTCAATTCAGCAATATGGTTCTCAGGCTTAGATAGAGTTGGGATTGGAAA
TTATTGCAGCCAGAAGTTAAAAGTTTCTCAGGA
>RgMenA1 Rehmannia_glutinosa OWAS-2001018 (1KP)
CGCCTTTGGCTATGGCAGCAGCCACGTTCTGTTCCATAAGCCATGGCTATGGCGTCAACAAACTCAATCAACACCTTCTCAACAGA
CACAATTTCAACAGGATTTATCAAGTAATACCTCTTCCAGATGCCAGTCGAAGGCAACTACGCATGGAAATGTGTTTCGACAAGAC
GATCATAAGGCAATTACATTGTATACGAAGGCATTATGGTATTTTGTTGAAGTGTAGATCAGAGCACAATGACAGTCATGTTGTAG
AAGAGAAGGAAGAAAGTATCTCTAAAGCAACCTTAATATGGAGAGCCATCAAATTACCAATGTACACTGTTGCATTGATCCCTATA
ACAGTCGGAAGTGCAGCAGCTTATTTACAGACAGGCCAGTATTTCGGAAAGCGTTATTTAATGCTATTGATTTCTTCAGTCCTCAT
AATAGCTTGGCTCAACTTAAGCAATGACGTTTACGATTTCGAAACTGGAGCGGATAAAAACAAGAAAGAATCGGTTGTTAATCTAT
TTGGCAGCCGGACAGGAACTCATGTTTTTGCTTGGTTACTACTTGCACTTGGTTTCACGGGCCTTACTTGGGTTTCTGTGGAGGCT
GGAAGTCTACGTTCAATATTTCTACTTGGTTGTGCCGTATTTTGTGGCTACATTTACCAGTGTCCGCCGTTTCGTTTGAGTTACTT
GGGACTGGGAGAACCCTTATGCTTTGCAGCATTTGGTCCATTTGCGACCACCGCCTTTTACTTGCTTCAGAGCGGTACAAGGGAGC
TCTCGATTTCTGTCACCATTGTTTCTTCATCAATTCTTGTTGGTTTAACGACATCCTTGATCCTCTTCTGCAGTCATTTTCATCAG
ATAGACGATGACAAGGCTGTCGGAAAATTTTCGCCTTTGGTCAGGCTTGGATCTGACGGAGGTGCAAAAGTAGTGAAAGTGGCTGT
CGGGACGCTCTATTCGCTTCTATTTATTCTAGGACTTAGCCAAACTCTTCCTTTTTCATCTATTGTTTTATGCGCGTTAACGTTAC
CCGTTGGAAAATTAGTAGTTAGCTTTGTTGAGAAGAACCACAAGGACAAGACGAAGATCTTCATGGCAAAATATTACTGTGTGAGA
TTGCACACAGTATTCGGAGCTGCATTGGCTGCTGGGATGGTGGCTGCTAGAATGTTTGCAAGAAAGCAACTTCCTCATGCTATTAT
TCTTTGAACTTTCCGAGTCGAATAAGGAATTAATATTCTCAGAGTATCCAAGCTCCATACAATTGGTGGTGTTTGATCGAATGAGT
GACGATATTATTTGAAATAGAGGATCATGTGGAGGACTAATGGTGCTGTCCCATTTTTCTCTCTTTTTTTGGGGGCTTTAATTTTG
TATTTCAGTTTATGCATATGTAGATTTAGGAGGGCC
>RgMenA2 Rehmannia glutinosa assembled from SRR999317
CTTTAATTTATTTGGCCTTTTAATTGAAACAATGGCAGGAACCAGCTTCAGTTCGTCTGACAATCCAAGCAAGGAAAAAAATGGTA
AAAGATCAAAGAAGGAGACTACTTCCCATGACGAGAAAAAAGAAGAAGATATAGGTAGAGCAACTTTAATATGGAGAGCTGCCAAA
TTACCTATGTATACTGTTGCATTGATTCCTCTTACTGTAGGAACTGCAGCTGCATATTGGGAATCAGGCTACTATTCTTTGGAGCG
TTATTTCATTCTCTTGGCATCTTTTGTTCTTGTCAATCTTTGGGTCAATTTAAGCAACGATGTATATGATTTCGATACAGGAGCTG
ATATAAACAAAAAAGAATCTGTTGTCAATATAGTTGGCAGTCGCAATGCCATCCATATTCTTTCGTGGTTATTACTTGCCATTGGT
TTTGCTGGCCTTTCTTGGGTCGGAGTCGAGGCTAAAAACCCGCGTGCTATACTCTTATTGGCTGCGGCAGTCTTTTGTGGCTACAT
TTACCAGTGCCCACCGTTTCGTTTAAGCTACCACGGTCTGGGAGAGCCCTTATGCTTTGCATCGTTCCGCCCATTTTCCACCGTTG
CATTTTATTTGCTCCAGAGCAGATCAAGTGAGCTACCGATATCTAGCACGATTATATCCGCAGCAATTCTTGTCGGTTTTACATCA
GCCTTGATTCTCTTCTGTAGTCATTTCCATCAGATAGAGGATGATAAGGCTGTCGGAAAAATTTCGCCTTTGGTGAGGCTTGGAAC
TGAAACAGGATCAAAAGTAGTGAAAATGGCTGTTTTGGGGCTGTATTGGCTTGTTTTTGGTTTAGGACTGGCCCAAACACTTCCTT
ATCCTTGTATAATGCTCTGCATAATGACACTACCCATGGGAAACTTAGTAGTTAGCTTTGTTCAAGAGAACCACAAGGACAAATCG
AAAATTTTCATGGCTAAATACTATTGTGTGAGGTTACACACTGTGTTCGGAGCTGCTTTGGCTGCTGGACTTGTGGCATCAAGATT

AATACAATCTGGAGAGCAACTTCAGAGTCTATCGACGTACTTTGATCGTGCCAAGTTTGCGTATTTCTGAGTTTAATAGTATATTC
AGAAGTCCAACCTTTTGCTGGGAA
>LpMenA1 Lindenbergia philippensis ZVFS-2005540 (1KP)
GATTTAGTGATCTCAAACAAAATGGGTTTTTTCATAATTCTTGATCCTTAATTCCTTATCCCAATTTTTCTTCCATTATTTACATT
TTCTTCAAAAAAAAGAAAAATTGATTGGCGCCTTTGGCTATGGCAGCAGCCACTTTCTGTTCCATAAGTCATGGCTATGGCGTCAG
CAGAGTCAACGACTATCTTCTCAACAGACACGGTATCAACAGGATTGATCAAGTATTACCTCTTCCAGATGCCAGTCGAAGGCGAA
AACTTGTGAAAAAAGATTTCAACAAGACAGTCATCAGGCAATTACATTCCATACGAAGGCATTATACAATGTCGTTGAAGTTTAGA
GCGGAGAACAATGGCGATCGTATCGAAGAAGAGAAGGAAGGAAATGTCTCGAAAGCGACCTTAATATGGAGAGCCATCAAATTACC
GATTTACACTGTTGCATTGATCCCTATAACAGTTGGAAGTGCAGCAGCTTATTTGCAGACAGGCCAGTATTTTGGAAAGCGTTATG
TTATGCTCTTGATTTCTTCAGTTTTTGTCATAGCTTGGCTCAACTTGAGCAATGACGTGTATGATTCCGAAACTGGAGCGGATAAA
AACAAGAAAGAATCCGTTGTTAATCTATTTGGCAGCCGGACAGGAACTCATGTTTTTGCTTGGTTACTACTTGCACTCGGCTTCAC
GGGCCTTACTTTGGTGTCTGTCGAGGCTCGAAGTCGACATTCTATATTTCTACTAGCTTGTTCCGTCTTTTGTGGCTACGTTTACC
AGTGTCCACCATTTCGGTTGAGTTATTTGGGACTTGGAGAACCCCTTATGCTTTGCAGCATTTGGTCCGTTAGCTACCATCGCCTTT
TATTTGCTTCAAGGCGGTACAAGGGAGCAGCTATCAATTTCTTGCGCTGTTGTTGCTTCATCAATTCTTGTTGGCTTCACAACATC
CTTGATACTCTTCTGTAGTCATTTTCATCAGATAGAGGATGATAAGGCTGTCGGCAAATTTTCACCTTTGGTCAAGCTTGGAACTG
AAGGAGGTGCAAAAGTAGTGAAAGTGGCTATAGGGACAATTTATTCGCTTCTATTTATTCTCGGACTTAGCCAAACACTTCCTTTC
TCATCTATTGTTCTCTGTGCTTTAACGTTACCTATGGGAAAATTTGTTGTTAGCTTCATCGAGGAGAATCACAAGGACAAGACGAA
GATCTTCATGGCAAAATATTACTGCGTGCGATTGCACACAGTATTTGGAGCTGCATTGGCTGCTGGGCTGGTGGCAGCTAGAATGT
TAGCAAGAAAGCCCCTTCCTTATGCTGTTATTCTTTGAATTTTTTAAGTTACGTAAGGTGATGGGGATTGTTATCTCGATTAATGT
TCAGTTTTTTTTTTTCTCTTCTTCTAACAAACAGTATGCAAATCTGGCATTGTTCAACATATGTTATCTTTCCCTTACACGACATA
ATTTTTAGGGTGAATTACAAAAACTACATGTGAGGTTTTTGATCTAGTGTTTAAAGTAGGAGGTTAGCTGTAATATTTTTATGAAT
GTAAGGTGGTATTTTGAAACATTTTTTCAGTTATAAAAGGATAAATGTAAGACAAAAACAAATCATGG
>LpMenA2 Lindenbergia philippensis EJCM-2018390 (1KP)
TTGCACCTTAATTAGCTAGGTTTTTAACATGATGGCAGGAACCAGCTTGAATTTGGTAGAGAATGCAAGCAAGGAAAAAAATATTA
AAAGGTCAAAAAATGGGAGTACTAGTCCAAATGAAGAGAAAAATGAAGAAGAAATATCAAGAGCAACTTTGATATGGAGAGCTGCA
AAATTACCCATGTACACTGTTGCATTAATTCCTCTAACAGTGGGAACTTCAGCTGCATATTGGGAGTCAGGCTATTACTCTTTGGA
GCGTTATTTTATTTTATTGGCTTCTTTTGTTCTTGTCAATGTTTGGGTCAATTTAAGCAACGATGTTTATGATTTTGATACGGGAG
CTGATAAAAACAAAAAGGAGTCTGTTGTCAATATAGTTGGCAGTCGCACGGCCATCCATATTCTTTCATGGTTAATACTTGCGCTT
GGTTTTGCCGGCCTTACATGGGTTGGAGTCGAGGCTAAAAACCCGCGTGCTATACTGTTGTTGGCTTCAGCAGTCTTTTGTGGCTA
CATTTACCAGTGTCCACCGTTTCGTTTAAGCTACCATGGACTGGGAGACCCCTTATGCTTTGCGGCGTTTGGTCCTTTTTCCACCG
TTGCATTTTATTTGCTACAGAGTAGTTCAAGTGAGCTACCGATATCTAGCACGATAGTTTCCTCAGCAATTCTTGTTGGTTTTACA
TCAGCCTTGATCCTCTTTTGTAGTCATTTCCATCAGATAGAGGATGATAAAGCTGTTGGGAAAATTTCTCCTTTGGTGAGGCTTGG
CACTGAAAAGGGATCAAAAGTAGTGAAGATGGCTATTTTGGGGCTCTATTGGCTCGTGCTTGGTTTAGGACTCGCCCAAACTCTTC
CTTATGCTTGTATTGTCCTATGTGCTATGACACTACCCATGGGAAATTTAGTAGTTAGCTTTGTTCAAGAGAACCACAAAGATAAG
TCGAAAATCTTCATGGCTAAATACTACTGTGTGAGATTACACACTGTATTTGGAGCTGCTTTGGCTGTTGGATTAGCGGCATCAAG
AATCAATGGTGGGGATTATCTTCAGAGTCCATCAACGTATTTTGATCGTGCCAAGTTTGCGTATTTCTGAGTTTAATTATTAATTA
GCACTCATTTTCTTATATCTACCATCAGATACATCTTGAACTACATTTATTCTTGTTATAAAAGAG
>VtMenA1 Verbascum Thapsus XXYA-2074947 (1KP)
AAAAGCATTACCGTAGTTTTTACTCAAGGCATTGAAGTTTCAGCTGGTGACATAGACTATGGCAGCAGCTAATTTCTGTTCAATGA
GCCATGGCTATGGCGTCAAAAACCTCAACCAGTTCCTTCCAAGAAGATGTAATAATTACAGGGCTTATCAAGAATTATCTCCTCAA
GATGCCGTTCGAATGCCATTATGCAGTAAAACGTATTTCGACAAAACAATCATAAGGCGATTACATTCTAAGAAACCGCATCACAG
AATTTTGTTCAAGTGTGCAGCAGAGCTCATTGACACTCACGGTGAAGACGAAAAGGAAGAACATATCTCGAAAGCAACTCTGATAT
GGAGGGCCATCAAATTACCAATATATACTGTTGCATTGATTCCTATAACAGTAGGAAATGCAGCAGCTTATTTGCAGACAGGCCAG
TATTATGGAAAGCGTTATGTTATGCTATTGGTCTCTTCATTTCTCATCATAGCTTGGCTCAACTTAAGCAATGACGTTTACGATTT
CGATACTGGAGCAGATAAGAACAAGAAAGAATCGGTTGTTAACTTAATTGGAAGTCGAAAAGGAACCCGTGCTTTTGCATGGTCAC
TACTTGCACTTGGTTTCATGGGCCTTACAAGGGTGTCGATGGATGCTGGAAGTTTTTACCCTTTATTTCTACTTGCTTGTGCCATA
ATTTGTGGCTACATTTACCAGTGCCCGCCATTTCGGTTGAGTTACATGGGATTGGGAGAACCGTTATGCTTTGCAGCGTTTGGTCC
ATTTGCCACCACAGCCTTTTACTTTCTTCAGACAGGCACAAGGGATCTGTCGATATCTGCCACTGTCATTTCTTCATCGATCCTTG
TGGGTATTACAACATCCTTGATCCTCTTCTGCAGTCATTTTCATCAGATAGACGATGATAAGACTGTTGGAAAATTATCCCCTTTG
GTTAGGCTTGGAACTGAAACTGGCGCGAAAGTAGTGAAAATCGCTGTAATAACACTCTATTCGCTGCTATTTGTATTAGGACTCAG
CCAAACTCTTCCTTTCTCATCCATTGTACTCTGTGCTTTAACATTACCGATAGGGAACAAAATAGTTAGATTCGTTGAGGAGAATC
ACAAGGACAAAACGAAGATTTTCATGGCGAAATATTACTGTGTGAGATTACATACTGTATTCGGAGCTGCATTGGCTGCCGGGCTG
GTGGGGGCTAAAATGTTCGCCCGAAAAAAAAATACCACATGCTATGATCATCTCATAACGGCGGCCCGAATACTC

>PaMenG2 Phelipanche aegyptiaca Pa.c176185_g1_i2.37080
TACACGACGCTCTTCCGATCTCAGGGGACCGAAAACCGAAAATGGCTACAGCTACACTTAGACGCCGAACAGGAACACAGACCGCA
ACCCATCAGGGCGCAGCTGAACGCCAGGAGCTCTTTAACCGCATTGCCCCGGTCTATGACAAATTGAATGACGTGTTTAGCCTGGG
CTTGCATAGATTATGGAAGAGGTGGTCTATCTCTTGGAGCGGTGCAAAAGAAGGTGACAAGGTGTTGGATGTGTGCTGCGGAAGTG
GGGATTTTAGTTTTCGGCTGGCTGAGAAAGTTGGGATCAATGGCAAGGTCATTGGTCTTGATTTCTCCAAGGAGCTATTACAGGTC
GCTGCATGTCGTCGGCGTGACCAATCAAGTTCAAAGCCGTGCTACAACAACATTGAGTTCATTGAAGGAGACGCAGTTGCTTTGCC
TTTTCCCGACTCTACTTTTGACGCCGCTACGATTGGTTACGGGTTAAGAAATGTCATTGATCGGAAAAAAGCTCTCGAGGAGATGG
TCCGAGTTCTTAAACCGGGCGCAAAATTATCTGTTCTTGACTTTAACAAAAGCACTCATTGGCTAAACATTAAAATTCAGGAATGG
ATGATTGATTATGTAGTGGCTCCAGTTGCAAGTTGGTATGGACTTGAAAGTGAGTACAGATACTTGAAGATTTCTGTCAAGGAATA

TCTCACAGGAAGTGAGTTGGAGAAGATGGCTTTGGAAGCTGGTTTTTCCGTTGCCAAGTTCCATTCGATTGCTGGAGGAGTCATGG
GAAATTTGGTCGCCACGCGCTAGAACATATTCCTCGTTCCGTTCTATTTCATCATTCTGGTGTTCGCTTTCTGCAATTTCTGAAAT
GCTTTGCGTTCGTGTTTGTTTTTGTTATCACATTATTATGATTTTCGATGATTGTGGCCCGATCCAGTCGATCTTGTTAAAATTTC
TATCAGAAATAAAATTTCTTTGAGCTTTTGTGATCCATTGTTTTTGTTATCAGAAATAAAATTTCTATGAATCTGTCTGTTGAAAT
GTTG
>ShMenG1 Striga hermonthica Sh.c17385_g1_i2.603
AAAAAATGGAAAATCTTGAGGGTAAATTAGACCGTTCACTTGCTCAGAACTGAAAAGGCTATGGCTACTCTTCACTTCACTCTCCC
GTCTACAACCGGCCGGCAGCGGCCGCCGGAATTCCGGTCCATCTTCAAACCGGCTCGGTGTGCAGCTGAGAGACAGGCCCTATTTG
ACCGAATTGCCCCTGTCTATGATAACTTGAATGATTTGTTGAGTTTAGGGGGCCACAGGGTATGGAAGAGGATGGCCGTTTCTTGG
ACTGGAGCAAAAGAAGGAGACACTGTGTTGGATGTGTGTTGTGGGAGTGGGGATTTGGCCTTTCTATTGTCTGAAAAAGTTGGAGT
CAAAGGCAAGGTTTTTGCTGTTGATTTTTCGAAGGAGCTATTACAGGTTGCTGCATCTCGACAGCTTAAGCGGTCAAAAGCGTGCT
ACAAAAACATCGAGTGGATTGGAGGAGATGCGGTCGATTTACCCTTCTCTGGCTCGTATTTTGATGCTGCTACCATTGGATATGGG
TTGAGAAATGTGGTAAATAGGAAAAAGGCCCTGGAGGAAATGAGTCGGGTACTAAAACCCGGTTCGAAACTATCTGTTCTTGATTT
CAACAAAAGCACTAACCCGTTAACGAGCTCAGTCCAGGATTGGATGATTGACTATGTAGTGGTGCCCGTTGCTAGTGGGTATGGCC
TAGCAAGTGATTATAAGTACCTGAAGAACTCAATCAAGGAATATCTCACGGGAGATGAGTTGGAGAAGGTAGCTTTAGAAGCGGGC
TTTTCCCGGGCCAGACATTATGAGATTTGTGGAGGGTTAATGGGTAATTTGGTGGCCACTCTCTAAGAATGTTCTTATTGTTCTTT
TAATAACGTTTTTAAGAGTCGTAAGTGTGATATGTTTGTTCTGAAATATAGTTTAGCTTTTTTTGTAACATTTAGAGTGCTTATAC
TCCCTGTTTCACGATATGAGGTTCTAGAATTTTTGCACATGGAATTTAGTCCAGAAATGGAGTTTATCATGATGTCGATAAAACTT
GATTTTAAAAAGCTTCGCTCATAATTTGATTCCAATAAACCGTATTAATAAGCAAGAAAACTGCATAATAAATTATCAAGAAATAT
AATTATTCTGTTTGCATATATGCCTATAGTTATATTGTCTGTAAGTATTCCTTTCGCTATACATATTCTTTTCTACAACTCTATTG
TACCTTTTATAAATAAAAAGCCGAACTTCCTACATCACCTAGGATAGAAATTGCTGTGGGACATATTAGCAATAGAATCACCATAG
CTGTACTCGAGGGGAGATACCGTTTTTGCCCTTGAGTCGTTAAGTTCCATTGTTATCTCATTCATGAAAGGAGGATTAGTCGGGCT
CGGTAATTCTTCTTTATCCACAAGCATTTGTAAAACCTTCGACATTGATGGCCTTAACGAGGGGTTTTCTTGCATGCACAAGAGCG
CCGCGTGGACTACACGCAACATCTCGTTTTTTGCATTGATATTGTTGTAGTTTTGCTGCATTAG
GTTCGGGTCGAG
>ShMenG2 Striga hermonthica Sh.c18886_g2_i1.0
AGATCAAAAGGCTCAAATGTTGGGTGGCCCTTAACAGATTTTCATCACAAATTGGATACATATGGCCACACTTAGACGCCGATCTG
AGCCACAGCCGCCGAGCCATGAAAGTGCAGCCGAGCGCCAGGAGCTTTTTAACCGCATAGCACCTGTCTATGATAAACTGAATGAC
TTGTTTAGCTTGGGATTGCATAGATTGTGGAAAAAGTGGTCTATTGCATGGAGCGGGGCAAAAGAAGGAGACAAGGTGCTGGACGT
GTGTTGTGGCAGTGGGGATTTGAGTTTTCTGTTGTCCCTCAAAGTCGGGATCGACGGCAAGGTCATTGCTCTGGATTTCTCAAAGG
AGCTATTAGAGATTGCGGCATCTCGCCAGCGCGAGTGGTCCAAGTCGAAACCGTGCTACGAAAACATCGAGTGGATTCATGGAGAT
GCAGTTGCTTTACCGTTCCCCGACTCGAGTTTTGATGCTGCCACCATCAGCTATGGGTTGAGAAATGTGGTGGATAGGAAAAAGGC
TCTCGAGGAGATGGTTCGGGTTTTGAAACCCGATGCCAAGTTATCCGTTCTTGACTTCAACAAAAGCACTAATCGGCTGACCTCTA
AAATCCAGGATTGGATGATCGACTATATAGTGGTTCCAGTGGCGACCTGGTATGGGCTTGCAAGCGAGTATAGATACTTGAAGAAT
TCAATCAAGGAGTACCTCACAGGGAAAGAGTTGGAGAAGTTAGCTTTGGAAGCAGGCTTCTCTGAGGCCAAGCACTATGAGATTGC
TTGTGGATCCATGGGGAATTTGGTCGCCACACGCTAGAAATTTGATTGTTCCACTTTTACTGTTTCTCTTGTGTTTGTTATATTCT
GCAATTTTCCAAATGCTTTCATTTTTGTCATTTTTTTTTTCAACCATTTAATGATTTCTGATGATCTTTTTGGTCTAAAGAACCAT
ATGCTAGAATCCCATATGCTAAAATTTCCTCTCCGGCCTCCGTACCTTTATACGGATGGACCAATTGGAATACTATTCATTCATAT
TTGTGTAATCG
>TvMenG1 Triphysaria versicolor Tv.c7280_g3_i7.0
AAATAATGTTGTTTACAAAGAACTGAAAAGAAAATGATTTCACTTCAATTCACTCTTCCCTCCATAACCAGCCGGCGAAGTTTACC
GGAATCCCGGCCCATTCTCAAACCTATCCGCTGCGCATCTGACCGTCAGGCGCTTTTTAACCGCATTGCCCCTGTCTATGATAACC
TGAATGATTTGCTTAGCTTAGGAGCACATAGAGTATGGAAAAGGATGGCTGTTTCTTGGAGCGGAGCTAAAGAAGGTGACACTGTG
TTAGATGTGTGTTGTGGGAGTGGAGATTTAGCTTTTCTCTTGTCTGAGAAAGTTGGAACCAATGGAAAGGTTTTTGCTCTCGATTT
CTCAAAGGAGCAATTATATATCGCGGGATCTCGACAAAGCAAACGATCGAAAAATTGCTACAAGAATATCGAGTGGATTGAAGGAG
ATGCGGTTGATTTACCATTCTCTGCCTCGTTTTTCGATGCTGCAACGATTGGTTATGGTTTAAGAAATGTATTAGATAGGAAAAAA
GCTCTTAACGAAATGTGTCGGGTTCTAAAACCGGGTTCGAAATTATCTGTTCTTGACTTCAATAAAAGCACTAATCCATTAACCTC
TTCAATTCAGGATTTGATGATCGACTATGTAGTTGTCCCCGTTGCTGACGGGTATGGCGTAGCCAGTGATTATAAGTATTTAAAGA
ACTCAATTCAAGAATATCTCACAGGGAACGAGTTGGAGAAGCTAGCTTTAGAAGCGGGCTTTTCTCGGGCCAAGCATTACGAGATT
AGTGGAGGGTTAATGGGTAATTTAGTCGCTACTCTTTAGAAATTTATTCATAGAAAAAAGAACAAAAATCGTGTCTTTATTACTTG
CTTCGTGTTTGAATAAATATAT
>TvMenG2 Triphysaria versicolor Tv.c6701_g1_i1.0
GCTTGTACGAAATCAAAGTGCTTAAACAGATCCAATGTCGACGACGCTTCTTCGCCGCCGATCGGAGTCAGAGCCGGCCAGCCATG
AGGGCGCAGCTGAGCGCCAGGAGCTTTTTAACCGCATTGCACCTGTCTATGATAAATTGAATGATGTGTTCAGCTTGGGTTTGCAT
AGATTATGGAAGAGGTGGTCTATTTCTTGGAGTGGGGGAAAAGAAGGAGACAATGTGTTAGATGTGTGCTGTGGAAGTGGGGATTT
GAGTTTTCTGTTTTCTCAGACAGTTGGAGTCAATGGCAAGGTTACTGCTCTGGATTTCTCGTCGGACCAACTGCAGGTGGCTAAAT
CTCGCCAGCACGAGCGATCAAAATCAAACCCGTGCTACAAGAACATTGAATGGATCGAAGGCGATGCGGTTGCTTTGCCATTTCCG
GACTCGACTTTTGACGCAGCTACGATTGGTTACGGGTTAAGAAATGTGTTGGATAGGAAAAAGGCTCTGGAAGAGATGGTTCGTGT
TCTTAAACCGGGTGCCAAATTATCTGTTCTTGACTTCAACAAAAACACTAGTCAAGTGATGTGTAAAATTCAGGATTGGATGATTG
ATTATGTAATTGTTCCGGTTGCAAGTTGGTATGGACTTGCAAGTGAGTACAGATACTTGAAGAACTCAATCAAGGGATATCTGACA
GGAAGTGAGTTGGAGAAGCTAGCTTTGGAAGCCGGTTTTTCTGAAGCCAAACACTATCCGATTGCCGGAGGAGCAATGGGAAATTT
GGTCGCCAAGAAATAGAAACGGGTTTTGTTGTTCTTGTTGTTTCAATAATTATCTGTTTGTTGTTGTTGTTGTTGTCGTTGTTTTA
ATAATAAATTGTAG

>OfMenG2 Orobanche (syn. Aphyllon) fasciculata PHOQ-2096404 (1KP)
TCATAGTTGACTCCTAAAAACAATGGAACTTGTGTAATATTAAAATAAACATTTACTCTCTTACAGCATAGATAAGATATACAAAC
ATTTATATATTGCTCGGCCTTTGGGACCGTACCAGAGACTGTAAATCGAAATGGCTACACTTAGACGCCGATCAGAAGCACAGCCT
TCAAGCCATGAGGCCGCAGCTGAGCGCCAGGAGCTTTTTAACAGCATTGCACCGGTCTATGATAAATTGAATGATTTGTTCAGCCT
GGGCTTGCATAGATTATGGAAGAGGTGGTCTATTTCTTGGAGCGGGGCAAAAGAAGGTGACAAGGTGTTGGATGTGTGTTGTGGAA
GTGGGGATTTGAGTTTTCTGCTGTCTGAGAAAGTTGGGATCAATGGCAAGGTCACGTCTCTAGATTTCTCAAAGGAGCAATTGAAG
ATCGCTGCATCTCGCCAGCGTGAGCGATCAAAGTCAAAGCCCTGCTACAACAACATTGAGTGGACCGAAGGAGATGCAGTTGCTTT
GCCGTTTTCCGACTCCACTTTTGATGCTGCTACCATTGGCTATGGGTTAAGAAATGTTATTGATAGGAAAAAAGCTCTTGAGGAGA
TGGTTCGGGTTCTAAAACCGGGTGCAAAATTATCTGTTCTTGACTTTAACAAAAGCACTAACTGGCTAAACTCCAAAATTCAGGAT
TGGATGATTGATTATGTAGTGGTTCCAGTTGCAAGTTGGTATGGACTTACAAGTGAGTACAGATACTTGAAAAACTCCATCAAGGA
ATATCTCACAGGAACTGAGTTGGAGAAGCTAGCTTTGGAAGCCGGTTTTTCCGTTGCCAAGCACCATGCGATTGCTGGAGGATCCA
TGGGGAATTTAGTCGCCACTCGCTAGAAATGTTTTCGCTCCATTCTATTGCAAGGACATGTTCCTCATTCTCTTTCTGCAATTTCC
GAAATGCTTTGTTTTTATTGTTGTTATCGTTGTCATGATTTCCGATGATTATGGCTGATCCGATGAACCTTGTTAAACTATCGGAA
TTTAGTCGGTTTGAGTTTATGTAATAATAGTCAGAGTAATGGTTTATCCATATGAAGTCTCTTTGTTGAACTGTTAACTTTTGCAT
CTTTGAGTGCTCAATATGTATTGAACTTTGTGTGTCTGTTGTATCTTATATTCACTTGTTCAATTAGTCCCGGCCTAGGTGCTGGC
CTGTCCGTAGCCTC
>CaMenG2 Conopholis Americana FAMO-2024401 (1KP)
TTGAGACTGAAAATCGAATGGCCGCTACACTTAGACGCCGATCAGAAGCACCTGCACAGCCCGCTGCCGCAAGCCATGAGGGCGCA
GCTCAGCGCCAGGAGCTTTTTAACCGTATTGCACCTGTCTATGATAAATTGAATGATTTGTTTAGCCTCGGGTTGCATAGATTATG
GAAAAGGTGGTCTATTTCTTGGAGCGGGGCAAAAGAAGGAGACAAGGTGTTGGATGTGTGTTGTGGAAGTGGGGATCTGAGTTTTC
TGATGTCTGAGAAAGTTGGAAGTGGCAAGGTTAATGCTCTAGATTTCTCAAAGGAGCAATTACACATTGCTGCATCTCGCCAGCAT
GAGCGATTGAATTCAAAGCCGTGCTACAAGAACATTGAGTGGTTTGAAGGAGATGCAGTTGCTTTGCCATTTTCGGACACAATTTT
TGACGCAGCTACGATTGGGTATGGGTTAAGAAATGTGGTGGATAAGAAAAAAGCTATGGAGGAGATGGTCCGAGTTTTGAAACCGG
GTGCCAAATTATCTGTTCTTGACTTTAACAAAAGCAACAATTGGCTAAACTCTAAAATTCAGGATTGGATGATCGATTATGTAGTT
GTTCCGGTTGCAAGTTGGTACGGACTTGCAAATGAGTACAGATACTTGAAGAACTCAATCAAGGAATATCTCACAGGAAAGGAGTT
GGAGAAGCTAGCTTTGGAAGCAGGTTTTTCTGAAGCCAAGCACTATGAGATTGCTGGAGGATACATGGGAAATTTGGTCGCCATTC
GCTAGAAATTTATACTTTTGTTCCCATTTTGCTTCCCTGATTACAATGCAAGGAGGACATGTTCTTCATTCCTTCCTTCCTTCGTA
TGC
>RgMenG1 Rehmannia glutinosa OWAS-2002674 + OWAS-2025114 + OWAS-2044184 (1KP)
GTGGTTTTTACCAAGAACTCAAAAAATAGAATGGCTACACTTCAATTTACTCTTCCCTCCATAACCGGCGGGCGACGTCTACCGGA
ATCCCGGTCCACTATTAGACCGGTTCGGTGCGCAGCTGAACGGCAGGCGCTTTTTAACCGCATTGCCCCTGTCTATGATAACTTAA
ATGATTTGTTTAGCTTAGGAAGCCATAGAATATGGAAGAGGATGGCTGTTTCTTGGAGCGGTGCAAAAGAAGGAGATACCGTGTTG
GATGTGTGTTGTGGAAGTGGGGATTTGGCTTTTTTGTTGTCTGAGAAAGTTGGAATCAATGGCAAGGTGATTGCTCTTGACTTCTC
AAAGGAGCAATTGACAATTGCTGCATCTCGCCAGCGCAAGCGATCGAGGGCGTTCTACAAGAATATTGAGTGGATTGAAGGAGATG
CAGTTGATTTACCATTCTCCGACTCCTCTTTCGATGCTGCAACCATTGGCTATGGGCTAAGAAATGTAGTAGATAGGAAAAAGGCC
CTGGAGGAGATGTGTCAAGTTCTGAAACCTGGTTCAAAAGTATCTGTTCTTGACTTGAACAAAAGCACTAACCCATTAACCTCTTC
AATTCAGGATTGGATRATTGACTATGCAGTGGTTCCTATCGCCAACGGGTATGGCGTAGCAAGTGATTATCAGTACTTGAAGAACT
CTATCAAAGAATATCTCACAGGAAATGAATTGGAGAAGCTAGCTTTAGAAGCAGGTTTTTCTAAGGCCAAACATTATGAGATTGGT
GCGGGGTTCATGGGAAATTTGGTAGCCACCCTTTAAGGAATGTTTTCATGCAGTCTCCTTCTTCTCAAGGCAGGTGCATTTTCGTG
TGCTCGCTTTCTGCAACTTCTAGAAGGTTCTATCTCTCTTTCTCTTTCTCCATCTGTATATGCTGA
>RgMenG2 Rehmannia glutinosa OWAS-2072104 (1KP)
GTATTTCTAATACATATTACCCAGATTCTTACATCTTAGATTAGTTTCTACACTTATATATTGCTACTCTAGTAGTGTTGAAAGAA
ATCGAGGGACTTTAGCAGAGAGTGAAAATCGAAATCGAATGGCAACACTTAGACGTCGATCGGAACCACAGACCGTAACCCATGAG
AGCGCAACTGAGCGCCAGGAGCTTTTTAACCGCATTGCTCCTGTCTATGATAAATTGAATGATTTGTTTAGCCTGGGGCTTCATAG
ATTATGGAAGAGGTGGACTATTTCTTGGAGCGAGGCAAAAGAAGGAGACAAAGTGTTGGATGCGTGTTGTGGAAGTGGGGATTTGA
GTTTTCGGTTATCTGAGAAAGTTGGAATCCATGGCAAGGTGATAGCTCTAGATTTCTCAAAGGAGCTATTAGAGATAGCTGCATCT
CGCCAGCGTGAGCATGAGCGATCAAAGCCATGCTACAAGAACATTGAGTGGATTGAAGGAGATGCAGTTGCTTTGCCATTTTCTGA
CTCCACTTTTGACGCTGCTACCATCGGCTATGGGCTAAGAAATATTGTTGATAGAAAAAAAGCTCTGGAGGAGATGCATCGAGTTT
TAAAACCGGGTGCAAAATTATCTGTTCTCGACTTTAACAAAAGCACTAATTGGCTAAACATTAAAATTCAGGAATGGATGATTGAT
TATATAGTGGTTCCTGTTGCAAGTTGGTATGGACTTGCAAGTGAGTACAGATATTTGAAGAACTCAATCAAGGAATATCTGACAGG
ACCTGAGTTGGAGAAGCTAGCTTTGGAAGCAGGGTTTTCTGAAGCTAAATTCTATGAGATTTCCAATGGATCCATGGGAATTTTGG
TAGCCAAGCGCTAGACACATTTTCTGTCCACATTCTGCTTCCCCGACAGGTTTCCGTTGTTGTTGTTGTTGTTGTTGTTGTTGTTT
ATGATGGTTATGTTGATCCAATGTAGCTTGCTAAAG
>LpMenG1 Lindenbergia_philippensis EJCM-2011304 (1KP)
AGCCATCTAAATTGTTTTCTTTGCACGGTCGATTTTCTGTATAAAACTCGAAATACAGTGGAGTTCTTCTAGAACTGAATTGAAAA
TGGCTACACTTCAATTCACTCTTCCCTCCATAACCGGCGGCCGACGTCAACCGGAATCCCGTTCCATTCTTAGACCGGTTCGCTGC
GCAGCTGAACGCCAGGCGCTTTTTAATCGCATTGCCCCTGTCTATGATAACTTGAATGATTTGTTAAGTTTAGGGGCCCATAGAAT
ATGGAAGAGGATGGCTGTCTCTTGGAGCGGAGCCAAAGAAGGAGATACTGTGTTGGATGTGTGTTGTGGAAGTGGGGATTTAGCTT
TTCTGTTGTGCGAGAAAGTTGGGATCGATGGCAAGGTGATTGCTCTTGATTTCTCAAAGGAGCAATTACAGATTGCTGCATCTCGT
CAGCTCGCGCGATCAAAGGCGTGTTACAAGAATATTGAGTGGATTGAAGGAGATGCAGTTGATTTACCATTTTTCGACTCGTCTTT
TGATGCTGCAACTATTGGCTATGGGTTAAGAAATGTGGTAGATAGGAAAAAAGCTCTAGAGGAAATGTGTCGAGTTTTAAAACCGG
GTTCGAAATTATCTGTTCTTGACTTCAACAAAAGCACTAGCCCATTAACCTCTTCGATTCAGGATTGGATGATTGATAATGTAGTG
GTTCCTGTTGCCAGTGGATATGGCGTAGCAAGTGATTATCAGTACTTGAAGAACTCAATCAAGGAATATCTCACAGGGAATGAGTT

GGAAAACTTAGCTTTAGAAGCAGGCTTTTCTGAGGCTACCCATTATGAGATTGGTGGCGGTTTAATGGGAAATTTAGTAGCCACGC
TTTAAGTGTA
>LpMenG2 Lindenbergia_philippensis EJCM-2011303 (1KP)
CGTTGTAAAAAAATCGAGGGGCTTTAGAAGAGACTGGAAAATCGAAAATGGCTACACTACGACGTCGTTCCGAGCCACCGGCCGTG
ACCCATGACAGCGCTGATGAGCGCCAGGAGCTTTTTAACCGTATTGCTCCTGTTTATGATAAATTGAATGATTTGTTCAGCCTGGG
TTTGCATAGACTATGGAAGAGGCGGGTTATTTCTTGGAGTGGGGCAAAAGAAGGAGACAATGTACTGGATGTGTGCTGTGGAAGTG
GGGACTTGAGTTTTCTGTTGTCTGAGAAAGTTGGAGTCAATGGCAAGGTTATTGCTCTAGATTTCTCAAAGGAGCAATTACAGATT
GCTGCATCTCGTCAGCTCGCGCGATCAAAGGCGTGTTACAAGAATATTGAGTGGATTGAAGGAGATGCAGTTGATTTACCATTTTT
CGACTCGTCTTTTGATGCTGCAACTATTGGCTATGGGTTAAGAAATGTGGTAGATAGGAAAAAAGCTCTAGAGGAAATGTGTCGAG
TTTTAAAACCGGGTTCGAAATTATCTGTTCTTGACTTCAACAAAAGCACTAGCCCATTAACCTCTTCGATTCAGGATTGGATGATT
GATAATGTAGTGGTTCCTGTTGCCAGTGGATATGGCGTAGCAAGTGATTATCAGTACTTGAAGAACTCAATCAAGGAATATCTCAC
AGGGAATGAGTTGGAAAACTTAGCTTTAGAAGCAGGCTTTTCTGAGGCTACCCATTATGAGATTGGTGGCGGTTTAATGGGAAATT
TAGTAGCCACGCTTTAAGTGTA
>VtMenG1 Verbascum Thapsus XXYA-2018913 (1KP)
CCCTCCAATACAAAATTACTTCGTCCAAGTCCACTTCCCCCAAGTATGGGCATAAAAGCCAATGGAATTGGATGGGTTTTAGCAAA
ACCGGAAAAATCAAAATGGCCACACTCCAATTCATTCCTCCCTCCATCACCGGCCGGTCCAGTTCCCGGTCAATTCGACGTCCGGT
TCAGTGTTCGGCTGAACGGCAGGCGCTTTTTAACCGCATTGCTCCGATCTATGATAACCTGAATGATTTGTTGAGCTTGGGGAGAC
ATAGAGTATGGAAGAGGATGAATGTTTCTTGGAGCGGAGCAAAAGAAGGAGATACTGTGTTGGATGTGTGTTGTGGAAGTGGGGAT
TTGGCTTTTCTCTTGTCCGAGAAAGTTGGAACGAATGGCAAGGTGATTGCTCTTGATTTCTCGAGCGAGCAATTACAAATCGCTGC
CTCTCGCCAGCGTGAACGAACAAAGGCCTGTTACAAGAACATTGAGTGGATTGAAGGAGATGCAGTTGATTTACCATTCTCTGACT
CGTCTTTTGATGCTGCTACAATTGGCTATGGATTAAGAAATGTAGTGGATCGAAAAAAAGCTCTTGAGCAGATGTTTCGTGTTTTA
AGACCTGGTTCAAAGTTATCCATTCTTGACTTCAACAAAAGCAGTAGCCCATTAACCTCTACATTACAGGATTGGATGATCGACTT
TGTAGTAGTTCCTGTTGCTAGTGGATATGGCCTTGCGAGTGAGTACCAATACTTAAAGAACTCAATCAAGGAGTATTTGACAGGAA
AAGAGTTGGAGAAGCTAGCTTTAGAAGCGGGATTTTGTGATGCCAAATTTTTCGAGATTGAGGCTGGATTAATGGGAAATTTGGTA
GCCACCCGCTAAAAATGATTTTTGATCCAATTTGCTATATACAGTCTCAAATGCCATCTTTCAGCGAGTCGGGTACTTTTTTTACA
GTGTTTTTTCCCCTTCACATTGTAGAATAATAAACTTATATCATTTGGTTGCTCATTGTATGATATGATGAAATAAAAATGTTGAA
ATTTGTATTTATTAAACTCTGTCTCCCTTTCAATACATGTAATAAATGCATGAACTTAGAGCATGGATATAAAAAATGTATTTGGT
TCCCCCAC
>VtMenG2 Verbascum Thapsus XXYA-2014945 (1KP)
GACGGCATCTCCTTAATTAACTATTTCCGGATTATATATTATTGCTCGCCTCTGGAACAGATGAGATGAGTTTTAGTGCGTATAGA
ATGGCATCAATACGACGTCGTTCGGAGTCACAGCCCGCCGGAAAATTCGCCGAGGGCGCAGCTGAGCGCCAGGAGCTTTTTAACCG
CATTGCTCCTGTTTATGATAAATTAAATGATGTGTTTAGTCTGGGGATGCATCGATTATGGAAGAATTGGTCTATTTCTTGGAGCG
GGGCAAAAGAAGGAAACAGAGTGTTGGATGTGTGTTGTGGAAGTGGGGATTTGAGTTTTCGATTTTCTGAGAAAGTGGGCATCAAT
GGCAAGGTGATTGGTTTAGATTACTCAAAGGAGCTGCTACAAATCGCTGCAACTCGCCAACATAAGCTCGTGCAATCGAAGCCAAG
CTACAAGAACATTGAGTGGATTGAAGGAGATGCAACTAAGTTACCATTCCCTAATTCCTCTTTCGATGCTGCTACAATTGGCTACG
GTTTAAGAAATATTGTCGATAGGAAAAACGCTCTGGAGGAAATGGTTCGAGTTCTTAAACCGGGCTCCAAATTATCTGTTCTTGAC
TTCAATAAAAGCACTAGCAAATCAATGTGTAAATTTCAGGAATTGGTGCTCGATTATATAGTGGTTCCTATTGCAAGTTGGTATGG
ACTCAGGGATGAGTATCATTACTTGAACAAGTCGATCAAAGAGTATATGACAGGAAACGAGTTGGAGAAATTTGCTTTAGAAGCGG
GATTTTCTGAAGCCAAGTATTATGAGATTGCCGGAGGAGCCATGGGAAATTTGGTAGCAACGCGCTAGAAAATTTGGCGACTTTTA
GTCGATTCTT

>PaQR2 Phelipanche aegyptiaca Pa.c21330_g1_i1.5664
AGAGAGACGACCCGTACCCTTCCAATGGCGACCACACTTTCTATTGTTTACTATTCGACATACGGGCATGTTGAGAAACTTGCGCA
AGAGATCAAGAAAGGAGCAGAGTCTGTTGAAGGAGTTGAGGCTAAACTCTGGCAGGTTCCAGAAACTCTGTCCGATGAGATTCTTG
GGAAGATGGGTGCCCCGCCAAAGAGTAGTGAAGTGCCTGTAATAACCCCAGACGAACTTGCCGAGGCTGATGGTTTTATATTTGGT
TTTCCAACTAGATTTGGAATGATGGCTTCTCAATTTAAAGCTTTCTTTGATTCAACTGGAGGGTTATGGAGGACTCAGGCATTAGC
CGGCAAGCCTGCTGGCATTTTTTACAGTACTGGATCTCAAGGCGGGGGGCAAGAAACTACAGCCTTAACAGCCATTACACAGCTGA
CTCATCACGGTATGATATTTGTCCCAATTGGTTACACATTTGGAGCTGGAATGTTTGAGATGGAGAAAATCAAAGGTGGAGCCCT
TATGGTGCTGGGACATATGCCGGAGATGGTTCCCGACAACCTTCCGACATTGAACTTGCACAAGCTTTTCATCAGGGAAAGTACAT
TGCCGCCATCACCAAGAAGCTCAAAAAAATCTGTCTAATTTCACACTTCCTTATCAACATCTCCAATATTAATTCCATCCTACGAAT
TAAATATGCTGTTTTTCCTGTTTCGGTTATAAGTTTGCTTTGTTATGTCTTCGTCAAATTAAATTGTTATGTTTTTCAGTCGGAAT
AAATTGGTTTTCACGTACTTTATTTTATTTTATTGGGTTATGTCGATGTTTTTCACATATGCCGCAAACTTATCATAAAAATCAAG
AATCATACATCATCATTACTTTTCCAAAAAA
>ShQR2 Striga hermonthica Sh.c13828_g1_i1.4428
ACTGAATCTTCTTCTTCCATAACCATAACAGTATTGTCTATAAATAGCAGCCTTAATTCTCATTTACTTTCACAACACTACTTCAC
ACACCAAATCAAACTAAAAGAAACACACTTCATTATTACAATGGCTGCTAAAGTATACATTGTTTACTATTCGACCTACGGGCATG
TCGAGAGACTCGCGCAAGAGATCAAGAAAGGAGTCGAATCCATACCAGAAGTTGAAGCTAAACTTTGGCAGGTCCCGGAAACTCTC
ACGAACGATATCCTCGCGAAAATGGGGGCCCCACCAAAGAACAACGACGTGCCGGTTATTAGCCCGAACGAACTTGTGGATGCCGA
TGGCATCATATTCGGTTTTCCGACAAGATTCGGGATGATGGCCGCTCAATTCAAAGCATTTTTCGACTCGACCGGTGGCTTATGGA
GGACTCAAGCACTTGCCGGAAAACCTGCCGGAATCTTTTATAGCACCGGATCTCAAGGCGGTGGCCAAGAAACCACACCATTAACA
GCCATAACACAACTTACTCATCACGGCATGATATTCGTGCCTATTGGCTACACGTTTGGAGCCGGGATGTTTGAAATGGGGGAAAT
TAAAGGCGGGAGTCCTTATGGTGCCGGGACATATGCAGGGGATGGGTCGAGACAACCCTCAGGAATCGAACTCGAGCAAGCTTTTC
ACCAGGGAAAACACATTGCAGCCATCACTAAGAAGCTCAAACAAACTACTTAATGTGGGTCACGAATAGTTTCCCGATTCATCAAA

TAAATTTGTATGCTTTTTTTTTTTTTGTCGGAATAAATTGATCTTAATTGGTGTACTTCGTTTAGATATCGATTACGAATTGCTTT
AAGTTCATTAGATATACAATTTGCTACAGAAAATTTTACTCAGTTGTAGCTATGAATTGATGTTATATTGTCTATGTTTATTTTTC
TTTGCGTATTGAAATTATGGCCTTTAATCGTGTCTAAAATTATATTTTGATTTGAAGAAAAAAATAAAAGTAGTCAAACTCGGGCA
ATTGCATATTCAAGCTTGTGTAATTTCGTCTCTATAGCTTTCTTGCAGCAAATTTTTT
>TvQR2 Triphysaria versicolor Tv.c6809_g1_i1.4592
CTTCCCAAGAACCTCATCCACCAAGAAAATCAAACTAAACCAAAATACAGAGCTAATTAAACAGTAACCCTTTAATATTCCAATGG
CAACCAAAGTCTACATCGTTTACTATTCGACATACGGACATGTTGAGCGGCTAGCGCAAGAGATCAAGAAAGGAGCTGAGTCTGTC
GGAAATGTTGAGGTTAAACTATGGCAGGTTCCTGAAATTCTGTCGGATGAGGTTCTTGGGAAGATGTGGGCCCCACCAAAGAGCGA
TGTGCCGGTGATCACACCCGATGAGCTCGTCGAGGCTGATGGTATCATATTTGGTTTCCCAACGAGGTTTGGAATGATGGCCGCTC
AATTTAAAGCCTTCTTCGATTCAACCGGAGGTTTATGGAAGACTCAAGCACTTGCCGGCAAGCCGGCTGGCATCTTTTTCAGCACT
GGAACTCAAGGCGGCGGTCAAGAAACTACCGCATTAACAGCCATTACACAGTTGACACATCATGGTATGATCTATGTCCCCATTGG
CTATACATTTGGAGCTGACATGTTTAATATGGAGAAGATCAAGGGTGGTAGCCCATATGGCGCTGGCACGTTCGCCGGAGCAGATG
GATCTAGACAGCCTTCGGACATCGAACTCAAACAAGCTTTTCACCAGGGAATGTACATCGCCGGCATCACAAAGAAGATCAAGCAA
ACATCGGCCTAATTTCGACCATAAATTTCATTCTACGAATAATAAACATATGTTGTTTATTATATACTTGATTCCTGTTTTATTTT
ATGATTTCCAGGCTTGATGTCTTCAACAAATAAATGTTACACTTTTTATTCGGAATAAAATTGATGTTCACATAAAAAAAAAAAAA
AAAAAAAACAATTACCACTAACTCCGAAAAATAAATAAAATAGTAACGACAAACAGTTGACCCCAGCGGCCCAGCCCATCAAAATT
GATCACAAAATCTCATTCTCATGAAGAATGATAATCAAGCCCAAGCTCCATTTTCCATAAGCCGTCTTCGCAAACAGGCTCATCAC
TATGGCTCGGCTTAA

>PaNDC1 Phelipanche aegyptiaca Pa.c24070_g1_i2.305
GTTTAAGTTGACTTGTGAGAGCTCGTCCTTTGTGAGACACTCGTGGAGCCAAGACGGCCACACACCCATACCGCCCAGCTCCGCCA
CCTTCGTCCCATTCTACCGTCGATCATAGATGGCACACGCAGTTGCATCTCTCACACCATCATCACTGCTCAGCTGTACGCTTTCG
TTTACCCCAGTATTTGGCGAGAAATCCCAGTTGCGGAATTTATTCCAGCAGCATTCGCTTAAATTGTGGACCAATTCGGGGTTCTA
TTCCAGAAGGCGTGGAGTCCCTGTTGTTGGTTCGAGCTCGAGCAGAGGGTACGGCGTCGTTTCTGCTGTTTCTGAAAGTGAGTCTC
AGCGTCCCAGTTACGTTTGGCCTGATAACGAGAAGAGACCAAGAGTGTGCATACTTGGTGGTGGTTTCGGAGGATTGTCCACTGCA
CTGAGATTGGAATCACTTGATTGGCCGGATAGAAAGAAACCACAGGTGGTTCTTGTTGATCAATCCGAGCACTTTGTCTTCAAGCC
ACTATTATATGAACTTCTCTCTGGAGAAGTAGATGAATGGGAAATAGCTCCTCGTTTTTCAGACTTGCTGTCCAGCACTGCTGTGA
AGTTTTTGAAAGACAGGGTTAAATCTTTACATCCCTCTTATCATTACGGACTGGTTGGGGCTTCCATAACTCATTCTGCTGGAGTC
GTGCATCTTGAAAGTGGTCTCCGTATTGAATATGACTGGTTGGTACTTGCCCTCGGAGCTGAAGCCGAACTTGATGTTGCGCCAGG
AGCAATAGAATATGCATTACCATTTTCCACTCTTGAGGATGCTCGTAGAGTCGATGAGAAGCTAAAAACACTTGAGCGGCAATTCT
TTGGTAAAAACTCTCCAATTCGTGTTGCCGTTGTAGGATGTGGTTACTCTGGAGTTGAATTGGCTGCCACAATATCAGAAAGACTA
CAAGCTAGGGGAGTTGTACAAGCAATCAATGTGGGCAAAACAATCTTGTCAAATGCTCCACCTGGCAACAGAGAATCTGCAATGAA
AGTTCTCTCATCCAGGAATGTTCAGCTTTTATTGGGTTACTCTGTTCGCTGCATAAGAAGAAATGTCGAGTATGCGGTTTCATCGG
AACCTACCAATGTAGAAGCTGATCATGATAAAGTGGAAGCTCATAATCAGAAAAGACTTGTATTGGAGTTGCAGCCTGCTGAAAGG
GGCATGCAGAATGAAGTTGTTGAAGCAGATTTAGTTTTATGGACTGTTGGGTCTAAACCTGTACTTCCTGATCTTGAACCCAGTGA
TGAACCCATTAAACTTCCTTTAAATGGTAGGGGGCAAGTAGAAACCGATGAAACTCTTCGCGTTAAGGGTCACCCACGTATATTTG
CAGTTGGGGACTCTTCTGCTGTGAGGGATAGACAAGGTAAATTGCTTCCAGGCACCGCACAGGTAGCATTGCAACAGGCAGATTCC
GCAGGCTGGAATTTATGGGCCGCAATTAATGGCCGACCGCTATTACCATTTAGGTTTCAGAATTTAGGTGAGATGATGACTCTTGG
GAGATATGATGCTGCTATTACACCAAGTTTCATCAAGGGTCTGACATTGGAGGGTCGAGCTGGTCACACTGCAAGGAAAATAGCCT
ACTTAATCCGACTACCAACAGAAGAGCACCGGTTAAAAGTAGGGATCAGTTGGTTAACGAAGACGGCTTTAGAGTCTACTGCGTTA
CTGCAGAATACTGTTACCAGAATGCTTTCGGGGAAGTAGAAGACGTTAGTATATCATGTGATTAAATTACTACCAAATTATCTTAC
ATGACAATATTATTTTTGAACTATCTTACATGCAATATTTTAAGAACAATAAGAAAATACCTGAAGCCTTCTGGCAGGTCTTTGAA
TTACATTGGATTGAGGTCTGTGTTTCTTACCATGTTATCTAATAGACCATCAATGCGAGA
>ShNDC1 Striga hermonthica Sh.c14874_g1_i1.8637
CGCTCACCTGCTCGCTCTCACTTGTGGAGCCACGACAGCGTAGTAATAGTATACCCCCACTTCCCGCCGGCAATTCCACACTCTTA
CACCCTCCACCGTAGATGGCGCTCGCTGTGTCTTCTATAACATCACTAACATTAATCCAGAGTGAGCGGTTCCGGCCGGTGAATCC
ATTCCGTGTTCTTGGTTCGAGTAGGTTATGGACCTACTCGATGTTCTTCTCAACCTCCGCCAAAAGTCCTCCACTCCGTGTTCTTG
GTTCGAGCTCCGGTGGAGGTTTTGGCGTCGTTTCTGATGCAGAATTTCAGCCTTCCAGTTACGTTTGGCCTGATAACAGGAAGAGA
CCAAGAGTGTGCATACTAGGTGGTGGTTTCGGAGGGCTGTACACTGCATTGAGATTGGAATCACTTGATTGGCCAGATGGAAAAAA
ACCACAGGTGGTTCTTGTTGATCGATCTGAGCATTTTGTTTTCAAGCCGCTACTGTATGAACTCTTATGTGGAGAAGTAGATGAAT
GGGAGATAGCCCCTCGTTTCTCAGACTTGCTGTCAAGCACTGGTGTGCAGTTTTTGAAGGACAGCGTGCAATATTTACTTCCCTTT
GATCAATATGGTATGGATGGGGATGCGTTAACTCATTCCGCTGGAGTAGTGCATCTTGAAAGTGGTCTCCTTATTGAATATGATTG
GTTGGTACTTGCTCTTGGAGCTGAAGCTAAACTTGATGTTGTGCCGGGTGCAACAGAATATGCATTGCCGTTTACCACTCTTGAGG
ATGCTCGTAGGGTCAATGAGAAGCTAAGAACACTTGAGCGTGAATTCTTTTGTAAGGACTCTCCAATTAGAGTTGCAGTCGTTGGA
TGTGGTTACTCTGGAATTGAATTGGCTGCCACAATATCAGAAAGACTCCAAACACGGGGAGTTGTACAAGCAGTCAATGTTGAGAA
AACCATTTTGTCGAATGCTACTCCTGGCAATCGGGAATCTGCATTAAAAGTTCTCAAGTCCAGGAGTGTTCAGCTTTTGCTGGGTT
ATTTTGTCTGTTCTATAAAAAGAGTTGCACAGTGTGACAATAAAGTAGGAGCACAAAACCATGAAAGACTTAAATTGGAGTTGCAG
CCTGCTGAAAGGGGTGTGCCTAGCCAAGATGTAGAAGCAGATTTAGTTTTATGGACTGTCGGTTCAAAACCTGTGCTTCCTCAGCA
CAAACCCGGTGATAAACCCATATCACTTCCTCTAAACAGTAGGGGTCAAGCTGATACTGATGAAACTCTTCGTGTTGAGGGTCACC
CGCGTATATTTGCTATTGGCGACTCTTCTGCTATGAAAGATGGTCGAGGAAATTTGCTTCCAAGCACTGCCCAGGTAGCATTTCAA
CAGGCTGATTTTGCCGGCTGGAATCTGTGGGCCGCAATAAATGGTCGACCTCTGTTACCATTTAGGTTTCAGAATTTAGGTGAGAT
GATGACGCTTGGGAGGTATGATGCTTCTGTTTCACCAAGCTTCACGGAGGGCCTGACATTGGAGGGTCCAATTGGTCACACTGCGA
GAAAAATAGCATACTTAAGCCGACTGCCAACAGACGAGCACCGATTAAAAGTAGGGATCAGTTGGCTAACAAAGGCTGCCATTGAA

TCAACTGCATTACTGCTGGATAGTATTACCAAAATGCCTTTCGAATTGTAGAGGACGGTGTAGTATCATATGCTGTGCTGCTCTCT
CTCTCTCTCTCTCTCT
>TvNDC1 Triphysaria versicolor Tv.c5468_g1_i1.261
GAAGCAAGGGCCATATATTTTTATCAGCCAACTGTTAGTAAAGCTGGTTCTTTCTCGACCACTCGTGGAGCCAAGACGGCCGCGTA
TACCATTTCGCCCCCGTCTCGTACCGTCGATCATAGATGGCGCTCGCAGTTTCCTCTCTCACGTCATTAACACTACGCAGCCGAGA
GCAATTCCAGTGCCATTCGCGTAAGTTTTGGACCAGATCGGGGTTCTTTTCTAGCTCTTCCTCTAGAAGAAGAGGAGCCTGTGTTG
TTGGTTCGAGCTCGAGCAGAGGATACGGCGTCGTTTCTGCTATTTCTGAAAGTGAATCTCAGCGTCCGAGCTACGTTTGGCCTGAC
ATAAAGAAGAGACCAAGAGTGTGCATACTAGGTGGTGGTTTCGGAGGGTTGTACACTGCACTGAGATTGGAATCACTTGATTGGCC
AGATGGAAAGAAACCACAGGTGGTTCTTGTTGACCAATCTGAGAACTTTGTCTTCAAACCACTACTATATGAACTCCTCTCTGGAG
AAGTAGATGAATGGGAAATAGCTCCTCGTTTCTCAGACTTGCTGTCAAGCACCAATGTGCAGTTCTTGAAAGACAGGGTGCGATCT
TTACATCCGTCTGATCATTATGGAATGGATGGGGCTGCTGTAACTAATTCTGCCGGGGTCGTGCATCTCGAAAGTGGTCTCCTTAT
CGAATATGATTGGTTGGTACTTGCCCTTGGAGCTGAAGCCAAACTAGATGTTGTGCCAGGTGCTATAGAATATGCATTGCCTTTTT
CCACCCTTGAGGATGCTCTTAGAGTCAACGAGAAGCTAAAAGCACTCGAGCGGGATTTCTTCGGTAAGGACTCTCCAATTCGTGTT
GCCATTGTAGGATGTGGTTACTCCGGAGTTGAATTGGCTGCCACAGTATCAGAAAGGCTGCAAGATCAGGGAGTTGTACAAGCAAT
CAATGTTGATAAAACAATTTTGTCAAATGCTCCACCTGGCAACAGGGAATCTGCAGTAAAAGTTCTCTCGTCTAGGAATGTTCAGC
TTTTGTTGGGTTACATTGTCCGCTGTATAAGAAAAAATGTTGAAGATCACAATCACGAAAGACTTGTACTGGAGTTGCAGCCTACC
GAAAGGGGAATACAAAGCCAAGAAGTCGAAGCGGATTTAGTTTTATGGACCGTTGGGTCTAAACCTCTAATTCCTGAGCTAGGATC
GAGTGATAAACCTGTTTATCTTCCTTTAAACGGTAGGGGACAAGCTGAAACTGATGAAACTCTTCGAGTTAAAGGCCACCCACGAA
TATTTGCAGTTGGTGACTCGTGTGCTGTGAGGGACGGAAAGGGTAAATTGCTTCCAGGCACCGCTCAGGTAGCATTTCAGCAGGCA
GATTTTGCAGGCTGGAATCTGTGGGCTGCAATTAATGGCCGGCCTCTATTGCCATTTAGGTTTCAGAATTTAGGCGAGATGATGAC
TCTTGGGAGATATGACGCTGCTATTTCACCGAGTTTCATCGAGGGCCTCACATTGGATGGCCCAATTGGTCACACTGCCAGGAAAA
TAGCCTATTTAATCCGACTACCAACAGACGAACACCGGCTAAAAGTAGGGATAAGTTGGCTAGCCAAGACTGCTTTAGAGTCTACT
GCATTACTGCAGGATACGGTTACCAGGGTGCTTTCCGGACAGTAGAAAGACGTATATTAATTATCATGGATTCTAGATACAAATTG
GCGCAGAGCATTGTAAAAGACAATTACAGTTTACAATGTATGCATAAATGTAATTTTGATGAAATAACTATATATGGGGTGCCCAA
TATCTGAATATTTAACATAATTATATTTCAAGAAAAGATTCTTTTTTAACATGTGGTAAAATTATATTATTGATTGTTATGTAGAT
TTTATGACATTGTGTTGCATATATTTC

>PaPRX1 Phelipanche aegyptiaca Pa.c19539_g1_i2.433
ACACAAATTATTTCAATATGGACAACACAAAATCCATCAACTCTCTTCTCACTCTAACAATCTCATCACTCATCCTCCTCTCGCTC
TCCATCACACCATCTCATGCACAACTATCTCCCACGTTCTACGACAGTACATGTCCCAATGCACTCAGTACCATCCGCACTTCCAT
CCGACGAGCAGTAGCTGCCGAGCGCCGCATGGCTGCTTCGCTTATCCGACTCCATTTCCACGATTGCTTCGTACAGGGTTGTGATG
CTTCCATCTTACTCGATGACAGCCCAACGATCCAAAGCGAAAAATCCGCGGGCCCAAATGTAGATTCTGTAAGAGGGTATGATGTG
ATAGAGACCGCCAAACGTGAGGTTGAGAGTATATGCCCTCGCAACGTTTCTTGCGCTGATATTGTAGCGTTAGCAGCACGTGACGC
CTCTGTCGCCGTCGGCGGTCCGACATGGACTGTAAAGCTTGGGAGAAGGGACTCTACCACCGCTAACCCTAACGAAGCCAACACAG
ATCTTCCTAGTCCGTTCGCTAGTCTCCAAACTCTCATCACTGCCTTCGACGATAAGGGACTTAGTGAAACAGACATGGTCGCCCTA
TCAGGATCACACACAATAGGTCAATCTCGATGCTTCTTGTTCCGCTCTAGAATATACAGCAACGGAACCGACATCGATCCCAACTT
TGCCAGCACCCGGAGGCGCCAATGCCCTCAGACAGGCGGTGACAATAATTTGGCGCCGCTAGATTTGGTGACGCCAAATTCATTCG
ACAATAATTATTTTAGGAATTTGATCCAACGGAAGGGACTTCTCGAATCTGATCAAGTCCTTTTCAATGGAGGATCCACAAACGCT
CTCGTCACTAGTTATAGTAACAACCCACGACTCTTTGCGACAGATTTCGCAAGTGCAATGGTCCGAATGTCGGAGATTCAACCCCT
CCTCGGATCAAATGGGATCATAAGGAGGGTTTGCAATGTTATCAACTAGTTATAAAATGTTTTAATTACTGCATGCATGCCACTAA
GCAATCAAATCAATGTTGGGGTTAATAAAGCGAGTGAAATCTGCTATTCTGTAATTGTGTATATCTTGTTTATCGCATTTTCGTAA
AATGTTCTTTGTCGTTATTTGCTCATGGAATAATATTTATTGAGATTTGTTATATTTTTCGATTAACAATTGTACTTGTGTATTAG
TGTTCATCTTGATTAACCCAATAATAGATCATATGTGCCATGGACTTATTC
>PaPRX2 Phelipanche aegyptiaca Pa.c24858_g1_i1.13544
TGATGAAAAAGCAGAAAATTGTTAGTCTTTCACTAAACTTCAGCGCCAAGCAATTAAGTTTAGTGAAAGACAACAATTTTCTGCTT
TTTCATCAATATGGAGAACAAAAAATCCATCAATTCTGTTGCTGTGACGCTAGCAATATTATCACTCATACTCTTGCTGTCCAGCA
CACCAACTCAAGCACAACTATCTCCCACATTCTACTCTCGCACATGTCGTAATGCGCCAACTATAATTCGCAATTCCATCCGCAGA
GCAATATCACGTGAGAGGCGAATGGCGGCCTCACTCATTCGTCTCCATTTTCATGATTGCTTCGTTCAGGGATGCGATGCATCAAT
CTTACTAGACGAAACACCTTCTATTCAAAGTGAAAAGACTGCATTCCCAAATGTCAATTCCGCAAGAGGGTTTGATGTGATAGAGG
CTGTCAAGCGTGAGGTTGAACGTATATGCCCTCGAGTTGTTTCTTGTGCCGATATTCTAACTTTAGCAGCTCGCGATGCCTCCGTT
GCTGTCGGGGGTCCATCATGGACTGTGAGGCTCGGGAGAAGGGACTCTACCACTGCTAATCGCGGTCAAGCGAACACTGATCTTCC
TAGTCCGTTTGCTGGTCTCCAAGCTCTCATCACTGCCTTTGCTGATAAAGGACTTAGTGAAACAGACATGGTTGCCCTATCAGGAT
CACACACGTTAGGTCAAGCCCAATGTTTCTTATTCCGTGCTAGAATATACAGCAATGGAACTGATATTGACCCCAACTTCGCCCGC
AATCGAAGACGCCAATGCCCTCAGACCAGTGGCAACAGTAATTTGGCGCCACTCGATTTGGTGACCCCAAATTCATTCGACAACAA
TTATTTTAGGAATTTGATCCAAAGGAGGGGACTGCTTCAGTCAGATCAGATCCTTTTTAGTGGAGGATCCACAAACACTCTCGTCA
CTCGTTATAGTAACAACCCACGACTTTTTGCTTCAGATTTTGCAAGTGCAATGGTCCGAATGTCAGAGATCCAACCCCTCCTCGGA
CCGAATGGGATCATAAGGAGAGTTTGTAATGCCACCAACTAGTTTGTTACTTAAAAACTTGTCACAACCACTATTTAGCAATCATA
TTATTTTTGGGATTATTATATAGCAAATAAATCAGCTGTTCAACAATTGTATGTACTTCTGGTACTATTCATGTTATGTCCAATGT
TGTGAAGCCATTATTTTCTGATAATGTAATGTGTCTTTTATTCAAATGTTTTACCATATTGAGTACAAAAAAAAAAAAAAAAAAAA
AC
>ShPRX1 Striga hermonthica Sh.c9817_g1_i1.12343
CAATTTCACCAGCTCCGGAATCAACTTCACCTCAAACCAAATTGTCCAAGTACGTACTTAGTCTTAGCATGGCCACCGAAAAGCCC
ATCAACTTTGTATTCACCCTAACCACACTCACAGTACTACTCATCCTCTCCCTGTCGATAATCCCATCCCGAGCCCAGCTGTCCCC

GACTTTCTATGCCCGCACATGTCCCAATGTGCTGCGTACAATCCGTGGTTCCATCTGGAGGGCCGTGGCTCGTGAGCGCCGCATGG
CGGCCTCACTCATCCGCCTCCACTTTCACGACTGCTTCGTGCAGGGCTGTGATGGTTCCGTGTTGCTGGATGATGGGCCCACGATC
CAGAGCGAGAAGTCGGCTGTCCCTAACCTCAACTCCGCAAGAGGGTTTGATGTGATTGAGGCCGCCAAGAGAGATGTCGAGCGAAC
GTGCCCAGGTGTCGTCTCGTGTGCCGACATTCTGGCTGTGGCCGCGCGTGATGCCTCTGTGGCAGTGGGTGGGCCGTCATGGAATG
TGAAGCTCGGGAGAAGGGACTCTACCACAGCTAACCGTGATGCGGCGAACCGTGAACTACCGAGCCCATTTTCCAGCCTTCAAGGC
CTAATCACTTCTTTTTCCGACAAGGGCCTCAGTGAAAGAGATATGGTCGCCCTCTCTGGATCACACACGATCGGGCAGGCCCAGTG
CTTCCTGTTCCGCAGCAGGATCTACAGCAACGGCACGGACGTCGATCCACTCTTTGCCCGTCTGAGGAGTCGGACGTGCCCTCAGA
CGGGTGGGGACAGCAACCTTTCCCCGCTCGACTTCGTAACTCCCAACAGGCTCGACAACAACTACTTTACGAACCTTCGTCAAAGG
AGGGGTCTCCTTCAGTCGGACCAAGTCCTTTTTAGCGGGGGCTCCACAGACAGTCTCGTCTTCAACTACAGCAGAAATCCCCACCT
GTTCGCCTCCGATTTCGCTAATGCCATGCTCAAGATGTCCGAGATTCAGCCCTTGCTCGGGTCCAATGGGATCATAAGGAGGGTTT
GTAGCGTCCCAAACTAAGTGTGCGGTTAAGTAATGTTTGCAAACACTTCAAAAAGAGAAATAACTAAGAGAGATACTTTTCCTAGT
GTTTGCGAATACTCTGTTTGCGGCTGCGAGGGTTATATATGAAAAGAGGTTATTTTACGTAATTGTATTGATTAATGGAGCATATG
AATATGCTACTCTATGATCTGTAACCTTTAATTGTAAAATGGCTGATCCGTCTTAGTAGTTTGCCATCAAATACTTGTGGTGTATT
CCTTTAATAAATATTTTGAAGGCTTCAGAAAAAAAAAAAAAAAAAAAAAA
>ShPRX2 Striga hermonthica Sh.c9802_g1_i1.12342
AACAATCAAAACAAGCACCAATTTTCCTTAGTAAATATTACAATAACAATGTCAGCAACGCGTTTTACCGATCAGCAGTGACCAGA
TTAAATTCAAACACACACACACACGTTCATCCCTGCTCCCTATAAATTCAGCACCCAACTTCTCAATCACCTTCATCTCCAAGTAA
ATTGACTTTAGCACTTGTACGTGTCTTTCGATTTCAATTAGTTTCTATATCTCTTTTTCGCAACATGGGCACGACGAAAGCCGTCA
ACTTTGTCCTAACCTTAACAACAATCTCACTACTCATCTCCACCACCCCGACCCAAGCCCAGTTGTCCCCGACTTTCTATAACCGT
ACATGTCCCAATGCACTGAGTACAATCCGTACATCCATCCAGCAGGCGATTTCCCGTGAGCGCCGCATGGCTGCCTCCCTCATCCG
CCTACATTTTCACGACTGCTTTGTACAAGGCTGCGACGCTTCCATCCTGCTGGACGACTCGCCCACAATACGAAGCGAGAAATCGG
CCGGCCCAAACGCCAATTCCGCCAGAGGGTTTGATGTGATTGAGGCTGCCAAGAGAGCAGTCGAGGGTATTTGCCCGGGCGTCGTC
TCGTGCGCCGACGTTTTAGCCGTGGCTGCACGTGATGCCTCAGTTGCTGTTGGAGGGCCGTCATGGAATGTAAGAGTTGGGAGACG
GGATTCTACGACTGCCAACTTCAATGCAGCAAACACAGATTTGCCGGGACCTTCTTCTAGTGCACAAGGCCTCATTACTGCTTTCC
GCAATAAGGGACTCAGTCAGACAGACATGGTTGCCCTCTCAGGGGCACACACGCTGGGGCTAGCCCAGTGCCAATTTTTCCGGGCC
AGAATCTACAGCAACGGCACGGACATCGACCCCAACTTTGCAACCACAAGAAGAAGCACATGCCCACAGTCGGGCGGGAACAGCAT
CCTTGCCCCGCTCGACATCCAAACCCCCAACTCGTTCGACAACATCTACTACAGAAACTTGGTCCAGCGGCGGGGGCTCCTCCAGT
CGGATCAGGTACTTTTCACCAGTGGGTTAACCAATGCCCTAGTCAACACCTACAACAACGACAGTCGGCGCTTCGCCTCCGACTTC
GCGAGCGCCATGCTCAGGATGTCTGAGATTTCGCCGTTGGTGGGGTCCCGTGGGGTCATAAGGAGGGTTTGCAATGCCCTCAGCTA
GACGATGTGGCTGTGGTGCTGTCAGGGTTATATAATGAATTATTAAAACCACTGAAATGCAGTGGCTTTATTATGTGATTTAATTT
GTTTCTTGTCCGCTTAATTTTTCATTTGTTGTCAGTCTTGGTTTTTTTTTTTTTTTTTTTTTCCTTTTTTTGTGAGTGCCAGTAACT
CCGATAACGCACCTAGCATACCGGTCCCGTCAATCGTCCAGCAGCCGGCGATCTTGGTTTAGCAACGACTCTGTCCGACCAAGCCA
AGGTCACCTCGCCATCGCCGTTAACATCACGTGGGTCGGCAGTTTCAAGCCATCGAGACCTGTTTGAAATTTCAGACCCCGTCACC
CTGTCTCGTCCATCCTCGAGCAAAACCGCCGACGAACCAGTCGAGAAATTCTCGTCGGCGACCTTCAACAGCAGTGGCCCTCCTTC
TGTCCGTTAACGTCGGTCAGTGAGCAGCGACGGTTTTCCTCCGAGGAGTTTCGCCACCAAAGTCGGAAAATCAGCATAGCCGCTCA
CCAGGAGTCCTGTTCGCAACGAGCAGTTTCTCAGCCGTCGCCGGCGATCCACCACCACCGCGCGACTCCTTCTGGCGAGCAGCAGG
TCCCGTCGGTTCGCTGTCGCGTCTGATCTCTTGTCACTTTCGTAGGCAAGCGAACAGGTACCGT
>TvPRX1 Triphysaria versicolor Tv.c5216_g2_i1.11956
TAACTTTTTATTCTTTCTTGTCAACGCTTAATTTCAAAATGAAGATCAAAAGCTTAGACTCCATAAACATTGCCACTATATTCACA
ATCACAACTCTCTTATTCTTGCACAATTTCATGCCATGTGAAGCACAATTATCTTCAACGTTCTACGACAGAACGTGTCCAAACGC
ATCTACTATCATCCGTAATTCCATCCGTCGAGCAATAGCTCGTGAGAGGCGAATGGCTGCATCCCTCATTCGCCTCCATTTTCATG
ATTGTTTCGTTCAGGGTTGTGATGCATCGATCTTACTCGACGAGACACCTTCCATCCAGAGCGAGAAGACTGCTTTCCCGAATGCC
AATTCTGTAAGAGGCTATGATGTCATCGAGGCGGCAAAGCGCGATGTTGAACGTGTTTGTCCTGGCGTCGTCTCTTGTTCCGATAT
TCTAACATTAGCCGCACGTGATGCCTCCGCTTACGTTGGTGGTCCATCGTATACCGTTCGTCTCGGGAGAAGGGACTCAACCACTG
CGAGTAGAGCCCAAGCAAACGTAGAACTCCCAAGCCCGTTTGCCGGTCTTGACGGCCTCATTTCCGCTTTCGACAACAAAGGACTT
AATGCAAGAGACATGGTTGCCTTATCTGGATCACACACATTAGGGCAAGCTCAATGTTTCTTGTTCCGCTCAAGAATTAGCACCAA
CCGAACCGATATTAACGCCGGATTCGCTAGCACTAGAAGACGACAATGCCCTCAAGTCGGGGGCGATGCAAACTTGGCCCCACTTG
ATTTGGTGACCCCAAACTCGTTCGACAACAATTATTTCAGGAACTTGGTCCAGAGGAGGGGTCTTCTTCAGTCTGACCAAGACCTC
TTCAACGGAGGGTCAACTGACAACATCGTCTCCGAGTATGTTAGGAACCCTCGACTTTTCGCTTCTGATTTTGCGAGTGCTATGAT
CAAGATGGGCGAAATTCCAGCAACACAACAAAATGGGATCGTAAGAAGGGTTTGCAGTGCCATCAACTAAGTGATCGATATATCTT
TATTCAAAACTAATTATTGATGGTTACATTGCTGATTTCATTTACTTTTTTGTCTTCATTGTTTATTAATTGTATTTGTAATAAAA
ATAACTTGTGATTTTTGTACTACTATTCTTTGATTTATGGTATTAAATTATGTAACTCTCATTTTTAAATTAAT
>TvPRX2 Triphysaria versicolor Tv.c5326_g1_i1.12010
ACTTGTAAAATACATCTTCAATTCAAATTCATCTCATTCAATTTTCCTTTTGCTCTATAGTTTTGTGATCTTTCAAACTCCAAATT
ATTTCAATATGAAGACCATAAACTTTCTTGCAACCCTAACAATATTATCACTAGCCCTCTCCATTATTATCACACCAACTCAAGCA
CAACTATCTCCCACGTACTATACCGCCACGTGTCCGACTCTCGTAGCCGTAATTCGTACATCAATCCGTCAAGCGATTGCCCGTGA
GCGGCGTATGGCCGCATCGATTATCCGTCTTCATTTCCACGATTGTTTCGTGCAGGGATGTGATGCCTCGATCCTTCTAGACAACG
CGGCGAATATCCAGAGCGAGAAGTCCGCGGGGCCCAATGTTAACTCGGTGAGGGGGTTCGATGTGATCGAAGCCGCCAAGCGTGCG
GTCGAGGCCGCATGCCCCGGTGTCGTTTCTTGCGCGGATGTTCTAACTATAGCGGCTCGCGAAGCGTCCATTGCTGTTGGTGGGCC
AACATGGAATCTGAATCTTGGAAGACGGGACTCGACCACGGCTAATCCTAATGCAGCCAACTCGGATCTTCCTTCACCCTTTTCTA
CTCTCCAAGGTCTCATCGACGCCTTCTCTAATAAGGGACTTAGTGTCACCGACATGGTTGCCCTATCAGGGGCTCACACATTAGGC
CAAGCCCAATGCTTCTTGTTCCGCGGTAGAATAAACAGCAACGGGACGGACATCGACCCTAACTTTGCTAGCACACGGAGACGCAC
GTGCCCATCGACCGGTGGGGACAGTAATTTGGCCCCGCTCGATTTGGTGACCCCAAATTCGTTCGACAACAACTACTATAGGAACT

```
TGGTTGCACGTAGGGGTCTTCTTTCATCTGACCAAGTCCTTTTCACGGGGTCCACGGCCGCCACGGTTACACGTTATAGTACTCGA
CCCAGAGATTTTGCAACGGATTTTGCAAGTGCGATGGTTAGAATGTCGGAGATTTCGCCCACTCTCCGACCTAATGGGGTCGTACG
AAGGGTTTGTAATGTCATCGGCTAGTTAATTAATTAAATTATGGGAGTCAAATCTGCTAGTCGATGTGTAATTGTAAGATAACTAT
TTATGGTGCATGTGTGTTTTTGTGTAATTTCAATATTGGTTGTTGTTACTTGTTGATCGTTAAATTGTGAAATAAAATTGATGTGA
TTTGTTGTGTTTTGTCAATCAATGGTTTCTTTAATTATTTTCGTGTCTTGTAGTTGTTGTCATTATTGATGTAATTTTATATTTTT
T
>TvPRX3 Triphysaria versicolor Tv.c105997_g3_i1.21359
TTAGCATCATCCTCTTCATCATCAATCAACTCAATTCATCTTTATTATCCACTTCTAAATCCTTGTTTTTTGCTCTACCCTAATCT
TTCAACATGAATTTTCTTGTAACCCTAACAATATTATCATTGGCCCTTGCCACTATTATTACACCAACTCAAGCCCAACTTTCTGC
GACGTACTACGCCACCACGTGTCCTAACTTCGTCACCATTGTCCGAACCTCCATCCAGGCAGCAATTACCCGTGAGAACCGAATGG
CGGCGTCAATCCTCCGCCTTCATTTCCACGATTGTTTCGTGCAAGGATGTGATGCCTCGATTCTTCTCGACGATACAACCACGATA
CGGAGCGAAAAGAATGCGGGCGCCAATGCTAACTCTGCTAGGGGGTTCGATGTGATCGAAGCCGCTAAGCTTGCCGTGGAGCAAGT
ATGTCCCGGGGTCGTTTCTTGCGCCGATGTGCTCACGCTAGCGGCCCGTGAATCCTCTTTTGCTCTTGGTGGGCCCACATGGAATC
TACTATTCGGGAGAAGAGATTCTACAACAGCTAACCTTGCGGTTGCAAACACCGATCTACCTGGACCTTCTTCTACTCTCCAAGGC
CTCATTACCGCCTTCTCGAGGAAGGGGCTTACTGCCACTGACATGGTTGCCCTATCAGGAGCCCACACACTGGGCCAAGCCCAATG
CGCCTTTTTCCGCTCAAGAATATACGGCACCGCGAACATCGACCCGATCTTTGCCAACAATACGAGACGCGCGTGCCCTCAATCGG
GTGGGAATAGTAATCTATCACCGCTCGACGTTCAGACACCCACTAGATTCGACAACTTCTACTACCAAAACTTGGTCGCACGGAGG
GGTCTACTCCAGTCGGACCAAGTCCTTGCTAGCGGGTCCACAAATGCCACCATCGTACGTTATAGTACCAATCCGGCAGCATTTGC
AGCCGATTTTGCAAGAGCGATGATTAAAATGTCGCAAATTGCACCGACGGTTCGACCGAATGGGATCGTGAGAAGGGTTTGTAGCG
CCATCAACTAGTTACGTGTGTGTGATTAATCACATGCATGCTAATTAAGCAATTTCTTGAGATTAATTAATTATATTGTTGGTGTT
AATAAGTTAAGCTAGTAAAATTGTGCTAGCTAGTTGATTTGTAATTGTTTATTCAGTTACGGTTCATGTTTCGTGTAATGTTTTTT
TAATGTTGTTGTTATTCGTTGTTAATCGTGTTGTCGTGGGATGAACTAAATGTGATTTGTCGTGTTTTTGTCGGTCAACGGTGTTT
TTTTAATTCATGCTTGTAATTTTAATTTTTTATTGTTCTAGTCTATTGTCAATTACATAAAAATGTCTCATATAATATTTTTT

>PaQR1 Phelipanche aegyptiaca Pa.c25149_g1_i2.4599
GGGGAACGTACGCATCACTTTTTTATTTATAATCAATTAATTAATGAATAATTGTGCGCCACCCACTTTCGATGCTGTAAATTCCG
GACGCCACTTTCAAGCTTCTCAGACCTTCTATTTCTTGCTACAATTCTCTTATTTTCTTCTTTAATTTTACAAATCGGTCTCCCAC
AGTCGCTGAAAATTACCCAGTATTTCGAGATTTGAGTATGGCGGGGAAGCTTATGCATGCGGTTCAGTACGACGGTTATGGCGGTG
GAGCTGCTGGTTTGAAGCATGTTGAAGTTCCAATTCCTACTCCTAGTAAGGGTGAGGTCCTGCTAAAGCTGGAAGCCGTAAGCTTA
AATCCTATCGACTGGAAGACACAGAAAGGCTTGCTTCGTCCTCTCCTTCCTCGAAAATTCCCTTTTATACCTGCAACCGATGTAGC
TGGAGAAGTAGTGGAGGTTGGAACTGGAGTCGAAACCTTCAAACCTGGTGACAAAGTCGTTGCAATGCTGAGTCATCTTACGGGAG
GCGGTTTGGCTGAATTCGCAGTGGCCAAGGAAAACTTAACCGTCTCTAGGCCTCCTGTGGTCTCAGCGGCCGAAGGTGCAGGCCTT
CCAGTTGCAGGCCTCACGGCCCACATGGCCCTAACCCAGTCAGCGGGGCTCAAGCTCGACGGAAGCGGACCCCAGAAAAACATCCT
CATCACCGCTGCCTCCGGCGGAGTTGGCCAATACGCCGTCCAGTTAGCAAAGCTGGGAAACGCGCACATAACCGCCACTTGTGGTG
CCAGAAATGTTGACTTAGTCAAAAGTCTCGGAGCCGATGAGGTTCTCGACTATAAAACTCCAGAAGGGGCAGCCCTTAAAAGTCCG
TCTGAGAAGAAATACGATGCGGTGGTTCACTGTGCTTCAGCTTTGCCCTGGTCAGTTTTTGAACCGAATTTGAGCGCCAATGGGAA
AGTAATTGATATAACTCCTGGGCCGAGTGCCATGTGGACTTATGCTCTAAAGAAACTTACATTCTCTAAGAAACAGTTGGTGCCAC
TACTTCTTGTTCCGAAAGGCGAGAATTTGAAGTTTCTTGTTGAGTTAGTGAGAGAAGGGAAGCTTAAGACGGTGATCGACTCTAAG
TTTCCTTTAAGCGAAGCTGCGGATGCTTGGGCCAAGAGCATCGATGGACATGCTACTGGGAAGATCATTGTCGAGCCGTAAATCAA
TGAGAAGAATGACCTTAATATTATTATGCTGTGAGAAACTTTCCTGTTTGATATGTTATTTGATCTGGTCACCTTTTACTCGTGTG
AAAATGTTACCTGGATATCAGAATGCATAGGAATATATAGTCGTTGGTTGATATAATAAATGCTTTTGTGTGTAAGGTTAAAGAAA
GTTGACGAAAGCTCTATGGATTAAGGCACACAGATCAGTTAATGGCTTAGTCGAAATTTTATTCATACTTCTGTTATACTTGTATT
ATATATGCATGAATGGATGTATATGAGTTGAGAAATCTCTGAACAAAAAAAAAAAAAA
>ShQR1 Striga hermonthica Sh.c18735_g1_i1.2952
TACATATATGCTCTTGATTCCATATTCGTTGAAATTATAGTATTTCGTGATTAAGGTATGGCGGGGAAGCTTATGCGTGCTGTTCA
GTACGAGGGTTATGGCGGTGGAGCTGCTGGTTTGAAGCACGTTGAAGTTCCAGTTCCTAGACCTAGTAAGGGTGAGGTCCTGCTAA
AGTTGGAAGCTACTAGCTTAAATCCCATTGATTGGAAAACTCAGAAGGGCGTACTTCGCCCTTTTCTTCCTCGAAAGTTTCCGTTT
ATACCTGCTACTGATGTAGCTGGAGAAATAGTGGAAGTTGGAGCTGGAGTCGAAAGTTTCAAACCTGGCGACAAAGTTGTAGCCAT
GCTGAGTCATACTACAGGGGGTGGCCTGTCCGAATACGGAGTAGCCAAGGAAAACCAGACCGTACCAAGGCCCCCAGAAGTCCCCG
CTCCCGACGCCGCAGGTCTCCCGATCGCGGGCCTCACGGCCCACATGGCCCTCACGCAGACCGCGGGGCTCAAACTCGACGGTACT
GGGCCTCGGAAAAACATCCTAGTCACCGCCGCCTCGGGTGGCGTGGGCCACTACGCCGTCCAGCTGGCGAAGCTCGGCAACGCGCA
CGTCACCGCCACATGTGGGGCCCGCAACCTCGACCTCGTCCGGAGCCTCGGGGCCGACGAGGTCCTCGACTACAAGACTCCGGAAG
GTTCCGCCCTCCGGAGCCCTTCAGGTAAAAAGTACGACTTTGTGGTCCACTGCGCCTCGGCTTTCCCTTGGTCAGTTTTCGAGCCG
AATCTGAGCGAAAACGGGAAAGTGATCGATATCACGCCGGGGCCTGCCGCCATGTGGACTTTTGCGATGAAGAAAATCACTTTTTC
AAAAAAGCAGCTCGTCCCGTTGCTTCTTTTCCCGAAAGGCGAGGATTTGAAGCGACTGGTCGAGTTGGTGAAAGAAGGGAAGGTTA
AGACGGTGATCGACTCGAAGTTCCCTTTGAGTAAGGCCGAGGATGCGTGGGCTAAGAGCATCGACGGGCATGCCACCGGGAAGATA
ATCGTCGAGCCATAGATTACAGGTAAAATGTTGTTATTACATACATGAGGTAAGAAAACTTGTGGGTTTTTTAAAGAATGTTTTTT
ACATGTTGTGGCTGAACTCTCGTCTGGTTTTAAGAATGTATGGGGACATGTATATGACTTGGTTTGATGAATGAAAAACATGTGAA
TGATATAGTGAATAAAGTTGGCAATTGGGACTTTTTGGATCTCATTATTGTTATAGTGGATCATGGTTGAGTTGTGAACTTTATAC
ATGTACCTATGGTTAGTTTGAG
>TvQR1 Triphysaria versicolor Tv.c107009_g2_i1.16950
AAAAAATCGCTGAAATTTAATTTAATTATGGCCGGAAAGCTTATGCGTGCGGTTCAGTACGACGGTTATAGCGGTGGAGCTGCTGG
TTTGAAGCATGATGAAGTTCCAATACCTAGTCCTGGCAAGGGCGAGGTCCTTATAAAGCTTGAAGCCATAAGCTTAAATCAACTTG
```

ATTGGAAGCTTCAGAATGGCATGGTTCGTCCTTTTCTTCCTCGGAAATTCCCTTTTATACCTGCTACCGACGTGGCTGGGGAGGTG
GTCCGGATCGGACCGGATGTCAAAAACTTTAAACCCGGTGACAAAGTTGTTGCTATGCTTGGCAGTTTTGGAGGAGGTGGCTTAGC
CGAATACGGCGTAGCAAGTGAAAAGCTAACAGTCCATAGGCCGCCCGAGGTATCAGCTGCCGAGAGCTCAGGCCTTCCCATTGCCG
GCCTTACAGCCCACATGGCCCTAACCCAACACATTGGCCTAAACCTCGACAAAAGTGGTCCCCACAAAAACATCCTCATCACAGCC
GCCTCCGGTGGTGTTGGCCAATACGCCGTTCAGCTCGCAAAGCTAGGAAACACACATGTAACCGCCACATGTGGGTCCCGAAACTT
TGACTTGGTCAAAAGCCTCGGAGCCGACGAGGTTATTGACTATAAAACCCCCGAAGGGGCAGCCCTTAAGAGCCCGTCGGGCAAAA
AGTATGATGCGGTTATTCATTGTGCATCGCCTTTGCCATGGTCCGTTTTTAAACCGAACTTGAGCAAACATGGGAAAGTGATCGAT
ATAACTCCCGGTCCGAGGGTTATGTTGACTTCGGCTATGACAAAACTTACGTGCTCGAAGAAACGATTGGTGACGTTACTTGTTGT
GATCAAGGGCGAGCATTTGAGTTATCTTGTTGAGTTAATGAGAGAAGGGAAACTTAAGACGGTTATCGACTCTAAGTTTCCGTTAA
GTAAGGCTGAGGAGGCTTGGGCTAAGAGCATCGACGGCCATGCTACCGGGAAGATCGTTGTCGAGCCATAAGTTAGTAAGATTTTG
TTTTGTTTTATGATATTGTAATGTGGAATTTGGCTTATGACTTGTTTTGGTGATCTTTATGTTTTGATATGTACTCTTTTGTTAAC
CTACTTGTGGTTAGAGTGGCAATTTGTGTCACCATGGTTGTGTTTGTTCGTGTCCTTAAGTCCTATAATGTAATTTTCATATTTTA
TACTTTATTTAGTC

>PaNQR1 Phelipanche aegyptiaca Pa.c18296_g1_i1.11052
AGAGGCATACACGACACTCATTCAAACATCAAAAAGCAAAAAACCAACTACATAATTCATGATGACAGGCGACGACCGCCGCCTCA
CTTCACTTACCCAGCTTTGACCTTTTACGGCAGAGAAGGAAGAATCCATGGCGGCGGTCTCAGCACCTTCTCCAGTCATCAAAGTT
GCTGCCCTCTGCGGTTCCCTTCGCAAAGGTTCTTACCATCGCGGCCTCCTCCATACGGCGATGGATCTATCAAAGTCAATTAAAGG
TTTGGAGATAGAGTATGTGGACATATCACCATTACCATTTCTCAACACAGATCTTGAAGTACATGGGACTTACCCACCAGTTGTTG
AGGCATTTAGGCAGAAGATCCTTGCTGCGGATAGCATACTCTTCGCTTCACCTGAGTACAACTACTCTTTTACTGGGCCTCTGAAA
AATGCAATTGATTGGGCTTCCAGGCCCCCAAATGTATGGGCTGACAAAGCCGCTGCAATTGTGAGCACGGGAGGAGGTTTTGGCGG
CAGCCGATCTCAATATCATCTCCGCCAGACCGGGGTTTACCTTGATCTTCATTTCATCAATAAGCCCGAGTTTTTTCTAAACGCAT
TTCAACCTACTGCAAAATTTGATGGCGACGGCAACTTGATAGATGAGGCAGCCAAGTCGAAATTGAAAGATGTTCTCTTATCCTTG
TACGCATTCACGCTACGACTTCAAGGTAAGTGTGCATAGTTTATCCCACGATCCTTGTTGTGCATCTTAAAGATAATATTATTCCA
AAAATATTCATCTTATTGTACCCGCCCGCGAGTACTTAACCTTTAATTGCTCAAGTTCAAGGTATGTAATGACATTGTTCTTGTAT
CATGTGTTGTTTTTGCATGAAATCTATATGCTAATGCAGTTCCTGTAGAAATTCTACGGGACATCCGGAACTCCTATATATTAAAA
ATCATATGTAGATTTTCT
>ShNQR1 Striga hermonthica Sh.c10873_g1_i1.9809
AAACAAGTGTCCATGGCGGCGATCTCAGCTCCTACTCCAATCATCAAAGTCGCCGCCATCTGCGGTTCCCTCCGCAAAGGTTCCTA
TAATCGCGGCCTCCTCCGTGCCGCTATGGATATATCGAAGTCGATTGAAGGGTTGGAGATTGAGTATGTGGACATATCACCATTAC
CGTTTCTGAACACGGATCTTGAGGTAAATGGGACATACCCACCTGCTGTAGAGGCATTTAGACAGAAGATTCGTGCTGCTGATAGC
ATACTCTTTGCGTCGCCCGAGTACAACTACTCTATCACTGGACCTCTGAAAAACGCGATCGACTGGGCTTCCAGGGCACCAAACGT
CTGGGACGACAAAGCTGCTGCAATCGTGAGTGCGGGCGGTGGTTTTGGCGGTGGACGGTCACAGTATATTCTCCGCCAGACGGGGG
TTTACATCAACCTTCATTTCATCAATAAGCCGGAGTTCTTCCTGAACGCATTCCAAACTCCCTCTCCATTCGATAGTGATGGCAAC
TTGATCGATGATGCCTCCAAGACGAGGTTGAGAGCCGTTCTCTTGTCCTTGCAGGCATTCACAATCCGGCTCCAGGCTTAGTGTGT
GCAGGTATACCTCCCAGTAGGTGAGGTAATATTGTTCAAGTATATTCATTATCAGTATGTGTGTAATGGTACTACTCTTGTATTGT
GTTTGTGTGTTTTAATGGGATTCATGTACTTTGTCTAAAATTTATAGCGATGTATGAATAAGGTTCATCATAAGATTTGATGAACC
GTTTTTCTTGTTAGACATCTTGAATCAGCTGCAGTGCTGTCTTTCAACTACAGACTATCAATAAACGTCGATCTGTGGTCTATTGC
TTGTGGCTTGTGCAATATAAATATTTACTGGTGTGCTAACTGTTGAGCAGTGGTGCATCTAATAGCTCGGTTGACACTTGACAGTG
TTATAAATGTGGTCTCAGTCTTGATTGATAACCAAATAAGACTCGGGTTTTAGTAAAGGGAAAAAAAAAAAAAA
>TvNQR1 Triphysaria versicolor Tv.c5241_g2_i3.1001
CTAGCAACGTATTGATCACAAATAATGACAGGGGTGATTAAGATTGCTTGTCTCTCTGGTTCTCTACGAAAAGGCTCTTTTCACAC
TGGCCTCATTCGTCATGCTATTCATCTGAACATATTTGGTGATGTTGATGTTGATGATATGGAATTCGTGTACGTTGATATTTCGA
ATTTGCCATTGTTGAACACTGATCTTGAGGAAGAAGGGACTTTTCCTCTTGAAGTTGAAGATTTTCGTGATTATATTCTTGGAGCT
GATAGCTTTCTATTTGCTTCTCCTGAGTATAACTATTCCATCTCAGCTCCACTAAAGAATGCACTTGATTGGGGATCAAGACCACC
AAATGTATGGGCTGGTAAAACTGCTGCCATAGTGAGTGTTGGAGGAGGACACGGCGGTGCAAAGTCACACTATCATCTCCGACAAA
TTGGAGTTTTCATCGATCTTCATTTCATCAATAAACCCGAGTTTTTCTTAAACGCGTTTCAGCCTCCAGCAAAATTTAACGGTGAT
GGCGATTTGATTGATCAAGATGCCAAGAATAATTTGGAGGGAGTTCTTTTGTCCTTGAAGGCATTTACCCTTCAACTTCAAGGCAA
AAATTGAAACAACTGAGGTCCATAAGTCTTTAATTTGAAATAAAATATGTTTTGAATTTCAC

>PaFRO1 Phelipanche aegyptiaca Pa.c170304_g1_i1.19665
TTGTAGCAAATCAACATGTACTATGAAGTCATGTTGTATATATACAGCACATAATTTTCCATAGGAACTACAATATATCTCTTTCA
ACAAAGCAACAGAGAAAAAGAAGATCATGGGAAACTATAGAGTCTTAAAGATTATATCCCTTTTGGTGTTCTTTGGATGGCTATTG
CTTTGGACATTGGTGCCAACAAAAGTTTATAAAAATAAATGGACTCCCAAACTAAAAAAAAGTCTCAGCTCAACATACTTCAGAGA
ACAAGGTGTAAACATCCTTCTATTCTCATTCCCGATAATGTTCATAGCAGCTTTCAGCTGTGTTTATCTCCATCTCCAGACCAAGA
AACCGGTAAACGGCTCAAAGAAGAGCGCCGCACGAGCTGAGAAGCGTCACTGTGCATCGTTGAGGCGTCCAATGTTTGTGAACGCG
CCTCTTGGGATCGTCACCGCCGTGGAGGTTCTATTTGCGATAATGTTCGTTGCTCTCCTAGCCTGGTCTCTAGGAAATTACTTGTA
CATCAGCTTCGGCGCTCTTCACATGCACAAAGTCGGAGAGAAAGTATGGCAAGCAAAGTTTAGAAGCGTGTCGTTGAGGCTAGGTT
ACGTCGGAAACACTTGTTGGGCGTTTCTCTTCTTTCCCGTAACAAGAGGGTCTTCGATTTTGCCTCTCCTTGGATTGACATCTGAA
TCGAGCATTAAGTACCACATCTGGCTCGGCCACGCCTCGAATCTCCTCTTCGTGTTGCACACCGTAGGGTTCTTCATCTACTGGGC
GATGACCGATCAAATGTCTGACGCCCTGGAATGGAGCAGCACGTACGTGTCGAACGTGGCGGGAGAGATTGCGACGGTGATAGCGG
TCGTGATGTGGGCCACGAGCCTCGAAAGGGTGCGGAGGAAGATGTTCGAGCTCTTCTTCTACACTCATTACCTCTACACGGTTTAC
ATTGTCTTCTACGTTTTGCACGTCGGACCCGCCTACACGTGTATGATTCTCCCCGGAATATTCCTCTTCCTCATAGACCGTTACTT

GCGCTTCTTGCAATCCCGAGACCATGCAAGATTGCTCGCCGCTCGCCTTTTGCCCTGCGGTGCCCTCGAACTCAACTTCTCCAAGT
CCCGAGGGCTGCAATATAGTCCGACAAGCATAATGTTCATAAATGTGCCATCTATCTCCAAGCTACAATGGCACGCGTTCACGGTG
ACGTCGAACTGCAATTCAGAGCCGGAGCAGTTGAGCGTTGTGATCAAAAGCGTGGGAAGTTGGTCTCAACGCCTTTACAAACAACT
CTCTTCTTCCCCTGAGCACTTGGCTGTTTCCGTTGAAGGACCCTATGGACCGTCTTCTTTCTCGTTTCTAAGTCATGAGTCTCTGA
TAATGGTAAGCGGTGGGAGCGGAATAACGCCGTTCATCTCAGTCATTCGCGAGATTCTATTCCAAAGCACAGATCCCAAAGCTCAC
ATCCCAAAGATTCGCCTCATTAGTGCGTTTAAAAACTCCTCCGATCTCTCAATGTTGGACTTCTTGGTCCCCATCTCCGGTGCATT
CTCCGACCAGATCCCCCAAATAGACCTCCAAATTGAGGCTTACGTCACCCAGGACAACGAAGGGGACAAAGTAGAAGCCCTTCAAA
CCATATGCTTCAAGCCCAATCCGCGCGACTCCCCAGTAACGGCGACTCTCGGCCCAGACAACTGGCTCTGGCTCGGCGCAATAATA
GCGTCGTCGTTTTTGCTGTTCCTTGTCGTTTTGGGGCTCGTCACGCGGTACTACATTTACCCGATCGACATGAGGGACGCAAAGCG
TCCTTACCACTACTCATATCGGTGTCTCTGGGACATGTTTTTGGCGTGCACGAGTGTTTGCGTGGTCTCTAGCGTGGCGTTTCTTT
TGTGCAGGAGAAGAAACGAGGCCAAAGGAAACATTAAGAACCAAATCACCAACGTCGAAATGACGACTCCAACGGCTACGCCGGCC
TCGTGGTCATACGCGGCGGGGCAGAACCTTGACCTTGAGAGCCTTCCTAATCAGTCATTAGCTCAAGCCATGAAGGTTCATTTTGG
CGCCAGGCCTGATCTCAAGAAGATTTTGTTCGAGTCGGAAGCGTCTGACGTGGGAGTTTTGGTTTGTGGCCGAGGAAGATGCGAC
ACGAGGTCGCCAAGATTTGTGCCTCCGGTTCGGCAGATAATCTGCATTTTGAGTCTATTAGCTTTAATCTCTGATTAATCATTTCC
CTTAATTTCTACACAGATCAATCAATATGTAAATGTCGTTTGTATGTATTTCTATCAACAAGCCTGTTTTTTCATTTTTCAACTTG
TAACAAGTTTGTGATCTATGTGTAATAAATAAGTGTATATATTAAAGATCGGAAGAGCGTCGT
>ShNOX1 Striga hermonthica Sh.c18212_g1_i2.3625
AATTCTTCTTGCAATTTCTGAGCCAAGGACACAATTATTAGGTAAGAGAATTATTCCCCATAACTGATTTAACCAACTTCTCTTCA
TTCTTCCTCTCTCTTTCACCTTCCAAAGATCTCTCTTATACTGGTAAAAAATGAGAGAAAACCTCATTGACATGGGATCATCCAAC
ATCGGCTCGGATCTTCCGAGGGACGAATCCGTAAAATGCACTCTCGAGCGTATAGAAGTCGACACCATGGCAAACGACTCGGACGG
GAAAACGGGCCCACGCATTGGAAGGCTCGAATCGGGCGTGGACCGAGGCCTCAGAAGCCTCCGTTTTCTCGACCGAACCACAACCG
GAAAAGAGGAGGACTCGTGGAGAGCCATAGAGAAGCGTTTTCATCAGCTTTCGGTTGAAGGCAGGCTGTTCAAGGACAAATTCGGA
GTCTGCATTGGGTTGGGGGATAGTAAGGAGTTTGCAGAGGAGCTTTACGACGCGCTGGCGAGGCGGAGGAACGTGAGCACGGAAAA
CGGGATAACCGTGAACGATGTTCGGGAGTTTTGGGAGGATATGACAAATCAGGACCTAGACACAAGGCTTCATATTTTCTTTGACA
TGTGTGACAAAAATGGTGATGGGAGGCTATCAGAGGATGAGGTTAAAGAGGTTCTGGTAATGAGCGCCTCGGCGAACAAGCTCTCG
AATTTCAAGCAGCAAGCCGCCACTTATGCTTCGTTAATCATGGAGGAGCTCGACCCGGACCACCAAGGATATATCGAGATGTGGCA
ACTAGAAGCTCTATTGAGAGGAATGGTCGGTTCCGAGCAAGGAAAAATGAAAGCTACCCGAAAAACCGAGACACTAGCGAAGACTA
TGATTCCCAAGGAATACCGAACGCCCGTCAGCAAATTCGTGTCCAAAAATACCGAGCGATTCTTCGAAAACTGGATGAAGATATGG
GTCTTGTGCCTTTGGTCCTTCATAAACATATCCCTCTTCATCTGGAAATTCAATCAGTACAAAAAACGAGCCGCGTTTCGAGTCAT
GGGCTACTGCCTCTGCTCGGCAAAAGCCTCGGCCGAGACCGTCAAATTCAACATGGCACTCATTCTCCTGCCCGTCTGCAGAAGGA
CGCTGACGTGGCTCCGCGAGTCTTTTCTCGGGACTTTTATACCCTTCGACGAGAACATTAATTTCCACAAAATCATTGCTGCTGGG
ATTGTGTTGGGTACATTGATCCACGTCGTAATGCACTGCACCTGCAACTTCGTGAGGCTTGTTTCGTGTCCCGATAATCAGTTTTA
CTCGATTTTTGGGCCAGCGTTCGATTTTCATCGGCCGAGTTATTTAGATCTCGCAGGGACCACGGTTGGGGTCACGGGGATAGTGA
TGCTCGTTTTGATGGTTTTTTCGTTCACGCTTGCGACTCATTCGTTTCGGAGGAATGTGGTGAAGCTGAGGTGGCCATTTTGCCAC
CTGGCGGGATTTAACTCGTTCTGGTACGCGCACCATTTGTTGGCTCTTGTTTATGTGCTCTTGATCGTTCATGGTTACTTCATATT
CATCACCCGAGAATGGTACAAGAAGACGACGTGGATGTACGTGGCAGTGCCGATGCTGGTCTATACGAGCGAGAGAATACTTACAC
TTTACGATCACAACTATAAAGTTGGGATCATCAAGGCTGTGATCTATACGGGTAATGTGCTAGCATTGTACATGAGTAGGCCTCCG
GGATTCAAATACAAGAGTGGGATGTACCTTTTCGTCAAGTGCCCGGATATATCGAACTTCGAATGGCACCCGTTTTCAATCACGTC
TGCACCGGACGATGACTACCTTAGCGTACACATACGAACACTCGGAGACTGGACAACTGAACTCAAGAACCGTTTTGCAACGGCAT
GCGAACCTCGATCGATCAAACCGAGAAGAGGTAATCTAGTGAGAATGGAAACAAAGGCATATTCCGAAACTCTCCAAGACTTCCCG
CGAATCGTGATAAAGGGTCCATACGGAGCTCCGGCACAAAACTACAAGAAGTTCGATATCCTTCTGCTCATCGGTTTGGGTATTGG
AGCCACCCCTTTTATTAGCATTATAAAGGATATAGTGAATAACGAGTCGAGATATGTACTAACAGATGATGCCTCTGGAGGAGATA
AGAAAGGTCCCGAGCGAGCGTATTTCTACTGGGTAACAAGAGAACAAGGATCGTTCGACTGGTTCAAAGGCGTTATGGACGACATA
GCAGAATACGATCACAATCATATAATAGAAATGCACAACTATTTGACGAGCATGTACGAGGAAGGAGATGCTCGGTCAGCACTAAT
AACAATGGTTCAATCGTTACAACACGCTAAAAATGGAGTCGATGTCTTATCGGAAAGCAGGATACGGACACATTTTGCTCGGCCGA
ACTGGAGGAAAGTATTCACTCACTTAACTAGCGTGCATCCTTCTACCCGAATAGGCGTATTCTACTGTGGGAGTCCTACGCTAACC
AAGCCACTCAAGAAGCTCTGTCAGGAGTTCAGCTTGAATTCGTCTACACGATTTCAGTTTCACAAAGAGAACTTCTAAAGCTTGTT
GCGATGCTTTTTGGTCGCTTATTATCAGCATCGGCTTTTGATTGTTCGATTTTTCGAACCTTCGACTAGCATAGAGAAATTAGGAA
GGTAATGAATGAATCTGGAAAGTGACCGAAAGATTTACGGAACTTCCTATATTTGTTAAAAGCTTCAAAGTTCGATTAACTGCTAG
ATCACAAGTAATTTTCTATAACTCCCTTAAAAGAGTG
>TvFRO1 Triphysaria versicolor Tv.c101806_g1_i2.15882
AAAAAACAAAGTGAAAAAGATGAAATATCGTGCAGCATGGCAGCTTTTTTTCATGTTGGTCTTTGTAGGATATATGCTTATATGGA
TTATGTTGCCTAATAAGACTTACAAGGATTCATGGACTCCTAAACTTAACAAACATCTCAACTCCACATACTTTCGCGAACAAGGG
ACGAATCTTGTTTTATTTAGTTTCCCGGTTATGCTCATTGCCTCTTTAGGATGCGTTTATCTACATTTCCAGCAAAAGTCGAATAA
TTCTAAGAGTAATAAGAGCTACTTACAATCATTGAAAACCCCGGCAATGGTAATGGCCCCACTCGGAATAGTCAACGCCATCGAGC
TTACATTCGCAGCCATGTTTATCGCTCTTTTAATCTGGTCATTAGCAAATTACTTGTATGTCAGCTTTGGCCACCTCCACATGCAT
ACGCCCGGAGAAAAAGTGTGGCACGCGAAGTTTCGCAGTGTGTCCTTAAGGCTTGGATACATCGGGAACACATGTTGGGCTTTTCT
GTTTTTCCCCGTGACACGTGGATCGTCGATCTTGCCCCTGATGGGCCTCACATCTGAGTCCAGCATCAAATATCACATTTGGCTTG
GCCATCTTTCGATGGGCCTTTTCGCCCTCCATAGTGTTGGCTTCGTCATCTATTGGTGGTCGATGACTCATCAAATGTACCTGATG
TTGGAGTGGAGCAGTACGTATCTGTCGAACGTGGCTGGGGTGATCGCGTTCGGTCTATCGTTGGCTATATGGGGAACGAGCCTAAA
TCGGGTCAGGAGAAAAATGTTTGAGCTTTTCTATTACACACATCATATCTACATTCCCTTTGTTTTCTTCTACATGCTTCATGTTG
GAGTGGGCTATCTATGCATGATCCTTCCCGGGATTTTCCTCTTTCTCATCGACCGGTACTTGAGGTTCTTACAATCGAGACGGAGG
GCCCGGCTAGTTTCGGCTCGAATTTTACCTAACCGCACCATGGAACTTACATTCGCCAAGAATCCAGGGTTGTGTTATAGTACGAC

TAGTACCTTGTTTGTGCATGTGCCGGTCGTGTGTAAATTACAGTGGCATCCGTTTACAGTAACTTCGAGCTGTGATTTGGAGACGG
ATAAGTTAAGTGTTGTGATTAAAAGTCAAGGGAGTTGGACACAAAAGCTCTATAACCAACTTTCCTCTTCCCTCGTCTGTCTTCAA
GTGTCGACTGAAGGACCTTATGGACCGACATCGTCTAATTTTCTAAGTCGTGAGTCGTTAGTGATGATCAGTGGTGGAAGTGGGAT
CACGCCGTTCATTTCCATAATACGCGAAACCATTTCCGAAAGCACAAAATCAAAAACAAAGGTTCCGAAAATCCTTCTCATCGCAG
CTTTCAAGAACACGGCCGATCTAACAATGCTCGACCTCTTGCTTCCTATCTCAAACACAACAACTCCTTTAGACATTTCCAAGCTA
CACCTCCGAATAGAAGCTTACGTCACTCAAGAGCAAGAAAAACCTCTCGACGACACAAAAACTCAAATCGAGACAAAAATGTTCAA
GCCAAATCCGTCTGATTCGCCGATATCTTCAACTCTTGGTAAAAACAGCTGGCTATGGCTCGGAGGGATAATATCATCGTCTTTTG
TGATGTTTCTACTTCTTTTGGGATTTGTGACTCGTTACCACATATATCCTGTCGAAAGAAGAGGAGAGAATTATCATTACAGTTAT
AAGATTTTATGGGACATGTTTCTTGTGGTTGCTAGTGTGTTTGTGGCTACCAGTGCTGTCTTTTTGTGGCAAAAGAGAGAAATTTC
AAAGGAAGCTGGGAAGCAAATTCAGAATGTGGATATGCTGACACCAGCGATGATGTCACCATCGTCGTGGTTATGTGGTGTGGGTT
CAGATAGAGAACTCGAGAGCCTTCCGAGTCAGTCTCTTGTTCAGTCGACAAACGTGCATTTTGGCTCACGGCCTGATTTAAAGAGG
ATACTTTTGGAGTGCAAAGAATCGGACGTTGGAGTTCTAGTTAGCGGACCTAAAAGCATGCGACACGATGTTGCTAAAATATGTTC
GTCTGGTGACATGAAAAACCTGCATTTCGAGTCTATCAGCTTCAACTGGTGATGCATGTGACATTGTATACCTTTTGTTCGTGTTA
CGTATTTTCTCTTTTCGGATCCTTGCCACGTTTCAATCGATCTGCAAAGGAATGTTGATTTGTGCCTTGGAATTATTTAAATTGAG
TTTGTTACAGCTCTGCGTGCATAAGAATTTGTTTGATTTTATTTTCTGAGTTATGTTGGAATATATTTGAAGCTAATTAAATCTGT
TTGAATGTATCATATAACCATTTG

>PaCSD1 Phelipanche aegyptiaca Pa.c163109_g2_i2.16053
CATCGGCATCCACCAAGTCGATTTTGACTATTTACTACTAATCACCAAGCAGCTTCCAATGATCCGCAGCTACGGGGAGCCCGCGA
AGCTTGTCTAGCCTCCAAATACTGTTTAAGCAGAGATACCGATGACACCGCAGGCAGGGCGGGGGCCAGCGTTGCCGGTCTTCTTA
GACTCCTCGGTGCCACCCCTGCCGAGGTCATCAGTGCCGGCGTGGACGACGACGGTACGGCCGATGATGCTCTCAGCACCAATGAG
CTTGATCAGCTTGTCGGTGGTGGAGCCAACGGCGTTGCCCTGGCCATCGGTCTTGATGTTGCCGAGGTCACCAACGTGGCGGTCCT
CGTCGGTGGGAGCACCGTGGCCCTTTCCGTGAGGGTTGAAGTGAGGGCCAGCGGAGGTGCAGCCGTTGGTGTTGTCACCGAAGGCG
TGAATGTGCATGCCACGCTCAGCGTTGGCGTCGTTGCCGCTGATGTTCCATGAGATGCTGGTAGGAGCAGACTCGGATTCCTGCTC
GAAGGTGACGGTGCCCTTGACGTTGGAGTCACCACGGACGACTGCGACGGCCTTGACCATTTTGACGATTTTGATGCTTTCTTCGG
ACGTTTGATGTGAGAATTGAGAGCTTTTGCTGATGGAGAAGAGGAGAGAAGGTGGACGATGATCAAGCTCTCGCGTTTT
>ShCSD1 Striga hermonthica Sh.c12050_g1_i1.14196
ATCCACCAAGTCGATTTTGACTATTTACTACTAATCACCAAGCAGCTTCCAATGATCCGCAGCTACGGGGAGCCCGCGAAGCTTGT
CTAGCCTCCAAATACTGTTTAAGCAGAGATACCGATGACACCGCAGGCAGGGCGGGGGCCAGCGTTGCCGGTCTTCTTAGACTCCT
CGGTGCCACCCCTGCCGAGGTCATCAGTGCCGGCGTGGACGACGACGGTACGGCCGATGATGCTCTCAGCACCAATGAGCTTGATC
AGCTTGTCGGTGGTGGAGCCAACGGCGTTGCCCTGGCCATCGGTCTTGATGTTGCCGAGGTCACCAACGTGGCGGTCCTCGTCGGT
GGGAGCACCGTGGCCCTTTCCGTGAGGGTTGAAGTGAGGGCCAGCGGAGGTGCAGCCGTTGGTGTTGTCACCGAAGGCGTGAATGT
GCATGCCACGCTCAGCGTTGGCGTCGTTGCCGCTGATGTTCCATGAGATGCTGGTAGGAGCAGACTCGGATTCCTGCTCGAAGGTG
ACGGTGCCCTTGACGTTGGAGTCACCACGGACGACTGCGACGGCCTTGACCATTTTGACGATTTTGATGCTTTCTTCGGACGTTGA
TGTGCAG

>PaCDS2 Phelipanche aegyptiaca Pa.c24060_g1_i1.8178
GTTGAAGTTGAAAATATGTATATTGAGATCCGACAGGTCGAGCTTGTCTCGAGCTCGAGTTTGACAGAGTATGAGTCAAGCTCTGA
CCCGAGTTGCTCGACTCGTTTGCAGCCGTATATGCAATCCATGGTTCGGTCCTCTAAGATAACCTAAGCAGACCCTTTCAGCCAAT
ATTGCTATGTAATACTTAACCTACAACCAAACTTGCAGCCTGACGTGACGTCATTAAACCATAGCTCTGTAATTATTCAACTCTTA
GTGAAGTAAAACTAGTTCCGATGGTAATACCGTAATAAGCATAACACAAGCTTACAATGTTTTGAGAAGGTTATATCAACTCTGAT
AGAGAACTCATAGATAGTTCAATTATTGGTCGATAATGCCGAATAACAGTTATATAATACAACGTAAAATCACACCGATGCCATAG
GCCTCTATATGAACCATAAAGAGTTATGGAGTTCGGACGCACCAACGTGGTTTCGGAACAAGAAGCTACTAAATTACAGTGGAGTC
AATCCAACAACACCACATGCCAAACGCCCGCCAGCATTGCCTGTAGTAAGACTAAGCTCATGACCGCCCTTTCCGAGATCGTCTTC
ATGTTCATGAACAACAAAAGCTCTTCCAACAACTGAGTGAGGACCACTCAACGGTATCTGTGTGTCTACAATTGTCGCTTCAGCCA
CACCCTCAGCATTAGCAATTATGTTTCCCAGGTCACCCGCATGACGAACATCGTCTTCAGGAGCTCCATGTGTCAAGCCATTCGGG
TTAAAATGTGCTCCTGTTGATATACATCCATTTGTGGTGTCACCGTACTCATGAAGGTGAAACCCGTGTTTGCCTGGTGCGAGTCC
AGTCACACGAACACTCACAGTAGTGGGGCCATCGTCTTCTTGAGTTAGAGTGACGACGCCTTCGACGCTCGAAGTGCCTTTGAGAA
CGGCGACAGCTTTGTTGCTGGCGGCGACG
>ShCSD2 Striga hermonthica Sh.c17257_g1_i1.7008
TTTGCAAATGAAAATCTTTCAATTTTGAGTTTTTTAACTAGAGAAAGAAGAGTTGTTTCAACTGTACATGCTTTTTGCCATTTATT
GAAACAAAACCTAGTTCATGCAAATCGCTTAACCATAGTTAGTACCCTTTTTATTAAACTCACTGAAGCAAGTGCCATTACACAAT
TTTCTGAACCACAAGGTTGTAGATTCAAACTGAAGGGTTACATCACTCAATTACTAATAGACAAAGCCAAAATAACAGTAAATACA
ACACAATGTAAAAACATAGTTCCACCATATATGGGCCTCTCTCTCTATATGAACCGCACAGAATCAGGCCCCAAGGGTGGGTTTTG
AACCATCTTACTGTGGGGTCAAACCAACAACACCACAAGCCAAACGTCCGCCAGCATTGCCTGTAGTTAGACTAAGCTCATGACCG
CCTTTTCCAAGGTCATCCTCAAGTTCATGAACAACAAAAGCTCGTCCAACAACTGAGTTGGGCCCACTCAACGGTATCTGTGTGTC
GACAATTGTTGTCTCAGCTACACCTTCGCCATTAGCAACAATGTTTCCCAGGTCACCCGCATGACGGACCTCATCTTCAGGAGCTC
CATGAGTCAATCCATTTGGATTAAAATGTGCTCCTGTTGATATGCATCCATTTGTAGTGTCACCATATTCATGAAGGTGAAAGCCG
TGTTTGCCTGGAGTAAGTCCGGTTATACGAACATTCACAGTAGTTGGTCCATTGTCCTCCTGGGTTAGAGTAACGACGCCCTCGAC
GGTGGAGGTGCCTTTGAGAACAGCAACGGCCTTCTTGGTGGCGGCGACGACGGTGAGCGGCCTGGGAGCGGTGGCGGCTGAGAGAG
ACAAGAAAGGGCGGACTTTGGCCTTTACAGCGACGCCGTTGAAGGAGGAATGGAACGGAAGGGACTGCGACGAAACAGAGGACGTC
GCGGGGAAGAGAGCAGGAAGCTGAGGTGTTTGAACGGCGGCGGAGAGGACTGAGTGGGCGGCCATGGCTGCAAGCGCCGCTTGCAT

CTAAAAGATTGGATTTTGGGGTGGAGAAGTGATGAGATTTGAGTGAGTGGGTGAAGGGTTTACCGAGTCATGTGAGGGCATAACGA
AGGATATGGTATATATTTGTGC
>TvCSD2 Triphysaria versicolor Tv.c6286_g3_i1.7001
TTTTTTTTTTTTTTTACGGTTTGTCACGTCACTGGGCATGAAGCTAGTTTGGTTAATCGGTAAAGAATTCAATATCACGAAAAGTA
CACTTCCCTGGCACATTTCTCATAACATTTTTTAACAAATTTATTCAACTCCAATTGAAGCAAAACTAGTTCCCAATGGTAATGGT
ACAATACAGAACTAACCAGAGAGAGAAATCATTCATACATTGGTAGACAATTCGAAATACCAGTTATCTAATGCAATGCAAGTCAC
TGCGCCACAATAGGCCTCTATACGAGCCATCGAGAGTTACAGCGCACACACGAGGCTGGTTTCAAGCATCTTAAAGTGGCGTCAAA
CCAACAACACCACAGGCTAAACGCCCGCCAGCATTTCCAGTACTAAGACTAAGCTCATGACCACCCTTTCCAAGATCATCCTCAAG
TTCATGAACAACAAGAGCTCTTCCAACAACAGAGTTAGGACCACTTAAGGGTATATTTGTGTCCACGAGTGTTGCTTCAGCCACAC
CCTCAGCGTTAGCAACTATGTTTCCCAGATCACCCGCATGACGGACTTCGTCCTCAGGAGCTCCATGTGTTAAGCCATTTGGATTA
AAATGTGGTCCTGTTGATATACATCCATTTGTGGTGTCACCATACTCGTGCAGGTGAAACCCGTGTTTGCCTGGAGTAAGTCCAGT
TATACGAACAGTCACAGTAGTTGGGCCATCGTTCTCCTGGGTGAGAGTGACAACGCCCTCTACGCTCGAACTGCCCTTGAGAACGG
AGACGGCTTTCTTGGTGGCTGCGGAGACAGTGAGAGGCTTTGGGGCGGTGGTGGCGATGGAGAGAGACAAGAAGGGGCGAACATTG
GCCTTAAGAGCGATGCCGTGGAAGGATGAATGAAGTGAGAGAGACGGCGGAGAAATGGAGGATGATACGTTCGGGAAGAGAACAGC
AAAGTGAGCATTTGAGACAGTGGCGGCGGAGAAAACTGTGTGGGCGGCCATGGCTGCTAGCGTCGCTTGCATCAGAATGGATTGGA
TTGGATTAGTTGATAATGTGAGCGAGTGAGTTGGTATAATTCAAGCGAGTTTTTTTTTTACCTTTGTTATAAAGGCACAAATATAG
GTACGTGTTAG

>PaGPX4 Phelipanche aegyptiaca Pa.c20866_g2_i1.5606
TTTTTTTTTTTTTTTTTTTTAAAACGAAAAGGTAATTTGTTTCATAAGTTATATATTTATTTTGTTTGTTACAATATCCCTAAACC
AAAATTACACACAGACACATTACAATACAATTACAATTACACAAGCATTCACCAACTGAAAATTGAGTAAAAAGAAGACACAAACA
CACACCTAGTTTTGCAGAATTCTATAGGTCTTCAAGTTTCACCCAAAGCTTCCTTTATGTCACCCTCAATTGCCGATGGTGAGGTA
GACGGTCCATAACGTTTGATCACATGCCCATCTTTGTCGACTAAAAATTTGGTGAAGTTCCATTTTATGCCGGAACCAAAATAACC
ACCTTTGCTTGCTTTAAGGAACTTATAAACCGGTGCTGCATCTGCTCCATTCACTTTTACCTTTTGAAATATGGGATACTCAGCTT
TGAATCTTGTACAAGCAAAATGTTCTGCTTCTTGGCTCGTTCCTGGCTCTTGATACAAGAATTGATTGCATGGGAAGGCCAATATC
TCAAAACCTTTATCCTTGTACTTAGAGTAAAGCTCAGTCATCTGCGTGTAATTTGAGTTCGTCAACCCACATTTAGACGCAACATT
AACAACAAGCAAGACTTTGCCTTTATAGATATTCAGATTCGCATCATTACCTTTAAAGTCCTTGACGTTGAAATCGTGGATCGATT
TTTCCGTGACCGATGAAGAAGCACCCATTGTGTTGTTTCTCTGTTTAAGCTCTCCTGTTTTTCCTTCCTACGCTTGACAAATACTG
CACTGATGCATCAATAATTGCA
>ShGPX4 Striga hermonthica Sh.c2340_g1_i1.4308
AAAAAACCCATTCCAAAGGGTTACCATCATTATAAACAACCAGATTTTCATGCCATATCACAAAACAATGCCATAAGAACCATTAA
TACAAGCACTCTATTAATGGAGGGAAATGGGTGAAAAAGAATACACACATCTAAGTATTTTTTAACTTTTTTTGCTGCTGTCTACA
GTTTTGCAATACCATAAAATTTTAAGGTCATAATATAACAACAAAACCGAATGCTCGCCTTTCAATCTTCACCGAGAGCTTTTTTT
ATATCACCCTCGATTGCCAAGGGTGGGGTAGAAGTCCCATAACGTTTGATCACACGACCATCTTTGTCAACTAGAAATTTGGTGAA
GTTCCATTTTATGCGAGACCCGCAAAAACCACCTTTACTTGCCTTAAGGAACTTATACACAGGTGCTGTATTTGGCCCATTCACTC
TTATCTTTTGAAAGATGGGATACTCAGCTTTGAATCTTGTGCAAGCAAATTGTTCTGCTTCTTGGCTTGAGCCTGGCTCTTGATAC
AAGAACTGATTGCACGGGAAGGCCAATATCTCAAAACCTTTATCCTTGTACTTGGAATAAAGCTCAGACATCTGTGTATAATTTGA
GTTCGTGAACCCACATTTGGACGCGACATTAACAACAAGCAAGACCTTGCCTTTATAAACACTCAGATTCACATCATTGCCTTTAC
TGTCCTTGACGTTGAAATCGTGGATCGAATTTTGTGGGGGGACTGAAACTGAAGAGGAAGAAGCACCCATTATTATGCTATCTGTT
TTATCTGCTTCTCTACGCTTGACCAGATAATCACAGGCCACATAAAGGCATTCATTCATTCACTTGTTATTGCAAAATGCGGAGGT
TTGAAAATTTTTTGTTGATGCTAAATGTGAAAACAGAGGAGAAAAGCCCCTTTATCACATGACAGGCTGGCACG
>TvGPX4 Triphysaria versicolor Tv.c4008_g1_i1.95
CCATAACGTTTAATGACTTGCCCCTTAAAACCCCAACATTTTTCAACCCTTTCACAACTACCCCACAACAAGAAAATGGGCGCTTC
TCCATCGGTCCCAGAAAAATCCATACACGAGTTTACTGTAAAGGACAGTAAAGGTAAAGATGTGGATTTGAACGCCTACAAAGGGA
AGGTCTTGCTTGTTGTTAACGTTGCTTCCAAATGTGGGTTTACAAATTCAAATTACACCCAGTTGACTGAGCTTCATTCTCAATAC
AAGGATAAGGGTTTTGAGATATTGGCATTCCCGTGTAACCAGTTCATGAATCAAGAACCGGGAACGAGCGAAGAAGCCGAGCAATT
TGCCTGCACAAGGTTCAAGGCTGAGTATCCCATCTTCCAAAAGGTGAGAGTGAATGGAGCAGAAGCAGCACCTGTATATAAGTTCC
TAAAATCGAGTAAAGGAGGTGGTTTCTTTGGCTCAAGCATAAAATGGAACTTTACTAAGTTTCTAGTCGACAAAGATGGGCAAGTC
ATTAAACGTTATGGAACAGCCACTTCGCCATCTTCAATCACGGATGATATCAAGAAAGCTCTGGGAGAAATTTGAAGGAGTTGTGT
TAATTTGATTTGGATGTTAGATACAAAGCTTTCTTCATTGCTCGTTTGTGCATTTGTTTTTCTTATTTTATCAAATTGTTTGAACC
AGAAG
>TvGPX5 Triphysaria versicolor Tv.c106689_g1_i1.19515
GAATAACACTAAAAAATCATAGTTTTGCATGTAAAAAAAGAAGAATTAAGAGAAACCCTTTTTCTTATGCCCACAACATTTTATTT
CGAAATAATATATCCTCCTATATCCAGTCCTTGAAGACAAATCATCCAATAAAAAACTATAGAATATACAATACAGCCATATGTTT
GTATGTCATATCCAAAACATCAAACTTACCATTTTACAATACTTGCTTTAATTATATGGATCAATTAATTAGCTTCGTGTTTATAC
TTCACCCAGGGCTCTTTTTATATCACCCTCAATTGCCAGCGGCGGGGTAGAAGGTGCATAACGTTTAATTACAAGCCCATTTTCAT
CAATTAAAAACTTGGTGAAGTTCCATTTTATGCGGGACCCAAAACAACCACCTTTACTTGCTTGAAGAAACTTATAAACTGGTGCT
GCATGCGAGCCGTTCACTTTTACCTTTTGAAAGATGGGATACTCAGCTTTGAATCTTGTACAAGCAAATTCCTCTGCTTCTTGGCT
CGACCCTGGCTCTTGATACAAGAATTGATTGCACGGGAAGGCCAATATCTCGAAACCTTTATCCCTGTACTTGGAATAAAGCTCAC
TCATCTGCGTATAATTTACGTTTGTGAACCCACATTTGGACGCAACATTAACAACAAGCAACACCTTGCCTTTATAAATCTCCAGG
TTCACATCATTACCTTTACTATCCCTGACGTTGAAATCGTGGATCGATTTCTGCAGGATTGAAGAAGAAGCGCCCATTTTCTTCTC
TGTTTGGCCTTCCTTGCGCCTGGTAAAACGGCGTCGTACGCACGTGAAAGCTTGATTAAGAATAATTATTGCAGGCAAGGAAATAA

ACGGCGTGCTACGATGTGAAAATAGAGGAAGGGTGGTGATGGTGAGTGGTATGGATGAGTGAATAATCAATCATCTGAGAATTGTT
GGATATTCTAAAAAAAATCAAAT

>PaGPX6.1 Phelipanche aegyptiaca Pa.c182494_g1_i3.24732
ATTTATTTCATATATTAAATTAAAATGACAGATTGCTTGCAGGATTTGAATCCATCTAATACGAAATGCTCATCAAGTAACTAGAT
GATGATTGATTGCTACATTGATATACCTATATAAATTGATTAGGTTGTCTATTAATTCACAGTTCGGCATTCCCCACATGATGGAA
TCACTGAGATAATATCAAAATATTTGCATACACTACTAAAGAAAAGAGAGAACTGCATTGTAAAACATCAGTAATATCCCGGAAAA
GATCTTACACAAACTCCGTACACTTTATTCACTCAAATATTCTCCACGGAGATCGTTATTAGTAGTAGTAGTAGTATGTATCGAGT
TACATACAGTCATGTGTACTCCTTACTCACGCTGTTCTTTGATAACCTTTCTTCGTTATTTTCTTTATATGTTTGCACCACTCTCC
AGGAGTTTCTTTATATCCTTCTCTATGCTGAGAGGAGAGGTAGTGGGCGCATATCGATCAACAACATGCCCCTTTTGATCAACAAG
GAATTTGGAGAAGTTCCATTTAATGTTATCCCCCAAAAACCCTCCTTTAACTGATTTCAGGTATCTGTATAGTGGAGCAGCATTTG
GACCATTCACCTCAATCTTGTCAAATATGGGATACTCTGCCTTGAAACGAGTGCAAGCAAAGTTTTGGATTTGTTCATTGGTACCA
GGCTCTTGTGAACCAAACTGATTGCAGGGAAATGCAAGAATCTCAAAACCTTGATCCTTGTACTTCTCGTATAACTTGGTGAGCTC
GGTGTAATTTGAGTTGGTCAAGCCACACTGTGATGCAACATTGACAATTAGAAGGACTTTTCCCTTATATGTACTCAGATCAACAT
CATTCCTCTGGGCATCCTTGATAGTGAAATCATGGACAGATTGAGTCTGGGTTGACTGGCTAGCCATCGTATGATCCGCTCTGAAG
CCTAAAAACAAAGACTTTCTCGAACTAATACCAGATGAAGAGTACAAAGACCTAGCTTTGATTGGCTGAAAAAGCAAACAATTCGA
CGCAACAGCTGGTGAATTAAATTGGATTCTCGCCAAGATCGCATAAATGGAGCCC
>PaGPX6.2 Phelipanche aegyptiaca Pa.c21890_g1_i5.5938
CATAAAATTTCATAAAATCTTCCATTTTTCATTTTTTCTTCTAAGTGAGGGGGCATTGTGGTCATTTTGGATAAAAATCAAGTTTT
AACGCCATGAAGCATAAAATCCAAAATTTCCCAAATATAGACTACTTCAAAACCACTGAACCAGACACATTTTCATTTCAACACCA
TGACATCAAATTCTCAGTCATTTTCACATAGTTTCACCTTACACAAATCCAGAATCCTCGTTCACTTGCATATTCTCCAAAGTGAT
AATATTATTAGTAGTAGTAGTATTTATTGAGTCGTACCATACAATCACGTATACACCCTTTCCATAATTCATTGATCACCTTTCTC
CATCATTTTCTTTATATGTTTGCACCATTCTCCAGGAGTTTCTTTATATCCTTCTCTATGCTGAGAGGAGAGGTAGTGGGCGCATA
CCGATCAACAACATGACCCTCTCGATCAACAAGGAATTTGGAAAAGTTCCATTTAATTTTATCCCCCAAAAACCCTCCTTTGACTG
ATTTCAGGTATTTGTAGAGTGGAGCAGCATTTGGACCATTCACCTCAATCTTGTCAAATATGGGATACTCTGCCTTGAAACGAGTG
CAAGCAAAGTCTTGGATTTGTTCATTGGTACCAGGCTCTTGTGAACCAAACTGATTGCAGGGAAATGCAAGAATCTCAAACCCTTG
ATCCTTGTACTTCTCGTATAATTTGGTCAGCTCAGTGTAATTTGAGTTGGTCAGGCCACACTGTGATGCAACATTGACAATTAGAA
GGACTTTTCCCTTATATGTACTCAGATCAACATCATTCCTCTGGGCATCCTTAATAGTGAAGTCATGGACAGATTGAGTCTGGGTT
GACTGGCTAGCCATCGTATGATCCGCTCTGAAGCTCAAAAACAAAGACTTTCTCGAACTAATACCAGATGAAGAGTACAAAAACCT
AGCTTTGATTGGCTGAAAAAGCAAACAATTCGATGCAACAGCTGGTGAATTAAAATGGGCGCCTAAGCCTATGGGTCTGATCAAAT
TCCTGTTTCTCACAAGGCAAGTTGAGCAAAAGGACATGAGGAGATATATATAAAGGAATATAAAAAAGAATTACGGGTTCCAAATG
GATTTGGATGTGAAAGCGCATACGATACATGCAAGAGAGATTGGCAATGGTATGATTTGCGTGGCTGGACATGCTAATTTACGCGT
GACATAGTGGAATGTACACCACGTGGGATGCCTTCCAACTATATTTCGATACTCCACCA
>ShGPX6.1 Striga hermonthica Sh.c13821_g1_i1.4424
TTATCCAACCTTTCAGTTTTATCCGAACTTTTGCGCAAGCGAACCAATTTCGATCCACTGCATAACTATCCAACCTTTCAGTTTTA
TCAGCCGATCGAAGATAATAACAATATGGCCAGCCAGAGCCAGTCCAGCCCGGCCCAATCAATCCATGAATTCACTGTAAAGGATG
CCAAGGGCAGTGATGTCAATCTGGGTACTTATAAGGGCAAAGTCCTGCTGATTGTCAATGTTGCTTCACAATGTGGTCTGACCAAT
TCAAATTACACTGAGCTGACACAATTATACGAGAAGTACAAAGATCAAGGTCTAGAGATACTGGCATTTCCCTGCAATCAGTTTGG
CTCACAAGAGCCTGGCACCAATGAAGAAATTCAGGAATTCGTGTGCACTCGGTTCAAGGCTGAATATCCTGTATTTGACAAGGTCG
AGGTGAATGGTTCGAATGCTGATCCAATATACAAGTACATGAAGTCGATCAAAAGCAGCATTTTTGGGGACAGTATCAAATGGAAC
TTCTCCAAATTCCTTGTGGACAAAGAGGGCCGAGTTGTCGACCGTTATGCTCCCACCACATCTCCTCTTAGCATTGAGAAGGATAT
AAAGAAACTCCTTGAGAAGGCGTGAGCGTTGGAGGTAGCCAAGATCAATATGTTTCTGTTGGTGGTTAATATGAATAGTCGTGTTA
ATAAGAATTTGACTTGATGTACGCCTTACGACTTTGTTTGAAATAAATTGCCATCATACTTAAACAACTTGGAATAATTAATTTTG
CTGCTGTTATTTGGATAGCTGATCAGATCATCTTCTACCAGTGACATATATAGTACCATCTGTGATTATATAAGCAGAACTACTTT
AATAGCCGTTCTTGTGGATCCCAGTTTTTACTTCGTTCGTTG
>ShGPX6.2 Striga hermonthica Sh.c13847_g1_i2.4434
CAAAAGTCAAAACTTCCAGAAACTAGAAGTATAAAGTAAATAAAATCGTCTATACGTTGAAACTGATTGGCCACATTGTCAACACG
TGGCGTGCATCCAGCCTCTTACGCGTACATTAGCAACGTCACATATCTTGCGCCATCGTTCAAATCTGCATTCACATTTGGACCCA
TAATAAAGAATAGTAGTACTCTTTATTTCTTTTCTTTATATATATCTGTTCATTCATGTCCCTTTTCAAATCTCGCATTCTGATAA
ATATAAGAACCTCATACATCACAGGCTTAATTTCTTCTTCCCCGATCAGAACACATCTCGATTCACCAGTTACCCCGCCGGAATCT
GCGCGTTTTGAGCCCATTAAAGCTAGGTTTCTGTACGGCCCTTCATCCTGTGTTTCTTCGAGGAAGTCTTTGATTTCGAGCTTCAG
AGCAGATCGTACGATGGCCAGCCAATCAGATTCGCCTCAATCTGTGCATGGTTTCACTGTTAAGGATGCCAAGGGGAATGATGTCG
ATTTGAGTACATATAAGGGAAAAGTCCTTCTAATTGTCAATGTCGCATCACAATGTGGCCTGACCAATTCAAATTACACCGAGTTG
ACTAAGTTGTATGAGAAGTACAAAGATCAAGGTTTGGAGATATTGGCATTTCCATGCAACCAGTTTGGCTCACAAGAGCCCGGTAC
AAACGAACAAATTCAAGAGTTTGCTTGTACTCGTTTCAAGGCTGAGTACCCCATATTCGATAAGATTGAGGTAAATGGTTCAAATG
CTGCTCCACTATACAAATATATGAAATCAGTCAAAGGTGGGCTCTTCGGGGACAGCATCAAATGGAACTTCTCTAAATTTCTTGTT
GACCAAGAGGGTCATGTTGTTGACCGATATGCTCCGACTACTTCCCCTCTCAGCATAGAGAAGGACGTCAAGAAATTGCTGGAGAA
AGGTGCAAAGAACTGAAGAAGTTCATGGAGTAAAGGTTAATAAGCGATGCAGTGTATGAATAACTCAATAACGATAATAACTTCTT
GGAGCATATGAGGAGCTTTTTACATTTGTATTATGAATGCTTTGTATTGCAAGGTTTCGACTGCATACATTTGTGTTGTGTGTTAT
GAATGCTTTATATTGCAAAGGTTTCGATCATTACTAAAGGATCGGAATTTGGTGATGCCATAGTTTGTGCCTTTAGTGCTTATTTG
TCATGTAGGGTTTATCTTCAAAATCTTGCTTAACATTTTATAACAGCTAGAAAAACCAAACTTTTGACTT
>TvGPX6 Triphysaria versicolor Tv.c6882_g2_i2.4631

CGCGTTACAACTTAGTCCATTAGTTCTAACAAAAAAAACTTGTGATCAAAAGCTTACTACTGTCTCCCCCATTTCAATACACTGCG
TAGTTATCCAACTTCCTTGTTTTCTCGATCGATCAAAGCTAGATTTGAATATAAGATGGCCAGCCAGTCCAGCACGCCCCAATCAA
TCCATGATTTCACTGTCAAGGATGCTAAGGGTAATGATGTAAATCTGGGTATCTACAAGGGCAAAGTCTTGTTGATTGTCAATGTT
GCTTCACAGTGTGGCTTAACCAATTCAAATTACACTGAGTTGACCCAACTATATGACAAATACAAGGGCCAAGGATTGGAGATTTT
GGCATTTCCTTGCAATCAGTTTGGCTCACAAGAGCCCGGCTCCAATGAAGAAATTCAGGAATTTGTCTGCACTCGTTTTAAGGCCG
AGTATCCCGTATTTGACAAGGTTGATGTAAATGGGCCGAATAGTGCTCCGATATACAAGTACATGAAGTCGGCCAAAGGCAGCATT
TTCGGAGACGGTATTAAATGGAACTTCTCGAAGTTCCTTGTCGATAAAGAGGGTCGTGTTGTTGATCGCTATGCTCCCACCACATC
TCCTCTAAGCATCGAGAAGGATATTAAGAAGCTCCTTCAGAAGGCTTAAGAATGGGTATTAATAAGCTCCTATACCGTTTGGTCCA
TAGCTATAAAAGAATAAACCCTCGTTGAGGATGTTTGTATTAATAATCGTGTGAAACGTACAAGACCTTGTTATTTTGCATCGTGT
TCGAAGAATATAGGTCATACTTCCGTGGGAATTTATCGTATTTATTGTTTGTTTCCCATGCTCGATCTTTATGAAAACTATGGTTT
TATATTTGTTCTTTAAATATCCCGGAATGTCTGATTTTCTTGTTTAGTAATTTTTTCAGTAATTGAGAAAACTGTGTTTCGTGTTA
CAAAGTGGGGCATATAATTTAACTAAAAGTAAATAAAACTACAAACAAG

>ShGPX8 Striga hermonthica Sh.c14473_g1_i1.4564
GTAGAGATGGGTATGAGCCTTGTTGCGTCTGGTGGGTCCCATGGCGGTATCGAGACCGTGAAGAAGAAGAGGGGTAGGCCGAGGAA
GTACGGGCCCGAAGGGCCTCCGCCTAAGGTGTCTTTGAAACTCTCGTCGCCTGTTCCAAAAACGCCTTCGTCGTCTGATCCGAACA
CTTCGGGGGAGAAAGCAAGACGAGGGAGGCCGCCCGGGACCGGGTGGAAACAGAAACTTGCTCCTCTTGGCGATTGGATGAATAGC
TCAGCTGGACAGGCTTTTACCCCCCATGTTCTTCACATAGGAACTGGAGAGGATATTTCAGCAAAAATATTGGCATTTGCGCAACA
GAGATCGAGAGCTTTATGCATAATGTCAGGAAGTGGATCGGTTTCTGTAGTGACACTAAAGCAACCTGCAAATCCTTCTAGCACCG
TAACTTTTGAGGGTCGATTTGAGATTCTATGCCTGTCTGGTTCTTACTTGGTAGCTGAGAATGGTGGTCCCCACACTCGAACTGGT
GGTATTAGCATATCTGTGTGTAATCCTGATGGCCATATGCTTGGGGGTTCAATAGGCGGCCGACTTATTGCAGCAAATAACGTGCA
GGTGGTGGGGTGCAGCTTTGTGTACGATGCTACAAAGGCAAAGACCAAACCCGAGTCCCAAATAACGAACGAAAGGAACCTTCTAG
AACAATCTTCTGAAGAGGCGTACACCCCGTATAGTGCTGCCGCCTCTAACCAAAACCCTAATGTGGGGCCGAGTGCTTGGCAGCTA
AGTTCACGACAAGACATAAAGAGATCACAAGAGGATATTGACCTGACTCGCGGATGATTTATGATGATAAAATGTGCTATTATGAA
TTCTCTGTATATATGATGGAAAACGAGAAAATGTTCGAGTGTTCGAGTTGTGCTAACGTATTCTTGTAACGAAATTCAGTCTGTAG
GATCTTAAGAAAGTAGGAGAAAGTAGTAGAGTTTGATATAGTCGGTTGAAGTTTAAGCAAGGATGTATTGTTCTAAATTTCTAATA
GGTGTAAGTGATGTTGTATTAATGGATTGTATTGTAAAAAACATTGTGGGATTTATTCATTTTCAGTTGGAAACACTAGTTGTGCA
TAGCTTGACCATATATTGTTAACTTGAAAAATTTTAGACAGAGGGTAGAAATTCTTCATAACTAGAATTGAAAATTAGGGGAACTG
AGTTCAAAATTATTACAAGAAATCCAACACAATTATAATTACAACAAGTCTCAATCTGCTAGCCATTTAATATATCATGCAAGATA
ACCGACCAGTCGAGCATCAATAGGACCCGAGGTTGCAAGGGTTATGAAAAAAACACGTTTTTTCGTTTACGACCCCAATAGCTTTA
CGATATCCCTCTCGATCGTGAGAGGTGAGGTCGTAGGGTAATAACGATCAACAGGCTGCCCATTCTTATCTACCAAAAACTTTGCA
AAATTCCACTGGACATCATCCCCAAAGATTCCCCATTTGCCCGATTTTAAGAACTTGTACAGTGGAGCAGCATTATCACCATTCAC
CTCAATCTTAGCAAAGATGGGGAAATCCGACTTGAATCGAGTGCAAACAAAGTCGAGAATTTGATCATTGCTTCCAGGTTCTTCTT
CACCAAACTGGTTGCATGGGAAAGCTAAAATCTCGAGGCCTTGATCTCTGAACTTTTCATACAGCTTATTTAGCTCGGTATAGTTT
GAATTTGTCATCCCACATTTTGAAGCAACATTGACTATCAACAACACTTTGCCTCTGTAAGTGCTTAGCTCCACATCACCTCCATT
AGCATCCTTGACAGTGAAATCGAATACAGACTGAGGCGACGCCATTTGTTTGAATTCGTGTTTTCGGAGAGCTAAAGAAGAATGAG
ATAGTGGTAGTGTATCTCGTGAGGGCCATAGCTAAGTACATAG
>TvGPX8 Triphysaria versicolor Tv.c6531_g1_i1.4491
CATTCGTAAAATGATCATCCAGTAACGAACATAAAATGGTAAATATCAAAGAACATTGAGTTCAAATACATTTAAAATACATTTTA
AAAGAAAAGTGGAGTATCTCATGTGTTCCCCGATATTATTTCATTTTTTTACAACATAAGGAATCAGATTTTCTATTCCAACTCGA
ATCAATCAAAGTCCAATGGTTATCGCAGTGACAAAATAGAATAAAAACTGCTGCCCATGGAAATAAATTCGACATGTTATATAACA
ATAGTACAATAAAGGTTCAACAGTACCGGTTATGAGAGCTCCAGGAGCTTTCGAATATCCCTCTCAATTGTAAGAGGCGATGTCGT
GGGGTAATAGCGATCGACAGGCTGACCATTCTTATCAACCAAAAACTTGGCAAAATTCCACTGGATGTCATCGCCAAAGATTCCCC
ACTTTCCCGACTTCAAGAACTTGTACAGTGGAGCAGCATTATCACCATTCACTTCGATCTTACCGAAGATGGGAAAATCTGACTTG
AATCGAGCGCAAACAAAATCAAGAATTTGATCATTGTTTCCAGGTTCTTCTTCGCCAAACTGATTGCATGGAAAAGCCAGAATCTC
GAGGCCTTGATCTTTGTACTTTTCATACAGGTGATTCAGCTCAGTGTAGTTCGAGTTAGTCATCCCACATTTGGAGGCAACATTGA
CAATGAGCAACAGTTTGCCTCTATAAATGCTTAGATCCACATCATCTCCTTTAGCATCCTTGACGGCGACATCAAACACTGATTGC
GGCGATGCCATTTGATTAAAGTGAGTCCG

**Supplementary Data Set 2.** Lists of transcripts used in coexpression network analysis. Transcript ID, functional annotations, expression levels, network connectivity and PhQ connectivity are provided for each species.
